# Models of the cardiac L-type calcium current: a quantitative review

**DOI:** 10.1101/2021.10.04.462988

**Authors:** Aditi Agrawal, Ken Wang, Liudmila Polonchuk, Jonathan Cooper, Maurice Hendrix, David J. Gavaghan, Gary R. Mirams, Michael Clerx

## Abstract

The L-type calcium current (I_CaL_) plays a critical role in cardiac electrophysiology, and models of I_CaL_ are vital tools to predict arrhythmogenicity of drugs and mutations. Five decades of measuring and modelling I_CaL_ have resulted in several competing theories (encoded in mathematical equations). However, the introduction of new models has not typically been accompanied by a data-driven critical comparison with previous work, so that it is unclear which model is best suited for any particular application. In this review, we describe and compare 73 published mammalian I_CaL_ models, and use simulated experiments to show that there is a large variability in their predictions, which is not substantially diminished when grouping by species or other categories. We provide model code for 60 models, list major data sources, and discuss experimental and modelling work that will be required to reduce this huge list of competing theories and ultimately develop a community consensus model of I_CaL_.

## 1 Introduction

The ‘long-lasting’ or L-type calcium current (I_CaL_) is an ionic current found in cardiomyocytes, neurons, endocrine cells, and other cell types throughout the body (Striessnig et al., 2014). It plays a crucial part in critical functions such as hormone secretion, regulation of gene expression, and contraction of cardiac and smooth muscle (Hofmann et al., 2014). Its critical role in cardiac cellular electrophysiology has led to extensive modelling efforts, and a large number of models of cardiac I_CaL_ electrophysiology have been published since the 1970s. Incorporated into models of the cardiac action potential (AP), I_CaL_ models have a long and successful history of application in fundamental electrophysiology research (Noble and Rudy, 2001). More recently, they have been used or proposed for use in drug safety assessment (Mirams et al., 2012) and risk stratification in cohorts with ion channel mutations (Hoefen et al., 2012). Choosing an I_CaL_ model for such safety-critical applications is not an easy task, as several models can be found in the literature, which vary widely in their structure and assumptions, and are often published in a paper form which does not lend itself to quantitative comparison without considerable effort (Cooper et al., 2011). On a more fundamental level, each model of I_CaL_ represents a testable theory of its physiology, and the existence of so many competing theories reveals both gaps in our knowledge and a need for their comparison and synthesis. In this review, we take the first major steps to facilitate a critical reassessment by the community, by collecting I_CaL_ models, analysing their qualitative differences, and providing quantitative comparisons based on simulations with freely reusable code. We start with a brief description of I_CaL_ physiology and the common structure of its models.

### 1.1 Biophysical properties of I_CaL_

The L-type calcium channels (LCCs) through which I_CaL_ flows consist of a pore-forming *α*_1_ subunit and the auxiliary subunits *β* and *α_2_δ* (Dolphin, 2016). The most common *α*_1_ subunit in Purkinje cells and ventricular and atrial cardiomyocytes is Ca_V_1.2 (encoded by the gene *CACNA1C*), while Ca_V_1.3 (*CACNA1D*) is more prevalent in the sinoatrial node (SAN) and atrio-ventricular node (AVN) (Gaborit et al., 2007; Zamponi et al., 2015).

Electrically, I_CaL_ is characterised by fast, membrane voltage (*V_m_*)-dependent activation at relatively depolarised potentials (around 0mV) and slower inactivation that depends on *V_m_* (voltage-dependent inactivation, VDI) and, via a separate process (Hadley and Lederer, 1991), on the intracellular calcium concentration (calcium-dependent inactivation, CDI). Although LCCs are highly selective and I_CaL_ is predominantly carried by calcium ions, smaller potassium and sodium components have also been measured (Hess et al., 1986) and are commonly included in models.

An influential study by Hess et al. (1984) on currents through isolated LCCs introduced the idea that they have three different modes of gating: A ‘mode 0’ in which channel openings are rare, a ‘mode 1’ characterised by rapid bursts of brief openings, and finally a ‘mode 2’ (rarely observed without first introducing a channel agonist) that features long channel openings and only brief periods of channel closure. These *long* openings, together with their *large* unitary conductance lead to the ‘L-type’ designation (Nilius et al., 1985). Although in this review we focus on whole-cell ‘aggregate’ I_CaL_, many of the models we will discuss are inspired by concepts from single-channel work.

In ventricular and atrial cardiomyocytes, I_CaL_ is the major inward current responsible for maintaining the ‘plateau’ of the AP (Catterall, 2011), making it a major determinant of the AP duration (Figure 1) and important cell properties such as restitution and refractoriness. The calcium influx during the plateau phase initiates the process of calcium-induced-calcium release (CICR): calcium flowing in through LCCs causes the nearby ryanodine receptor (RyR) channels to open, leading to a further, and much larger influx of calcium from the sarcoplasmic reticulum (SR, see Figure 3), the major calcium store inside the cell. The resulting increase of calcium concentration in the cytosol allows formation of cross-bridges between myosin heads and actin filaments, and leads to contraction (Winslow et al., 2016; Eisner et al., 2017).

**Figure 1:**
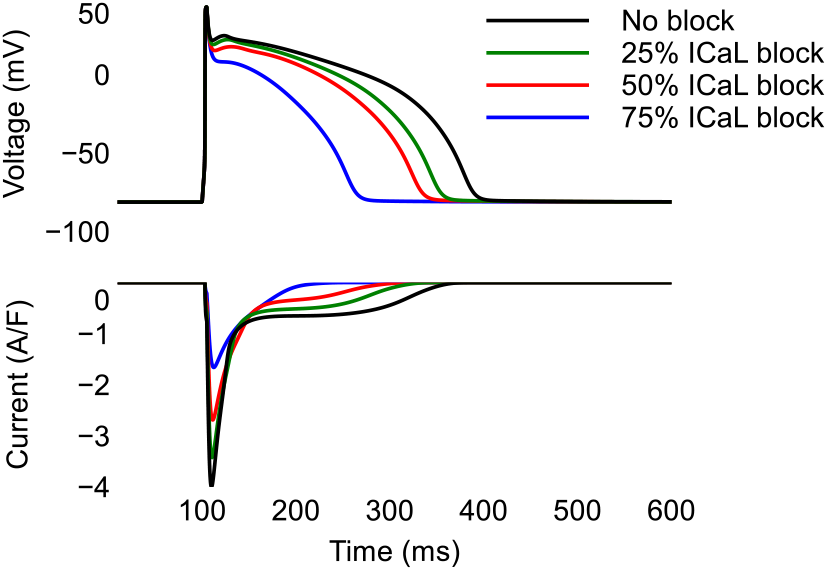
Blocking the L-type calcium current (I_CaL_) shortens the AP duration. **Top**: AP and **Bottom**: I_CaL_ during 1Hz simulation of the AP model by Grandi et al. (2010).

The crucial role of calcium channels in maintaining healthy cardiac function is borne out by clinical evidence. For example, reduction of the expression of the Ca_V_1.2 *α*_1_ subunit by less than half can lead to heart failure (Goonasekera et al., 2012), and mutations in *CACNA1C* and in its subunits have been linked to Brugada syndrome, Timothy syndrome, arrhythmia, and structural heart defects (Hofmann et al., 2014). This same vital role makes LCCs an important target for pharmacological antiarrhythmic therapy. For example, nifedipine, verapamil, and diltiazem inhibit I_CaL_ (Ortner and Striessnig, 2016). Unintended pharmacological modulation of I_CaL_, conversely, can lead to drug-induced arrhythmias and other adverse cardiovascular effects. Initiatives like the comprehensive *in vitro* proarrhythmia assay (CiPA) therefore take I_CaL_ into account when attempting to predict pro-arrhythmic risk (Li et al., 2019).

Several of the body’s regulatory mechanisms target I_CaL_. Following sympathetic stimulation (‘fight-or-flight’ response), *β* 1-adrenergic membrane receptors are activated, which triggers intracellular processes resulting in the conversion of ATP to cyclic adenosine monophosphate (cAMP), which activates protein kinase A (PKA), which phosphorylates the LCCs. This increases I_CaL_ leading to a larger Ca^2+^ influx and a larger Ca^2+^ release through CICR, which ultimately increases contractile strength (Gao et al., 1997; van der Heyden et al., 2005). Parasympathetic stimulation (‘rest-and-digest’) can reduce cAMP levels, which is known to inhibit the effects of sympathetic stimulation on I_CaL_ but appears to have little effect on the basal current (McDonald et al., 1994). In addition to these mechanisms, Ca^2+^/calmodulin-dependent protein kinase II (CaMKII) is both regulated by [Ca^2+^], and a regulator of [Ca^2+^] via phosphorylation of both LCCs and RyRs (Maier and Bers, 2007; Winslow et al., 2016). This co-dependence between [Ca^2+^] and CaMKII activity can create a positive feedback mechanism in which peak I_CaL_ increases when successive voltage pulses are applied, in a process called Ca^2+^-dependent facilitation (CDF, Bers and Morotti, 2014) and sometimes equated with ‘mode 2’ channel gating. Unusually high CaMKII levels have been observed in conditions such as heart failure, and are thought to be arrhythmogenic (Soltis and Saucerman, 2010; Bers and Morotti, 2014).

Finally, LCCs are not spread uniformly throughout the cell membrane, but occur mostly (90%) in specialised regions called *dyads* (Scriven et al., 2000). In dyads, the outer (t-tubular) cell membrane is in close proximity to the SR membrane (< 15nm, see Fawcett and McNutt, 1969; Eisner et al., 2017) and LCCs are in very close contact with RyRs. This enables CICR (Stern, 1992) and allows fast signalling (Harvey and Hell, 2013; Abriel et al., 2015), but the small size of these ‘nanodomains’ also means that [Ca^2+^] can fluctuate much faster and with a much higher amplitude than in the rest of the cytosol. Models of I_CaL_ are often not created in isolation, but as part of a larger AP model that makes certain assumptions about LCC localisation and hence about the [Ca^2+^] affecting I_CaL_. As a result, the assumed channel localisation must be taken into account when comparing models of I_CaL_.

### 1.2 Models of cardiac I_CaL_

The first published models of a cardiac calcium current appeared in the 1970s (Bassingthwaighte and Reuter, 1972; McAllister et al., 1975), only a few years after the first calcium current recordings in cardiac preparations. Although Bassingthwaighte and Reuter (1972) confidently referred to a ‘calcium current whose role is primarily in the maintenance of the plateau [and in] regenerative depolarisation in low-sodium medium’, the exact nature of the current and the species it was carried by was controversial at the time, so that the more cautious names ‘secondary inward’ and ‘slow inward current’ were commonly used (Fozzard, 2002).

Although the complexity of I_CaL_ models has increased over time (keeping pace with new discoveries about the current and its role in cardiac function), all models that we assess in this manuscript can be expressed in a common form:

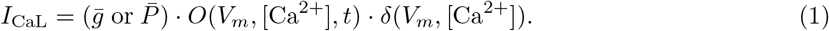

Here, *O* is the fraction of open channels (or the *open probability* of a single channel). The time-varying behaviour, i.e. the *kinetics* (or *gating*), of the channel(s) is captured by *O*, which is voltage- and time-dependent and may also depend on [Ca^2+^]. The term *δ* represents the ‘driving term’ accounting for the electrochemical forces driving the ionic movement through the open channels. *δ* can depend on both *V_m_* and the ionic concentrations on either side of the channel. The current is scaled by either a maximal conductance 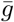 or a maximum permeability 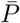, with different units depending on the choice of driving force model (see Section 2.5). Conductance and permeability are functions of the single channel permeability and the total number of channels in the membrane (Schulz et al., 2006), which can vary over longer periods (hours to days); however, they are treated as constants in all of the models we surveyed.

We shall discuss a variety of approaches taken to modelling *O* and *δ*, as well as factors not included in Equation 1 such as channel localisation and regulation by other variables. In general, we shall aim to take a reductionist approach and regard components affecting I_CaL_ (e.g. local calcium dynamics, CaMKII signalling or *β*-adrenergic stimulation) as separate entities. Examples of the full equations for early (McAllister et al., 1975) and more recent models (Faber and Rudy, 2000) can be viewed in the online repository accompanying this paper, at https://github.com/CardiacModelling/ical-model-review.

### 1.3 Outline

In this review we identify, discuss, and compare the models that have been used to describe cardiac mammalian I_CaL_ to date. In the next section we describe the process by which we have identified models to be included in the review, and classify the 73 models we identified by their origins, the assumptions made by their authors about LCC localisation and the variables that regulate I_CaL_, and the equations used for I_CaL_ gating and driving force. In section 3 we compare a subset of 60 models *quantitatively*, by simulating the application of several experimental protocols. These include voltage-clamp protocols for activation, inactivation, and recovery, as well as an AP-clamp protocol and a protocol in which AP and calcium transient (CaT) are both clamped. We then test whether we can reduce the observed variability in predictions by grouping models according to the qualitative aspects identified earlier. In the final section we discuss possible approaches for choosing an I_CaL_ model for a simulation study and developing and documenting new models of I_CaL_.

## 2 Qualitative comparison

To identify models of I_CaL_ we searched on PubMed (in March 2021) for publications containing the term “L-type calcium channel”. Because many I_CaL_ models are presented as part of a larger modelling effort, we also included the term “cardiac cell model”, and scanned lists of models such as those given in Noble et al. (2012) and Heijman (2012). To constrain our study, we only included I_CaL_ models that were presented by the authors as representing I_CaL_ from healthy human or mammalian cells. Similarly, we focused on basal (versus adrenergically stimulated) I_CaL_ models.

Older models of ‘calcium current’ and ‘slow’ or ‘second inward’ current were included if they modelled atrial, ventricular, or Purkinje cells, in which L-type is the dominant calcium current (Gaborit et al., 2007). The models included this way were by Bassingthwaighte and Reuter (1972), McAllister et al. (1975), Beeler and Reuter (1977), DiFrancesco and Noble (1985), Hilgemann and Noble (1987), and Noble et al. (1991). Some older models were excluded if they modelled the sino-atrial node, in which the dominant calcium current is of the T-type (Mesirca et al., 2014). Next, we selected modern models where the authors explicitly included an L-type calcium current.

Some studies included several model variants. The models by Pandit et al. (2001), Ten Tusscher et al. (2004), Ten Tusscher and Panfilov (2006), O’Hara et al. (2011), and Tomek et al. (2019) have epi-, endo-, or mid-myocardial variants but the I_CaL_ kinetics are the same. In simulations where a whole-cell model was used (Section 3.6), we used the epicardial variant of these models. Similarly, the apical cell model (representing cells at the apex/tip of the ventricles) was chosen for Bondarenko et al. (2004). The I_CaL_ kinetics varies among the different versions presented for Inada et al. (2009), Paci et al. (2013), and Varela et al. (2016), for which we chose the atrio-nodal (a transitional region between atrium and atrioventricular node), atrial-like, and right-atrial model respectively.

The next selection was based on the model equations. Models were excluded from our overview if they only changed the value of conductance or permeability parameter from a previous model; models were included if they introduced new equations, made changes to equations, or changed rate or driving force parameter values. Models for which the equations were not fully given in the publication or an online addendum were also excluded in this step (9 models in total), as were models for which the equations were suspected to contain typesetting errors (1 model). Due to its historic importance, an exception was made for Bassingthwaighte and Reuter (1972) for which the published equations did not match the accompanying figures. This model is included in our qualitative, but not quantitative, analysis.

Finally, studies that calculated the current by considering the sum of all individual LCCs stochastically were only included for qualitative analysis. These included the studies by Greenstein and Winslow (2002), Restrepo et al. (2008), Hashambhoy et al. (2009), and Nivala et al. (2012). The studies by Hinch et al. (2004), Greenstein et al. (2006), Asakura et al. (2014), and Himeno et al. (2015) described I_CaL_ using models in which the gating was linked to the ryanodine receptors, and were also included for qualitative analysis only.

Applying these criteria we found a total of 73 distinct I_CaL_ models, spanning five decades of research.

### 2.1 Model provenance

Models are not created in isolation, but inherit ideas, equations, and parameter values from previous models. Figure 2 presents a tentative ‘phylogeny’ of I_CaL_ models, showing how the models that are in use today were derived from one another over time. To establish these relationships, we searched the model publications for explicit references to parent models. In some cases (e.g. Rasmusson et al., 1990) the parent models did not meet the selection criteria used in this study, which is indicated in the figure by a dashed line. In other cases (e.g. Lindblad et al., 1996; Demir et al., 1999; Bondarenko et al., 2004) the parent model was not mentioned explicitly but could be inferred from the equations.

**Figure 2:**
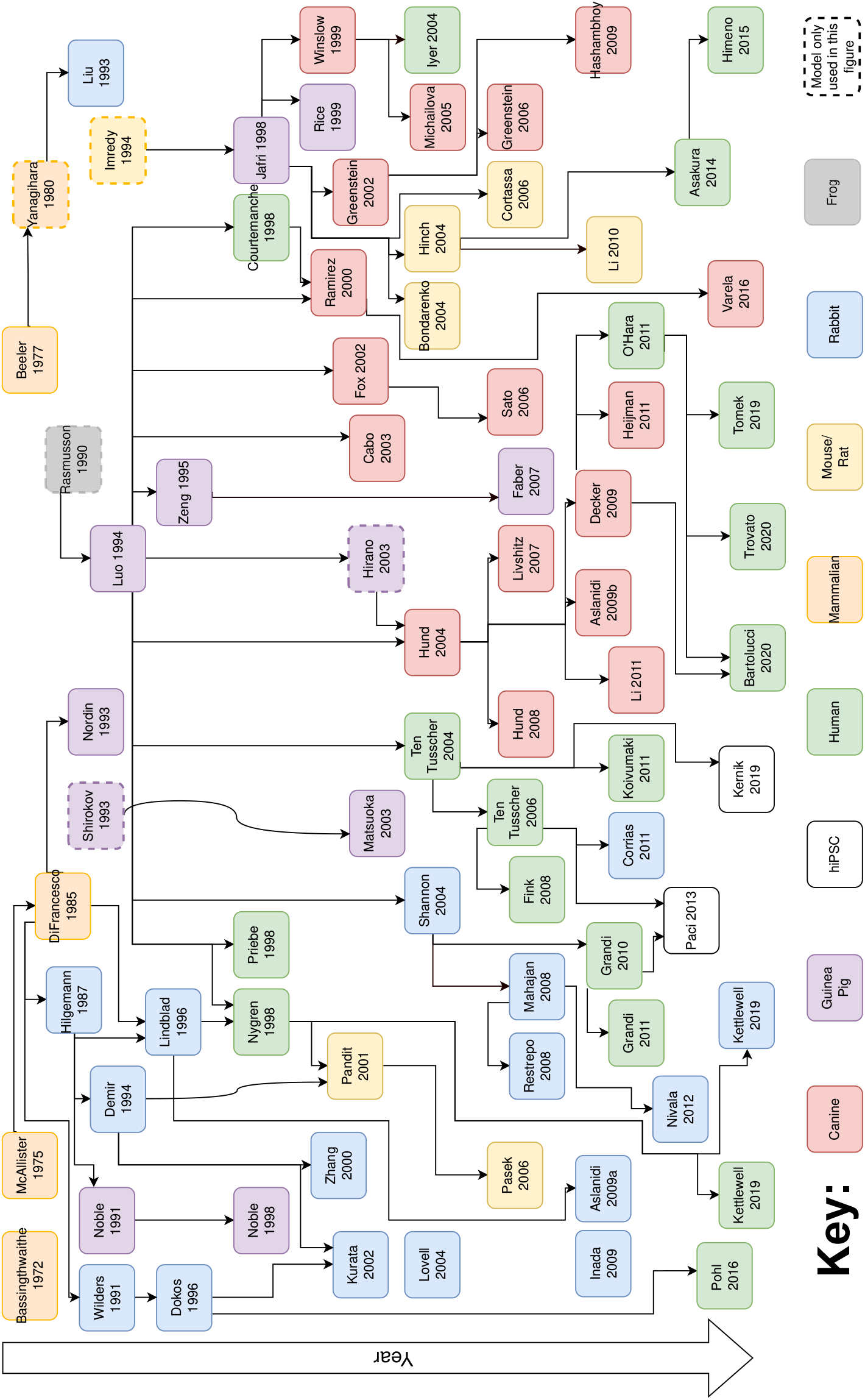
A schematic (and simplified) representation of human and mammalian I_CaL_ model development from 1972 to present. The key describes the species associated with each model.

**Figure 3:**
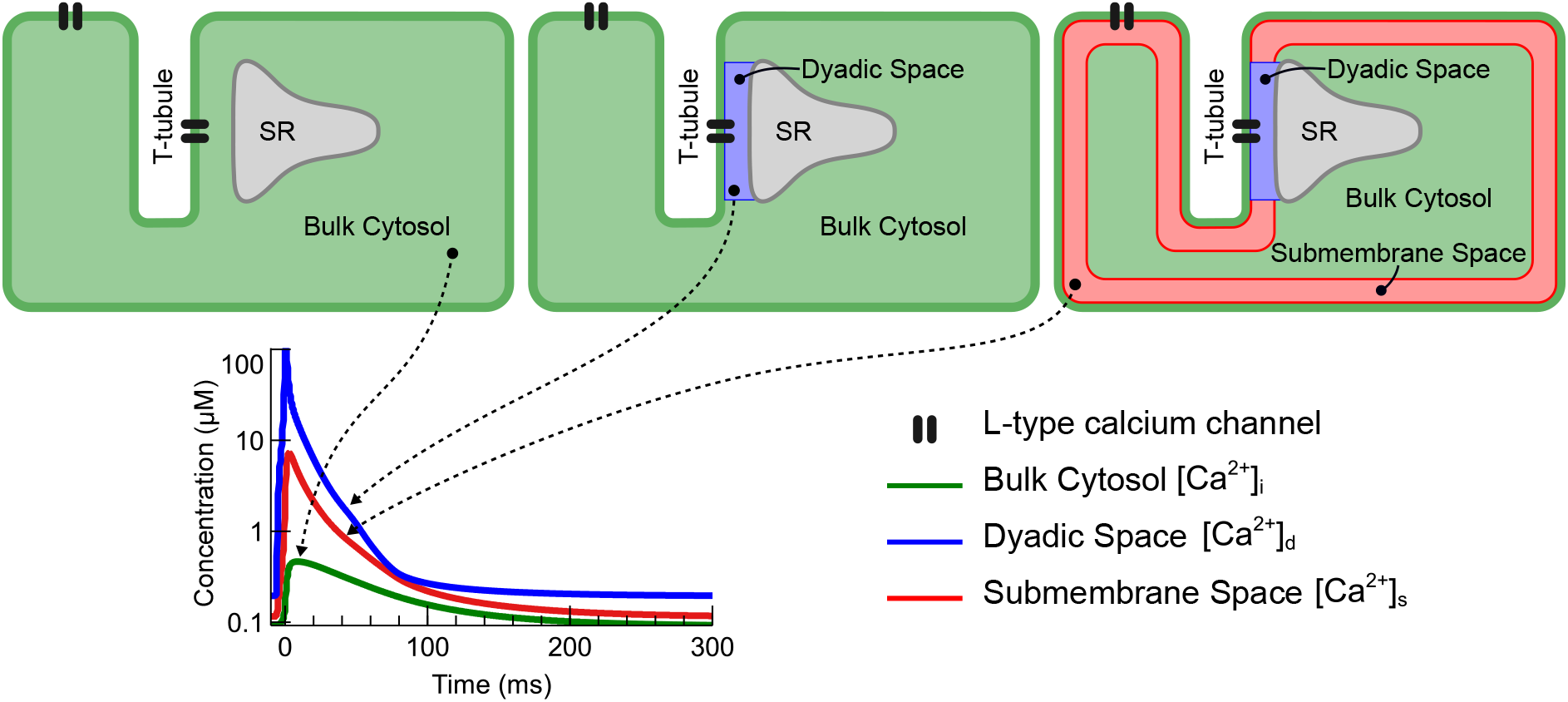
Schematic representations of a cardiomyocyte illustrating three common models of subcellular subspaces and LCC localisation. In small subspaces [Ca^2+^] can differ from the bulk cytosolic concentration by orders of magnitude, as illustrated in the lower panel which shows calcium transients from the Grandi et al. (2010) AP model stimulated at 1 Hz (note the logarithmic scale on the y-axis).

In constructing the figure, we attempted to limit each model to a single parent. This is a simplification, and the figure should not be read as a statement on originality. For the models up to around 1995 it is likely that each model ‘inherited’ in some way from all other models published up to that time.

The phylogeny shows that many models have encountered species switches over the course of their development. For example, Grandi et al. (2010) (human) inherits from Shannon 2004 (rabbit), Luo-Rudy 1994 (guinea pig), and Rasmusson 1990 (bullfrog). Similarly, the histories of the O’Hara et al. (2011) and Paci et al. (2013) models also contain several species switches. Note that this does not necessarily suggest re-use of data, as models are frequently re-parameterised to new data sets. However, inheritance of parameter values (and thereby indirectly of data sources) is also very common in electrophysiology models (Niederer et al., 2009).

### 2.2 Local [Ca^2+^] and spatial organisation

Several studies, both experimental and mathematical, have argued that I_CaL_, CICR, and I_CaL_-regulation cannot be understood without treating the Ca^2+^ concentration at the intracellular channel mouth as a separate entity from the ‘global’ or ‘bulk’ cytosolic concentration. Perhaps the earliest suggestion is found in Bassingthwaighte and Reuter (1972), who argued for the existence of a local subspace with elevated [Ca^2+^] to explain the observed reversal potential for I_CaL_, which was much lower than predicted by theory. An influential analysis of CICR by Stern (1992) reviewed the experimental evidence that the amount of Ca^2+^ released by the SR is proportional to the ‘trigger’ [Ca^2+^] with a very high amplification factor, and went on to show that such a *graded, high-amplification* response could not be achieved in models that considered only a single global [Ca^2+^]. A series of experimental studies investigating the mechanism behind (and calmodulin dependence of) CDI suggested that both a ‘local’ and a ‘global’ [Ca^2+^] were involved (Peterson et al., 1999; Tadross et al., 2008) (in this theory, the ‘local’ level refers to very brief spikes in [Ca^2+^] that occur at the channel mouth when an LCC is open, while the ‘global’ level is the lower [Ca^2+^] between spikes, which is dominated by diffusion from the cytosol). Finally, some models of I_CaL_ regulation have argued for highly localised presence of signalling molecules near LCC, in concentrations very different from the bulk cytosol (Harvey and Hell, 2013; Abriel et al., 2015).

Modellers of the cardiac AP have accounted for these local phenomena by dividing the cell into smaller-volume ‘subspaces’ in which calcium ions, carried in by I_CaL_ or released from the SR, can cause [Ca^2+^] to transiently rise to much higher levels than those seen in the bulk cytosolic space (Colman et al., 2022). Figure 3 shows three commonly used subspace configurations. In the simplest type of AP model, shown on the left, there are no subspaces and I_CaL_ flows directly into the bulk cytosol (e.g. DiFrancesco and Noble (1985)). The second type of cardiac model adds a ‘dyadic’ or ‘junctional’ space, sometimes called the ‘cleft’, between LCCs, located in the t-tubules, and the RyR in the adjacent SR. The illustration also shows LCCs connected directly to the bulk cytosol, as some cardiac models have a fraction (~10%) of I_CaL_ flowing directly into the cytosol (e.g. Tomek et al., 2019). The third type of subspace model retains the dyadic space, but adds a ‘submembrane’ or ‘subsarcolemmal’ space just below the remainder of the cell membrane, so that I_CaL_ flows either into the dyadic or the submembrane space. An example of this configuration, including compartment sizes and experimental references, is given in Shannon et al. (2004). Note that there are no hard boundaries between the subspaces: calcium can diffuse freely between them, but it is assumed that the rate of diffusion is slow enough for subspace [Ca^2+^] elevations to arise. In this review, we use [Ca^2+^]_*i*_ to denote the bulk cytosolic concentration, [Ca^2+^]_*s*_ for the submembrane space concentration, and [Ca^2+^]_*d*_ for the dyadic space concentration.

In Figure 3, and in the models it represents, a single t-tubule is used to represent a vast t-tubular network, and a single dyadic space represents all (thousands of) dyads. To allow further spatial heterogeneity, some studies (particularly those interested in calcium ‘sparks’ arising locally within a myocyte) have gone further and discretised the cell into a 3d network of ‘functional release units’, each with their own calcium concentration (see e.g. Rice et al., 1999; Nivala et al., 2012). Alternative, less computationally intense, approaches have been used by e.g. Nordin (1993), who define a ‘superficial, middle, and deep myoplasm’, and Koivumäki et al. (2011), who modelled the myocyte as a series of concentric shells.

These different models of cell geometry and [Ca^2+^] concentrations complicate the task of comparing I_CaL_ models in an AP model. Different assumptions about channel localisation in an AP model will lead to LCCs experiencing different intracellular calcium concentrations. I_CaL_ models subsequently have different assumptions about CDI and driving force. Furthermore, since I_CaL_ gating models are often calibrated in the context of an AP model, several more parameters in the I_CaL_ model will be sensitive to the chosen localisation.

To gain an overview of how the spatial model affects the model of I_CaL_ electrophysiology, we grouped the 73 I_CaL_ models by the calcium concentrations assumed to affect I_CaL_ gating (*O*) and driving force (*δ*), leading to the twelve categories shown in Table 1. Note that not all models shown in this table fit neatly into the three subspace configurations shown in Figure 3: the studies by Asakura et al. (2014) and Himeno et al. (2015) both define an extra intermediate subspace, between dyadic and bulk cytosolic; the model by Heijman et al. (2011) has a very small subspace *within the dyadic space*, into which only I_CaL_ flows; and the models by Noble et al. (1998), Mahajan et al. (2008b), and Restrepo et al. (2008) use different concentrations for the gating and driving force, which means they cannot be neatly assigned a position in Figure 3.

**Table 1:** I_CaL_ models grouped according to the [Ca^2+^] they are assumed to depend on in their open probability (*O*) and driving force (*δ*). For a graphical overview, see Figure 4.

| No. | Models | $O$ | $\delta$ |
| --- | --- | --- | --- |
| 1 | Bassingthwaighte and Reuter (1972), McAllister et al. (1975), Liu et al. (1993), Demir et al. (1994), Lindblad et al. (1996), Zhang et al. (2000), Aslanidi et al. (2009a), Inada et al. (2009), and Kettlewell et al. (2019) | - | - |
| 2 | Beeler and Reuter (1977), Noble et al. (1991), and Wilders et al. (1991) | - | $[Ca^{2+}]_i$ |
| 3 | Courtemanche et al. (1998), Ramirez et al. (2000), Lovell et al. (2004), and Varela et al. (2016) | $[Ca^{2+}]_i$ | - |
| 4 | Kurata et al. (2002) | $[Ca^{2+}]_s$ | - |
| 5 | Jafri et al. (1998), Rice et al. (1999), Nygren et al. (1998), Winslow et al. (1999), Pandit et al. (2001), Bondarenko et al. (2004), Iyer et al. (2004), Michailova et al. (2005), Cortassa et al. (2006), Pásek et al. (2006), and Koivumäki et al. (2011) | $[Ca^{2+}]_d$ | - |
| 6 | DiFrancesco and Noble (1985), Hilgemann and Noble (1987), Luo and Rudy (1994), Zeng et al. (1995), Dokos et al. (1996), Priebe and Beuckelmann (1998), Fox et al. (2002), Cabo and Boyden (2003), Matsuoka et al. (2003), Ten Tusscher et al. (2004), Paci et al. (2013), Pohl et al. (2016), and Kernik et al. (2019) | $[Ca^{2+}]_i$ | $[Ca^{2+}]_i$ |
| 7 | Nordin (1993), Sato et al. (2006), and Corrias et al. (2011) | $[Ca^{2+}]_s$ | $[Ca^{2+}]_s$ |
| 8 | Hinch et al. (2004), Hund and Rudy (2004), Greenstein and Winslow (2002), Greenstein et al. (2006), Ten Tusscher and Panfilov (2006), Faber et al. (2007), Livshitz and Rudy (2007), Fink et al. (2008), Hund et al. (2008), Aslanidi et al. (2009b), Decker et al. (2009), Hashambhoy et al. (2009), Li et al. (2010), Rovetti et al. (2010), Heijman et al. (2011), O'Hara et al. (2011), Li and Rudy (2011), Nivala et al. (2012), Bartolucci et al. (2020), and Trovato et al. (2020) | $[Ca^{2+}]_d$ | $[Ca^{2+}]_d$ |
| 9 | Tomek et al. (2019), Asakura et al. (2014), and Himeno et al. (2015) | $[Ca^{2+}]_i$,<br>$[Ca^{2+}]_d$ | $[Ca^{2+}]_i$,<br>$[Ca^{2+}]_d$ |
| 10 | Shannon et al. (2004), Grandi et al. (2010), and Grandi et al. (2011) | $[Ca^{2+}]_s$,<br>$[Ca^{2+}]_d$ | $[Ca^{2+}]_s$,<br>$[Ca^{2+}]_d$ |
| 11 | Noble et al. (1998) | $[Ca^{2+}]_d$ | $[Ca^{2+}]_i$ |
| 12 | Mahajan et al. (2008b) and Restrepo et al. (2008) | $[Ca^{2+}]_d$ | $[Ca^{2+}]_s$ |

Interestingly, only the six models in rows 9 and 10 of Table 1 show a dependency on more than one [Ca^2+^], and in all six this arises from the assumption that there are two distinct populations of LCCs, located in separate subspaces. The models in rows 11 and 12 assume that gating depends on the [Ca^2+^] in the dyadic nanodomain near the LCCs, but that the driving force is determined by the larger space surrounding it, which is the bulk cytosol in (Noble et al., 1998) and the subsarcolemmal space in Mahajan et al. (2008b) and Restrepo et al. (2008). None of the models we surveyed had the local and global components of CDI suggested by Tadross et al. (2008) in their O term.

The naming of spaces varies between studies, and sometimes overlaps or conflicts. In preparing Table 1 (and Figure 4), we classified a subspace as ‘dyadic’ if its primary role was in calcium handling (e.g. if it contained LCCs and connected to the SR) while we used the term ‘submembrane’ space only if several ion species flowed in and out of it, and it covered all or most of the (outer and t-tubular) membrane. An example where this conflicts is Sato et al. (2006), where we classified as ‘dyadic’ a subspace termed ‘submembrane’ by the authors.

**Figure 4:**
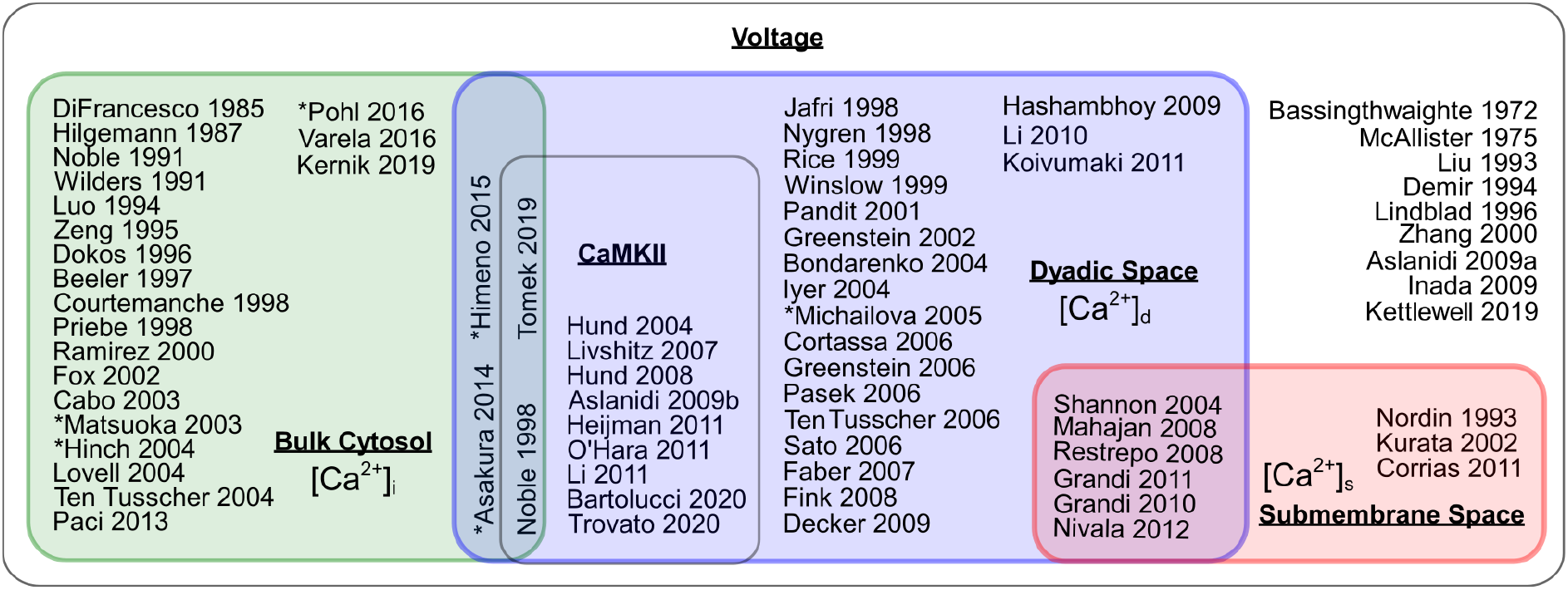
A Venn diagram showing the dependency of I_CaL_ gating and/or driving force on [Ca^2+^] and [CaMKII]. All models shown here depend on voltage. I_CaL_ models inside the green, blue, and red boxes depend on bulk cytosolic calcium, dyadic space calcium, and submembrane space calcium respectively, and models inside the black-edged central box also depend on the concentration of CaMKII. Models marked with an asterisk depend on additional variables (as described in the main text).

### 2.3 Regulating variables

In addition to the membrane potential (*V_m_*) and (local) intracellular [Ca^2+^], I_CaL_ models have been developed that are sensitive to several other variables. The most common of these is the CaMKII concentration, which is tracked in the model by Hund and Rudy (2004) and its descendants, and is used to estimate the fraction of CaMKII-phosphorylated channels in order to model CDF. The models by Matsuoka et al. (2003), Asakura et al. (2014), and Himeno et al. (2015) multiplied O by a term depending on the intracellular ATP concentration, to reproduce an effect observed by Noma and Shibasaki (1985), while the model by Michailova et al. (2005) based itself on experimental work by O’Rourke et al. (1992) showing that these effects were due to [MgATP] rather than the free ATP concentration. Parasympathetic (ACh) inhibition of I_CaL_ was included in the model by Pohl et al. (2016). Finally, the work by Hinch et al. (2004) uses a single model to describe the L-type calcium channels and ryanodine receptors, leading to a dependency on the state of the SR (interestingly, this model uses the [Ca^2+^]_*i*_ to estimate the concentration in the dyadic space).

Not many models surveyed in this review include effects of *β*-adrenergic stimulation on I_CaL_, but this is partly due to our selection criteria (see Section 2). An overview of studies focussing specifically on *β*-adrenergic stimulation (which almost always included effects on I_CaL_) is given in the supplement to Heijman et al. (2011).

An overview of the dependencies of each I_CaL_ model in this study is provided in Figure 4.

### 2.4 Gating mechanisms

The feature that varies most between I_CaL_ models is the set of equations used to describe the fraction of open channels, O. This part of the model determines how the current changes over time, i.e. it describes the current kinetics or gating. In this section we present a classification of I_CaL_ models into 15 distinct groups based on their gating equations.

I_CaL_ gating is commonly modelled using either Hodgkin-Huxley models (HHMs) or Markov models (MMs). In HHMs (Hodgkin and Huxley, 1952) the open probability is calculated as the product of several independent ‘gates’, e.g. one gate representing voltage-dependent activation, one representing VDI, and one representing CDI. The opening and closing of each gate is modelled using a single ordinary differential equation (ODE), although CDI gates are frequently assumed to be very fast so that their ODE can be replaced by an analytic equation for their steady state (e.g. the Hill equation). In MMs the channel is postulated to be in one of a finite number of states, and transition rates are defined between the states that can depend on *V_m_* or [Ca^2+^]. In contrast to HHMs, where each process (activation, VDI, CDI) is independent, MMs allow a more general structure where e.g. activation and VDI are coupled (Rudy and Silva, 2006).

In the original and most common formulation, where the open probability (*O*) of a HHM is the product of a number of independent gates, an equivalent MM can be written for each HHM (Keener and Sneyd, 1998). However, many I_CaL_ models use extensions to the classical HH scheme, e.g. they introduce a noninactivating fraction of channels (McAllister et al., 1975, see B in Figure 5), or model the current as if arising from two separate channel families (Nygren et al., 1998). Such models have no obvious single MM equivalent. For examples of MM with and without a HMM counterpart and a graphical explanation of how to convert between the formalisms, see Figure 4 in Rudy and Silva (2006). Taking these equivalences into account, we identified 15 distinct gating models, as shown schematically in Figure 5. Where possible, we have shown models in an MM representation.

**Figure 5:**
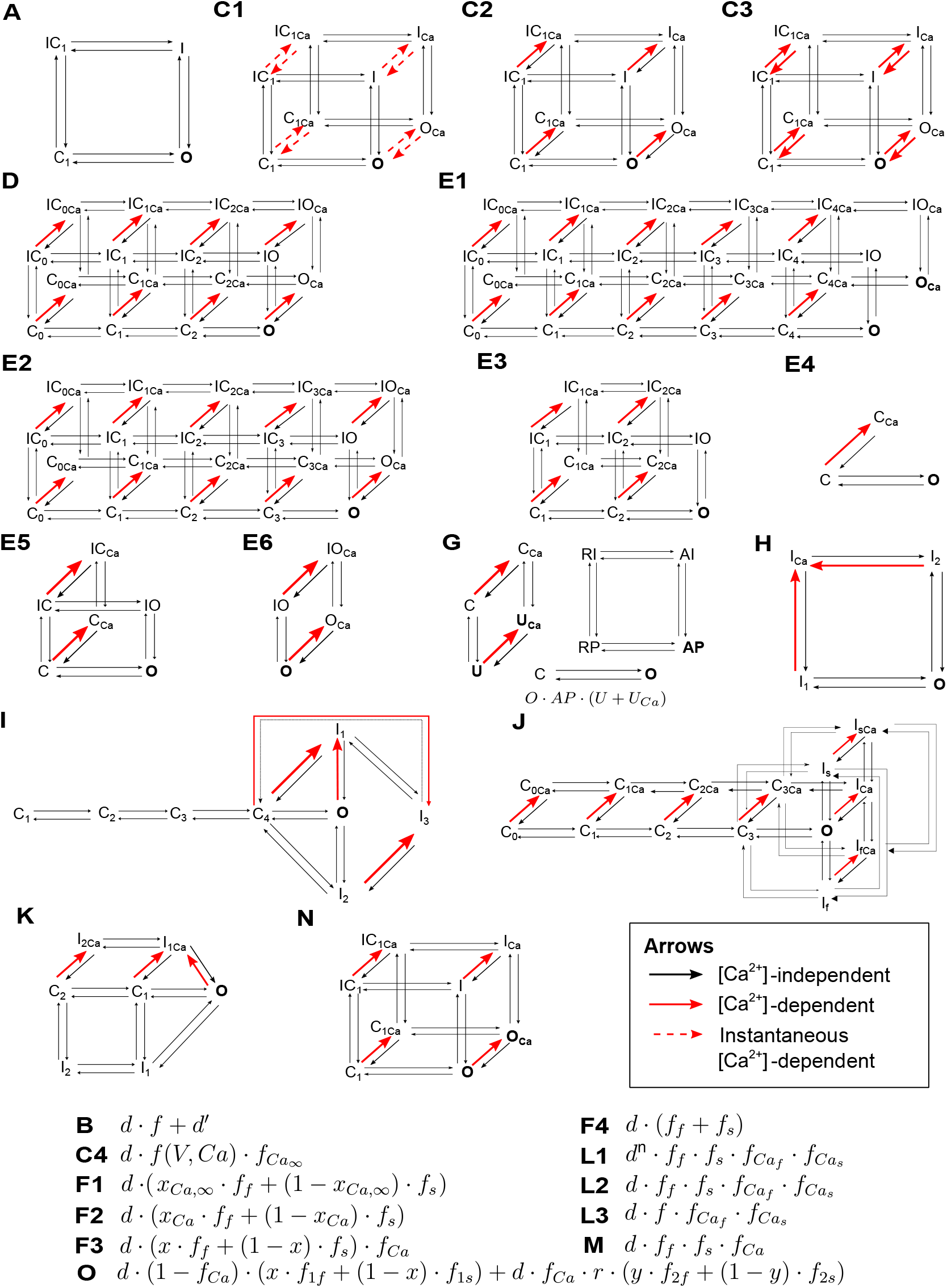
Models of I_CaL_ gating. Each label in this figure (A, C1, C2, …) indicates a distinct gating mechanism, as described in detail in the text. Where possible, models are shown in a MM representation. MM state labels include *C*losed, *I*nactivated, *O*pen, Activated, *U*ncovered, Resting, and *P*rimed. Subscripts *s* and *f* are used to indicate states involved in ‘slow’ and ‘fast’ transitions, while the *Ca* subscripts indicate states involved in calcium-dependent transitions. The conducting states in each Markov structure are shown in bold. Where models have more than one conducting state, the total open fraction is found by adding the occupancy of all conducting states together. The open probability equation for G is shown within the Figure. In the HH equations, *d* and *f* denote activation and inactivation gates respectively, and *f_Ca_* gates are calcium-dependent.

Each gating type is described briefly below. Some models define more than one copy of the same gating model but with changes in parameters or [Ca^2+^] to account for LCC localisation or phosphorylation; we do not focus on these properties in this section and are only concerned with the gating model structure.

**Type A** is the earliest and most straightforward gating mechanism, consisting of voltage-dependent activation (d) and inactivation (*f*), modelled as independent gates that are not affected by [Ca^2+^] (Bassingthwaighte and Reuter, 1972). Wilders et al. (1991) is a unique model within this group because its time constant of inactivation is not voltage-dependent but rather depends on the fraction of inactivation *f*. **Type B** extends type A by adding a non-inactivating fraction of channels *d*′, leading to an open probability *O* = *d* · *f* + *d*′ (McAllister et al., 1975).

The **type C** gating mechanism extends type A by adding a CDI gate, usually written as *f*_Ca_. In **type C1** this is modelled as an instantaneous process, so that the fraction of open *f*_Ca_ gates is given by a Hill equation with Hill coefficient 2 in Luo and Rudy (1994) but 1 in the other C1-type models. **Types C2** and **C3** use an ODE to model the evolution of *f*_Ca_. C2 has calcium-dependent inactivation but calcium-independent recovery (DiFrancesco and Noble, 1985), while both inactivation and recovery are calcium-dependent in C3. **Type C4** has an instantaneous *f*_Ca_ gate, but also incorporates calcium-sensitivity in its gate for VDI, making it a dual voltage-calcium-dependent inactivation gate.

Gating **type D** is similar to type C in its HH form, except that the activation gate d is cubed (Nordin, 1993). In the MM structure, this equates to adding several closed states, as shown in Figure 5D.

An even larger Markov scheme is used by models of **type E1**, in which five steps are required to fully activate (similar to a fifth power in HH terms). In the original implementation, this model was written as a combination between an HHM (for VDI) and an MM for activation and CDI (Jafri et al., 1998). The mechanism for CDI in this model is a switch to a ‘mode Ca’ in which transitions to the open state are extremely slow (Imredy and Yue, 1994). A slight variation, which we still classified as E1, is given by Iyer et al. (2004) who removed the *O*_Ca_ and *OI*_Ca_ states. Several simplifications of the E1 scheme have been proposed, reducing it from 24 to 20 states (**type E2**, Michailova et al., 2005), 10 states (**type E3**, Greenstein and Winslow, 2002), and even 3 states (**type E4**, Hinch et al., 2004). Note that, to arrive at the representation of the E2 type shown here, we have reinterpreted the model by Michailova et al. (2005) as an MM and omitted the term accounting for [MgATP] effects and a constant factor *x*. The study by Li et al. (2010) extended the E4 type by re-adding VDI, creating the 6-state **type E5**. Similarly, Asakura et al. (2014) extended E4 with a fourth state to create **type E6**.

**Type F** gating mechanisms again assume that I_CaL_ has fast and slow modes (citing evidence from Imredy and Yue, 1994; You et al., 1995). In **type F1**, the fraction of channels in slow mode is determined by an instantaneous calcium binding process (Nygren et al., 1998) or fixed to a constant (Kettlewell et al., 2019), while in **type F2** this process is modelled with an ODE (Pandit et al., 2001). Type F3 separates fast and slow kinetics from calcium by fixing the fraction of channels with fast VDI to 73% and introducing a separate CDI process (Pohl et al., 2016). A similar two-component model but without CDI was introduced by Inada et al. (2009), which we have labelled as **type F4**.

The **type G** gating mechanism (based on earlier work by Shirokov et al., 1993, for which we could not find the full equations) consists of three independent MMs for activation, VDI, and CDI (Matsuoka et al., 2003). In the MM for CDI, the binding of a Ca^2+^ ion brings the system into a state where recovery from inactivation is much slower.

A new **type H** gating mechanism was introduced by Lovell et al. (2004), consisting of a four state MM with activation, VDI and CDI. In this model, CDI can only occur when the channel is already in a non-conducting state. More complex MMs were introduced by Bondarenko et al. (2004), Faber et al. (2007), and Mahajan et al. (2008b) to create **types I**, **J**, and **K** respectively. The scheme in type I was inspired by structure-function relationships in a homology model based on *Shaker B* voltage-gated *K*^+^ models. Type J includes a fast and a slow VDI, to reflect gating current experiments by Ferreira et al. (2003). The Type K model incorporates ideas from Soldatov (2003) and Cens et al. (2006), who argued that CDI is a voltage-dependent process that is greatly accelerated when Ca^2+^ binds to the channel (see also Pitt et al., 2001). Its Markov scheme is reduced by combining two open states into one, and reducing the five-state activation to a three-state process (Rose et al., 1992).

The **type L1** gating mechanism by Hund and Rudy (2004) consists of activation, fast and slow VDI, and fast and slow CDI. In this model, CDI depends on [Ca^2+^]_d_ but also has an I_CaL_-dependence, to approximate the raised [Ca^2+^] at the cytosolic channel mouth (this was based on the model by Hirano and Hiraoka, 2003, for which we could not find the full equations). The activation variable is raised to a time- and voltage-dependent power *n* (modelled as an ODE) which accounts for a ‘fast voltage-dependent facilitation’ described by Kamp et al. (2000). In **type L2** this is simplified to use a constant *n* = 1 (Aslanidi et al., 2009b), and **type L3** simplifies this further by combining fast and slow VDI into a single gate (Hund et al., 2008). All type L models have a fast CDI gate (*f_Ca_f__*) that accounts for CDF by CaMKII-dependent phosphorylation.

The study by Ten Tusscher and Panfilov (2006) extended their previous C3 type model with an extra gate, leading to **type M**. Note that this can also be seen as a simplification of type L2.

The MM scheme for **Type N** is somewhat similar to C2, but with the important difference that it has two conducting states (so that this MM can no longer be reduced to an equivalent HHM). Like type K, this gating model is based on the idea that Ca^2+^ binding in CDI ‘removes a brake’ on a voltage-dependent process (Pitt et al., 2001). This is very similar to the ‘mode-switching’ idea underlying E1. Heijman et al. (2011) uses a variation of this scheme, in which the rates of CDI and VDI are modulated by [CaMKII]. In addition, this model uses a second copy of the same MM structure but with modified rates to represent PKA-phosphorylated (*β*-adrenergically stimulated) LCCs. Another variation is used by Bartolucci et al. (2020), who borrow two state variables from the model by O’Hara et al. (called *n* and *j* in the publication, but shown as *f*_Ca_ and *r* respectively in Figure 5O) which are used to determine the transition rate into the ‘Ca’ states. As in the model by Heijman et al., two copies of the same model are maintained, but this time the phosphorylated population is CaMKII-rather than PKA-phosphorylated.

**Type O** gating, introduced in (O’Hara et al., 2011), splits the channels into several fractions. First, a [Ca^2+^] dependent variable (called the ‘n-gate’ in the original publication but *f*_Ca_ in our notation) splits the channels into a fraction in ‘VDI mode’ and a fraction in ‘CDI mode’. As in types K and N, the underlying assumption is that Ca^2+^ binding (modelled by the ‘n-gate’) brings the LCCs into a state where a voltage-dependent ‘CDI’ process can occur (Pitt et al., 2001; Kim et al., 2004). In turn, each fraction (VDI mode and CDI mode) is split into a fast and slow fraction, with a fixed ratio (*x* in Figure 5) between fast and slow ‘VDI-mode inactivation’ and a voltage-dependent ratio (*y* in Figure 5) between fast and slow CDI-mode inactivation. All three type O models in this review include CaMKII-phosphorylation via a second copy of the model with altered rate equations.

Table 2 shows how each model fits into this classification. In addition to the gating type, it lists each model’s cell type, species, and the simulated temperature (where stated). The final column provides an indication of the major data sources used in the model’s construction. For the earlier generation of models, this data source is often hard to establish, as parameters were commonly set by hand to create a model that was acceptably close to a wide range of observations (typically measured in tissue preparations, and sometimes spanning several species). A full list of data sources is available online at https://github.com/CardiacModelling/ical-model-review.

**Table 2:**
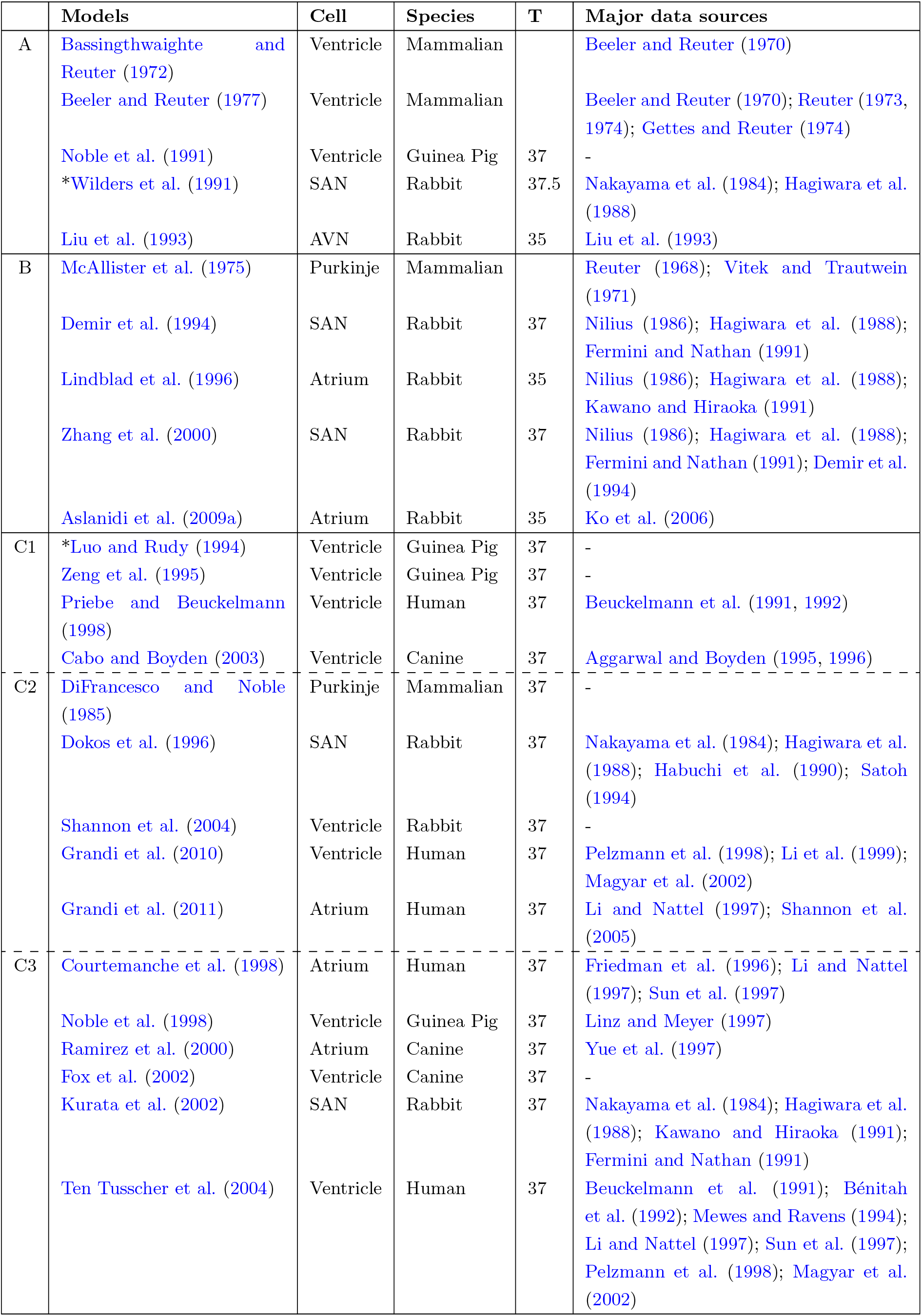
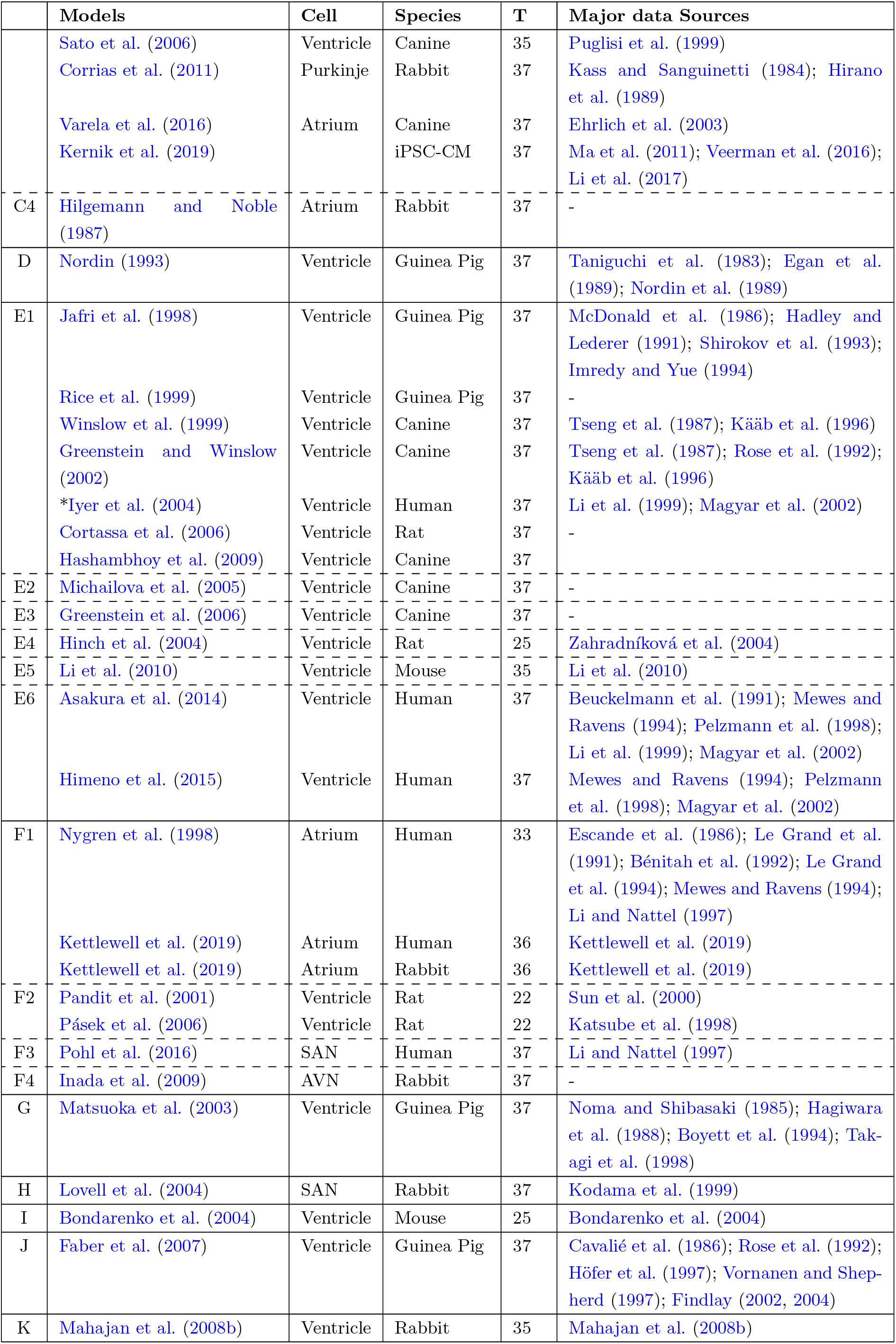
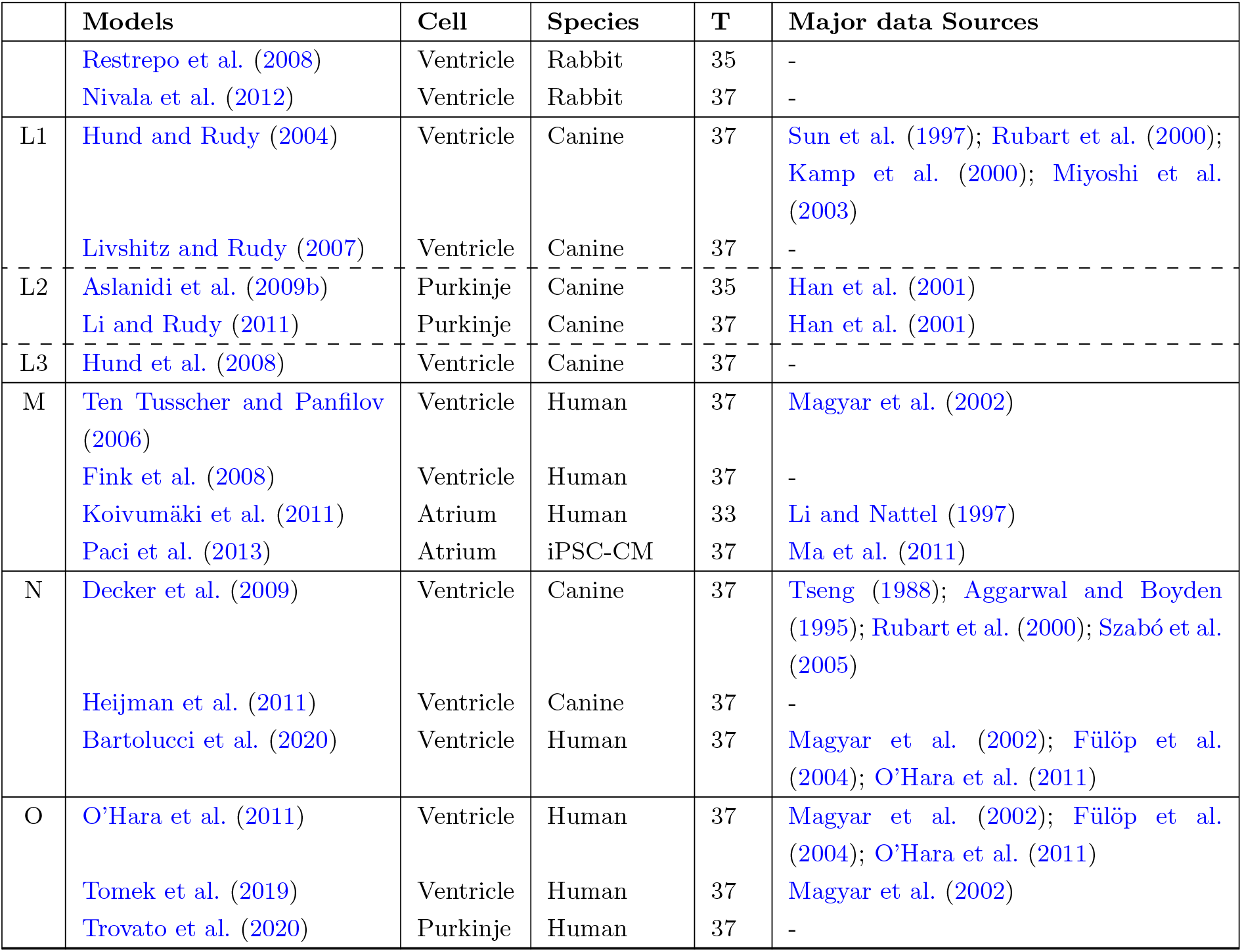
Models classified by their gating type, with additional information of cell type, species, temperature (T) and the major data sources used in each model’s construction. The equivalent MM schematic/equation for each gating class is shown in Figure 5. Minor variants within each class are indicated with (*) and explained in text.

### 2.5 Driving force

Two types of driving force are common in I_CaL_ models. An Ohmic driving force for a current carried by a single species takes the form

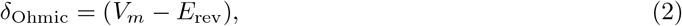

where *E*_rev_ is the *reversal potential*, at which the current reverses direction. For currents carried by a single species, the reversal potential can be calculated using the Nernst equation

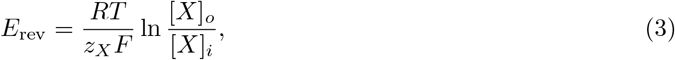

where *R* is the universal gas constant, *T* is temperature, *F* is the Faraday constant, *z_x_* is the valence of a single ion of species *X* (2 for [Ca^2+^]), and [*X*]_*o*_ and [*X*]_*i*_ are the external and internal concentrations of *X* respectively. Although commonly used in the form given above, a more accurate version uses *γ_o_*[*X*]_*o*_/*γ_i_*[*X*]_*i*_, where *γ_o_* and *γ_i_* are dimensionless *activity coefficients* that account for non-ideal behaviour of the solutes. The units of the gas constant are frequently chosen as mJ/K/mol to yield a reversal potential in mV. A model with an Ohmic driving term is written 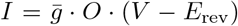, where 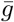 is in Siemens (S) for a current in Amperes (A).

A more complex model of the electrochemical driving force is given by the Goldman-Hodgkin-Katz (GHK) flux equation (Goldman, 1943; Hodgkin and Katz, 1949):

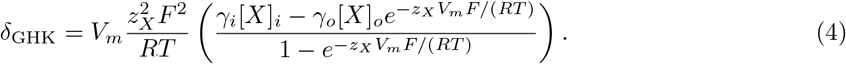

Because its derivation involves assuming a constant electrical field throughout the channel, the GHK equation and its derivatives are also known as ‘constant field theory’. Assuming concentrations in mM = mol/m^3^ and using matching units for *V* and *RT/F*, a model with a GHK driving term is written as 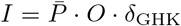, where 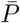 is in L/ms (= m^3^/s) for a current in A, in L/F/ms for a current in A/F, or in cm/s for a current in *μ*A/cm^2^.

A number of early studies used a modified GHK equation, which accounts for the (hypothesised) presence of a charged particle near the channel mouth, causing a change, *V*_0_, in the voltage across the channel (Frankenhaeuser, 1960):

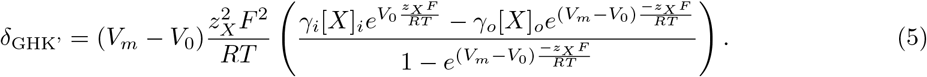

So far, we have assumed a current carried exclusively by Ca^2+^, i.e. a perfectly selective channel. For this situation, *δ*_Ohmic_, *δ*_GHK_, and *δ*_GHK_, all predict a current that reverses at the potential given by Equation 3. For typical concentrations, this would lead to a current that reverses in the range 50–150 mV, which is significantly higher than experimental estimates of 40–70 mV. If, however, we assume that the channel is even very slightly permeable to Na^+^ and K^+^, then the much higher internal concentrations of those ions compared to [Ca^2+^] has a significant effect on the reversal potential.

A simple model for currents with multiple carriers can be made by assuming that each species travels through the channel without affecting the others, so that the total current is simply a sum of single-species currents (independent flux assumption). Equations for *E*_rev_ can still be derived with this assumption (see e.g. Campbell et al., 1988; Keener and Sneyd, 1998), but the predicted reversal potentials from Ohmic and GHK models are no longer the same. In both equations, the reversal potential of a multi-species current depends strongly on the ratio between the species’ permeabilities 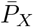 (or, if an Ohmic term is used, their conductances 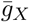). A study by Campbell et al. (1988) measured the reversal potential of I_CaL_ in bull-frog atrial myocytes, and compared this to predictions using different ratios of 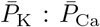. They found that good agreement with experiment could be found using a GHK equation (Equation 4), provided the channel was highly selective to [Ca^2+^] (>95% of the I_CaL_ is carried by [Ca^2+^] at 0mV).

To compare different models’ driving terms, we set internal and external concentrations to fixed levels, and plotted the driving term as a function of voltage in Figure 6. We used [Na^+^]_*o*_ = 140 mM, [Na^+^]_*i*_ = 10 mM, [K^+^]_*o*_ = 5 mM, [K^+^]_*j*_ = 140 mM, [Cl^-^]_*o*_ = 150 mM, [Cl^-^]_*i*_ = 24 mM, [Ca^2+^]_o_ = 2 mM and [Ca^2+^]_s_ = [Ca^2+^]_i_ = [Ca^2+^]_d_ = 10^-4^ mM. (Note that these values may differ from the concentrations that the models were designed for, the concentrations used in the experiments they were calibrated with, or the physiological concentrations in real cells.) In models where I_CaL_ is carried by multiple species, we calculated the ‘net driving term’ by summing up the contributions of each component, and weighting by the component’s permeability relative to the calcium component (e.g. the calcium component has weight 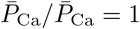, the potassium component has weight 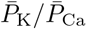, etc.).

**Figure 6:**
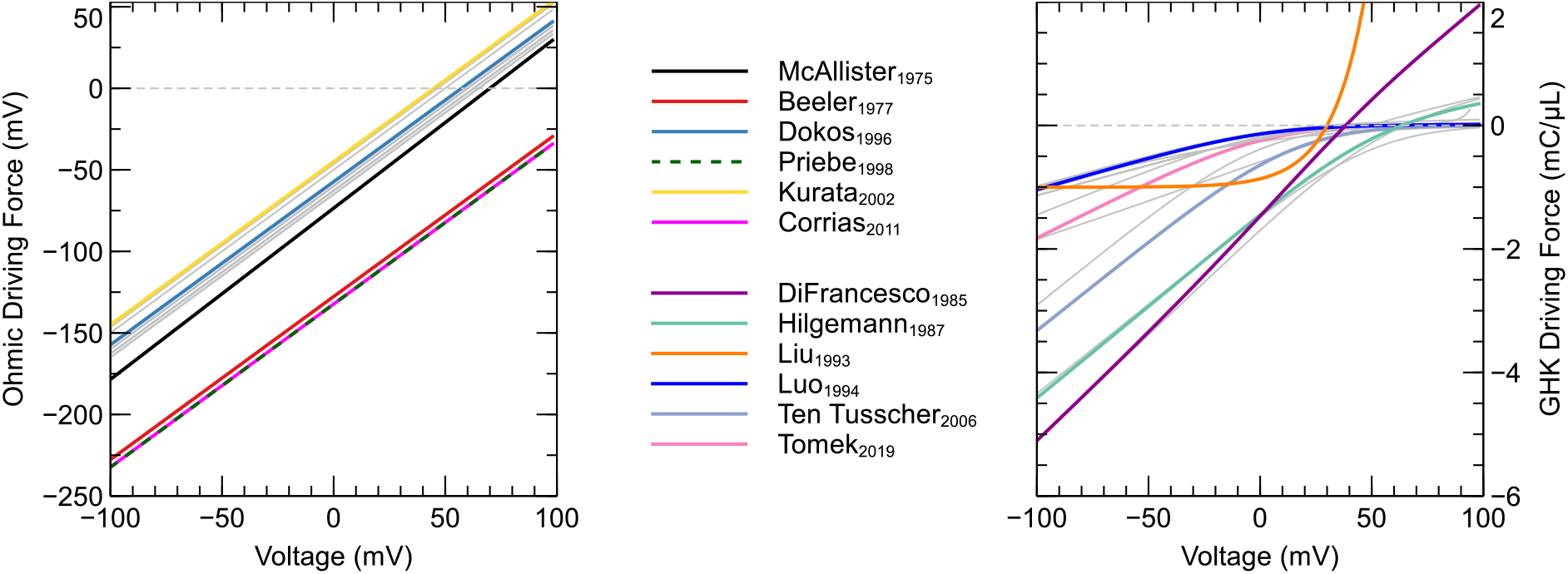
Driving term as a function of voltage for models with an Ohmic driving term (**Left**) and a (normal or modified) GHK driving term (**Right**). In the left panel, the majority of models are plotted in grey and are shown to form a tight cluster, reversing at around 60 mV. The lower and upper bounds for this cluster are marked by McAllister et al. (1975) and Kurata et al. (2002), which are indicated in colour, along with three outliers (Beeler and Reuter, 1977; Priebe and Beuckelmann, 1998; Corrias et al., 2011) that do not show a reversal potential within the physiological range. In the right panel, most currents are again shown in grey, but a selection of models are shown in colour to show the range of behaviours observable by varying the activity coefficients and voltage offset V_o_ (DiFrancesco and Noble, 1985; Hilgemann and Noble, 1987; Luo and Rudy, 1994; Hund and Rudy, 2004; Ten Tusscher and Panfilov, 2006; Tomek et al., 2019). An interactive view of these results is available at https://chaste.cs.ox.ac.uk/q/2O22/ical/fig6.

The left panel of Figure 6 shows *δ* for the models in this study that used an Ohmic driving term (models for which we could not find the full equations are omitted). Rather than use the Nernst equation, most of these models set a constant reversal potential in the range of 45–70 mV. As expected, models that do use the single-species Nernst equation (without any modifications) (Priebe and Beuckelmann, 1998; Corrias et al., 2011) show a reversal potential well outside of the physiological range. The right panel in Figure 6 shows *δ* for models with a GHK (or GHK’) driving term. Most of these models do not show a true reversal of current, with the driving force approaching zero for high voltages. Only models with a high permeability for sodium and potassium ions (DiFrancesco and Noble, 1985; Hilgemann and Noble, 1987) show a positive driving force for high voltages. Finally, the model by Liu et al. (1993) uses neither an Ohmic nor a GHK driving term, but an exponentially increasing flux and is shown in the right panel.

A list of models with Ohmic driving terms is given in Table 3. For models using a fixed *E*_rev_, the numerical value is shown, while the equation used for *E*_rev_ is shown for the remainder. None of the models with Ohmic driving terms in this study explicitly modelled potassium or sodium components of I_CaL_ (although their existence may be implicitly acknowledged in the models with a fixed reversal potential).

**Table 3:**
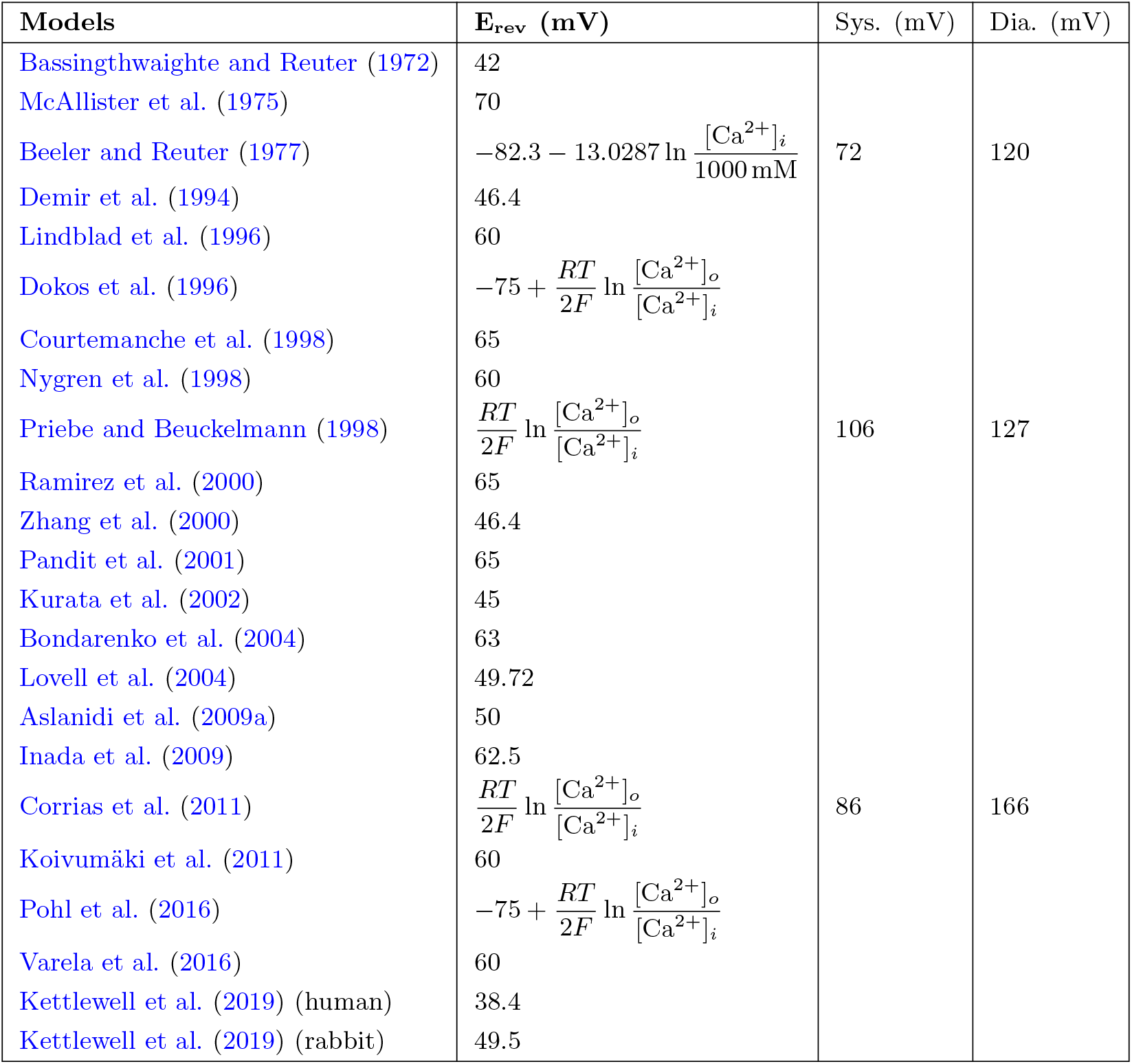
Reversal potentials for models using an Ohmic driving force (Equation 2). The final two columns provide representative E_rev_ values during the AP (‘systole’) and in the refractory phase (‘diastole’), for models in which E_rev_ is given by an equation and for which a full AP model was available.

A similar list for models with GHK (and GHK’) driving is given in Table 4, along with the offset *V*_0_ (if used), the activity coefficients for internal and external calcium, and the ratio between the permeability coefficients of potassium and sodium with respect to calcium.

**Table 4:** GHK and GHK’ parameters in models using GHK driving terms (Equation 5). Models marked with a (*) use variations of the GHK equation (as discussed in the main text).

| Models | $V_o$ (mV) | $\gamma_i$ | $\gamma_o$ | $\bar{P}_K : \bar{P}_{Ca}$ | $\bar{P}_{Na} : \bar{P}_{Ca}$ |
| --- | --- | --- | --- | --- | --- |
| DiFrancesco and Noble (1985) | 50 | 1 | 1 | 1:100 | 1:100 |
| Hilgemann and Noble (1987) | 50 | 1 | 1 | 1:500 | 1:100 |
| Noble et al. (1991) | 50 | 1 | 1 | 1:500 | 1:100 |
| Wilders et al. (1991) | 50 | 1 | 1 | 1:100 | 1:100 |
| Nordin (1993) | 50.1 | 1 | 1 | 1:197.3 | - |
| Luo and Rudy (1994) | 0 | 1 | 0.341 | 1:2798 | 1:800 |
| Zeng et al. (1995) | 0 | 1 | 0.341 | 1:2798 | 1:800 |
| *Jafri et al. (1998) | 0 | 1 | 0.341 | $f(V, Ca)$ | - |
| Noble et al. (1998) | 50 | 1 | 1 | 1:500 | 1:500 |
| *Rice et al. (1999) | 0 | 1 | 0.341 | - | - |
| *Winslow et al. (1999) | 0 | 1 | 0.341 | $f(V, Ca)$ | - |
| Fox et al. (2002) | 0 | 1 | 0.341 | $f(V, Ca)$ | - |
| Greenstein and Winslow (2002) | 0 | 1 | 0.341 | - | - |
| Cabo and Boyden (2003) | 0 | 1 | 0.341 | 1:1554 | 1:444 |
| Matsuoka et al. (2003) | 0 | 1 | 1 | 1:2730 | 1:54054 |
| Hinch et al. (2004) | 0 | 1 | 1 | - | - |
| Hund and Rudy (2004) | 15 | 1 | 0.341 | - | - |
| *Iyer et al. (2004) | 0 | 1 | 0.341 | $f(V, Ca)$ | - |
| Shannon et al. (2004) | 0 | 0.341 | 0.341 | 1:2000 | 1:36000 |
| Ten Tusscher et al. (2004) | 0 | 1 | 0.341 | - | - |
| *Michailova et al. (2005) | 0 | 1 | 0.341 | 1:540 | - |
| *Cortassa et al. (2006) | 0 | 1 | 1 | $f(V, Ca)$ | - |
| Greenstein et al. (2006) | 0 | 1 | 0.341 | - | - |
| Sato et al. (2006) | 0 | 1 | 0.341 | - | - |
| Pásek et al. (2006) | 0 | 1 | 0.341 | - | - |
| *Ten Tusscher and Panfilov (2006) | 15 | 0.25 | 1 | - | - |
| Faber et al. (2007) | 0 | 0.01 | 0.341 | 1:2780 | 1:800 |
| Livshitz and Rudy (2007) | 15 | 0.341 | 0.341 | - | - |
| *Fink et al. (2008) | 15 | 0.25 | 1 | - | - |
| Hund et al. (2008) | 0 | 1 | 0.341 | - | - |
| Mahajan et al. (2008b) | 0 | 1 | 0.341 | - | - |
| Restrepo et al. (2008) | 0 | 0.341 | 0.341 | - | - |
| Aslanidi et al. (2009b) | 15 | 1 | 0.341 | - | - |
| Decker et al. (2009) | 0 | 1 | 0.341 | - | - |
| Hashambhoy et al. (2009) | 0 | 1 | 0.341 | - | - |
| Grandi et al. (2010) | 0 | 0.341 | 0.341 | 1:2022 | 1:36000 |
| Li et al. (2010) | 0 | 1 | 1 | - | - |
| Grandi et al. (2011) | 0 | 0.341 | 0.341 | 1:2000 | 1:36000 |
| Heijman et al. (2011) | 0 | 1 | 0.341 | - | - |
| Li and Rudy (2011) | 15 | 1 | 0.341 | - | - |
| O'Hara et al. (2011) | 0 | 1 | 0.341 | 1:2798 | 1:800 |
| Nivala et al. (2012) | 0 | 1 | 0.341 | - | - |
| Paci et al. (2013) | 0 | 1 | 0.341 | - | - |
| Asakura et al. (2014) | 0 | 1 | 1 | 1:682 | 1:54054 |
| Himeno et al. (2015) | 0 | 1 | 1 | 1:2730 | 1:54054 |
| Kernik et al. (2019) | 0 | 0.341 | 0.341 | 1:2000 | 1:36000 |
| Tomek et al. (2019) | 0 | $f$ | $f$ | 1:2798 | 1:800 |
| Bartolucci et al. (2020) | 0 | 1.2 | 0.341 | 1:2801 | 1:800 |
| Trovato et al. (2020) | 0 | 1 | 0.341 | 1:2798 | 1:800 |

The earliest studies in this table all use a voltage shift parameter *V*_0_ = 50 mV, but this form is rarely seen in later models. Models with a large *V*_0_ also assume a large contribution of sodium and potassium ions, consistent with the analysis by Campbell et al. (1988).

The model by Luo and Rudy (1994) abandons the use of *V*_0_ and introduces the values *γ_i_* = 1 and *γ_o_* = 0.341 for the activity coefficients of internal and external calcium. These values can be calculated using the Pitzer equations (Pitzer and Mayorga, 1973), while the frequently used value *γ_i_* = 1 appears to arise from assuming that low internal [Ca^2+^] leads to near-ideal behaviour. Instead of using constant values, the model by Tomek et al. (2019) calculates the activity coefficients using the Davies equation (Davies and Malpass, 1964), which is a precursor to Pitzer’s work (although the first version of this model erroneously used a natural logarithm instead of a base-10 logarithm for the calculation, see https://bit.ly/3tqD4gP). Activity coefficients for sodium and potassium are usually taken to be 0.75 internally and externally, and are not shown in the table.

Although there is a common consensus that I_CaL_ is carried by multiple ion species, the ratio of permeabilities between the different species varies greatly between models, and several studies have cited the low permeability to potassium and sodium as a reason to omit these components from the model altogether (avoiding their impact on the reversal potential by using a fixed rather than a calculated value). Estimates of LCC selectivity have gone up over time: in general, older models assume larger contributions from sodium and potassium, while more recent models assume a much more selective channel. The high selectivity of LCCs is especially evident when comparing values from Table 4 to estimated selectivity ratios of other channels, e.g. 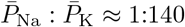 for *I*_Kr_ (Sanguinetti et al., 1995), while 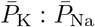 has been estimated as 1:10 for *I*_Na_ (Amin et al., 2005).

Modellers starting with Jafri et al. (1998) have attempted to incorporate evidence that monovalent permeation goes down when there is a significant Ca^2+^ influx — i.e. that the independent flux assumption does not hold. In these models, which consider only a calcium and a potassium component, the permeability to potassium is a function of the calcium driving force, so that 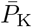 goes down when 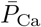 is high (Jafri et al., 1998; Winslow et al., 1999; Fox et al., 2002; Iyer et al., 2004; Cortassa et al., 2006). This is indicated with *f* (*V*, Ca) in Table 4. Ca^2+^ influx, on the other hand, seems able to continue even when net I_CaL_ is outward (Zhou and Bers, 2000). (It is also interesting to speculate whether Ca^2+^ flux might reverse *locally*, e.g. when a high [Ca^2+^] is reached in the dyad.)

Finally, some of the models in Table 4 use a slight variation of the (normal or modified) GHK equation. The model by Jafri et al. (1998) and many of its descendants, (Rice et al., 1999; Winslow et al., 1999; Iyer et al., 2004; Michailova et al., 2005; Cortassa et al., 2006) use a constant internal calcium activity *γ_i_*[Ca^2+^]_*i*_ = 0.001 mM (Smith, 1996). The model by Ten Tusscher and Panfilov (2006) uses a GHK equation in which V is offset by 15 mV, but without the extra term 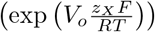 used in Equation 5.

## 3 Quantitative comparison

In this section, 60 models are reviewed and compared *functionally* (Cooper et al., 2011) by simulating voltage-clamp protocols and studying the predicted I_CaL_ response. Voltage-step experiments are simulated to compare activation, voltage-dependent inactivation and recovery, and CDI. To compare the predicted I_CaL_ during an AP, we perform simulations where either the AP (AP-clamp) or AP and CaT (AP-CaT-clamp) are ‘clamped’ to a predetermined time-course.

### 3.1 Ionic concentrations

Extracellular concentrations were set to [Na^+^] = 140 mM, [K^+^] = 5mM, [Cl^-^] = 150 mM, and [Ca^2+^] = 2mM, and kept constant at these values. Similarly, intracellular concentrations were fixed to [Na^+^] = 10 mM, [K^+^] = 140 mM, and [Cl^-^] = 24 mM in all compartments. This corresponds to a patch-clamp setting where these concentrations are well buffered by the bath and pipette solutions. Except where stated otherwise, the intracellular Ca^2+^ concentrations ([Ca^2+^]_*i*_, [Ca^2+^]_*d*_, and [Ca^2+^]_*s*_) were fixed to 0.1 μM. These fixed concentrations allow the current kinetics to be compared in the absence of channel localisation effects (but may vary from the values for which individual models were designed); we will revisit the impact of this choice later. Variables representing [Ca^2+^] in the SR and [CaMKII] were left to vary as dictated by the model equations. However, these variables remain constant (or close to constant) when all internal [Ca^2+^] are fixed.

### 3.2 Simulation methods

As the basis for our simulations, we used CellML 1.0 or 1.1 files (Hedley et al., 2001) containing either a single I_CaL_ model, or an I_CaL_ model embedded into a larger model of the AP. When simulating basic voltage-clamp protocols (activation, VDI, recovery, CDI) AP models were reduced to I_CaL_-only models by fixing the values of *V_m_* and all external and internal ionic concentrations to predetermined levels (see Sections 3.2.2 and 3.1). In the AP- and AP-CaT-clamp simulations, some non-I_CaL_ model features are preserved (e.g. the internal calcium and CaMKII dynamics), so that these simulations require a full AP model (see Section 3.6).

Where possible, model implementations were obtained from the Physiome Model Repository (PMR, Yu et al., 2011). Thirty-five models obtained this way could be used directly, while a further thirteen needed corrections to either units or equations. To limit the scope of this correction work, five models were reduced to I_CaL_-only models in the process. Eight models were corrected in AP form, after which the new versions were uploaded to the PMR. A file for the model by Pásek et al. (2006) was also obtained from the PMR. This model has a separate variable for the voltage across the T-tubular membrane, distinct from the ‘main’ transmembrane potential. To compare the channel kinetics consistently, we modified this model manually to set this secondary voltage equal to *V_m_*.

For eleven models no CellML files were available so new implementations were created based on published equations or code. An AP model implementation was created for Heijman et al. (2011), while we created CellML models containing only I_CaL_ for Wilders et al. (1991); Liu et al. (1993); Nordin (1993); Cabo and Boyden (2003); Cortassa et al. (2006); Sato et al. (2006); Faber et al. (2007); Hund et al. (2008); Li and Rudy (2011); Kernik et al. (2019).

This resulted in a set of 60 model files that could be used to simulate I_CaL_ in isolation, and a subset of 44 files that provided a full model of the AP.

#### 3.2.1 Software

Simulations were run using the Cardiac Electrophysiology Web Lab (Cooper et al., 2016; Daly et al., 2018). This is a resource developed to allow the characterisation and comparison of cardiac electrophysiology models in a wide range of experimental scenarios. It uses a novel protocol language that allows the user to separate the details of the mathematical model from the experimental protocol being simulated. We also performed simulations for calcium sensitivity (see Section 3.5) by solving the model equations using CVODE version 3.1.2 as incorporated in Myokit (Clerx et al., 2016) version 1.28.4, using Python version 3.7.3. All model and simulation files are available under an open source BSD 3-Clause license, and can be obtained from https://github.com/CardiacModelling/ical-model-review (doi: 10.5281/zenodo.6653898).

#### 3.2.2 Model annotations and modifications

To run simulations in the Web Lab, models are encoded in CellML format and annotated with metadata terms. These terms allow variables in different models with the same physiological meaning to be identified, read out in units of choice, and manipulated. This is a useful feature for this study as the 60 models do not use consistent naming.

The following variables were annotated in all models: time, membrane potential (*V_m_*), the total I_CaL_, driving term (*δ*), and the open probability (*O*). When annotating driving term variables, distinct meta-data terms were used for Ohmic and GHK formulations. When annotating I_CaL_ in models that split the current into multiple components (by ionic species, phosphorylated/non-phosphorylated fraction, or biological subcompartment) we selected the variable that represents the sum over all these aspects. For this purpose, a new variable was often introduced in the CellML files. Similarly, *O* is sometimes split into components with or without phosphorylation, *δ* can be split by ionic species, and both *O* and *δ* are occasionally split by the biological compartment. For all such cases, we introduced and annotated new *O* and *δ* variables as the weighted average over all these components (see e.g., the annotations in our file Bartolucci et al., 2020). This weighting was made based on the fraction of phosphorylated channels, the relative permeability of each ionic species with respect to calcium, and/or the fraction of channels in each biological compartment. For example, where ICaL was calculated as a linear combination of a calcium, potassium, and sodium component

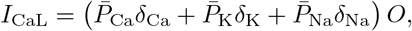

a weighted term for a GHK driving force and current carried by three ion species was calculated as:

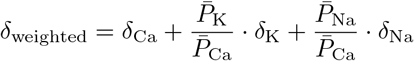

where *δ*_Ca_, *δ*_K_, and *δ*_Na_ are the driving force terms calculated for each species separately.

Finally, we annotated the variables representing intra- and extra-cellular concentrations of Ca^2+^, Na^+^, K^+^, and Cl^-^. For external concentrations, most models use a single constant-parameter but some make a distinction between a constant ‘bath concentration’ and a time-varying ‘cleft concentration’ near the cell. All three types of external concentration (‘bath’, ‘cleft’, or simply ‘external’) were annotated and manipulated by the Web Lab during simulation. Similarly, we annotated (and later manipulated) intracellular concentrations for bulk cytosol, submembrane space, and dyadic space concentration.

### 3.3 Voltage dependence of activation

To show model predictions of voltage-dependence of activation, a voltage-clamp protocol was adapted from O’Hara et al. (2011). This is a typical example of the type of protocol commonly used to study I_CaL_ activation experimentally, but there are some notable differences between these experiments and our simulation. First, the simulated voltage is set *exactly* to the values dictated by the protocol (corresponding to an ideal rather than a realistic voltage clamp, see Lei et al., 2020a). Secondly, internal and external concentrations are held constant during the simulation (see 3.1), so that CDI is kept at a fixed (but model-dependent) level. When using AP models, this ‘clamping’ of voltage and concentrations also serves to remove all interaction with other ionic currents, so that this is a simulation of I_CaL_ in isolation.

The activation protocol consists of repeating units called *sweeps*. Each sweep starts with a 20 s segment where *V_m_* is kept at the *holding potential* of −90 mV, followed by a 120 ms step to the *test potential*, denoted by P1 in Figure 7 (left). The test potential P1 is −60mV for the first sweep and increases by 2 mV per sweep up to +60 mV in the final sweep (for clarity, only a subset of these fine-increment sweeps is shown in the figure). In our simulations, current was recorded during P1 to measure the peak I_CaL_ (denoted I) at each sweep (Figure 7, centre) which was then normalised as 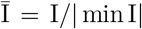. Next, the normalised peak current, 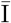, was plotted against the test potential for all models, leading to the normalised I-V curves shown in Figure 7 (right). We also calculated the midpoint of activation (*V*_0.5_) for each model, by finding the point where the normalised I-V curve first crosses −0.5.

**Figure 7:**
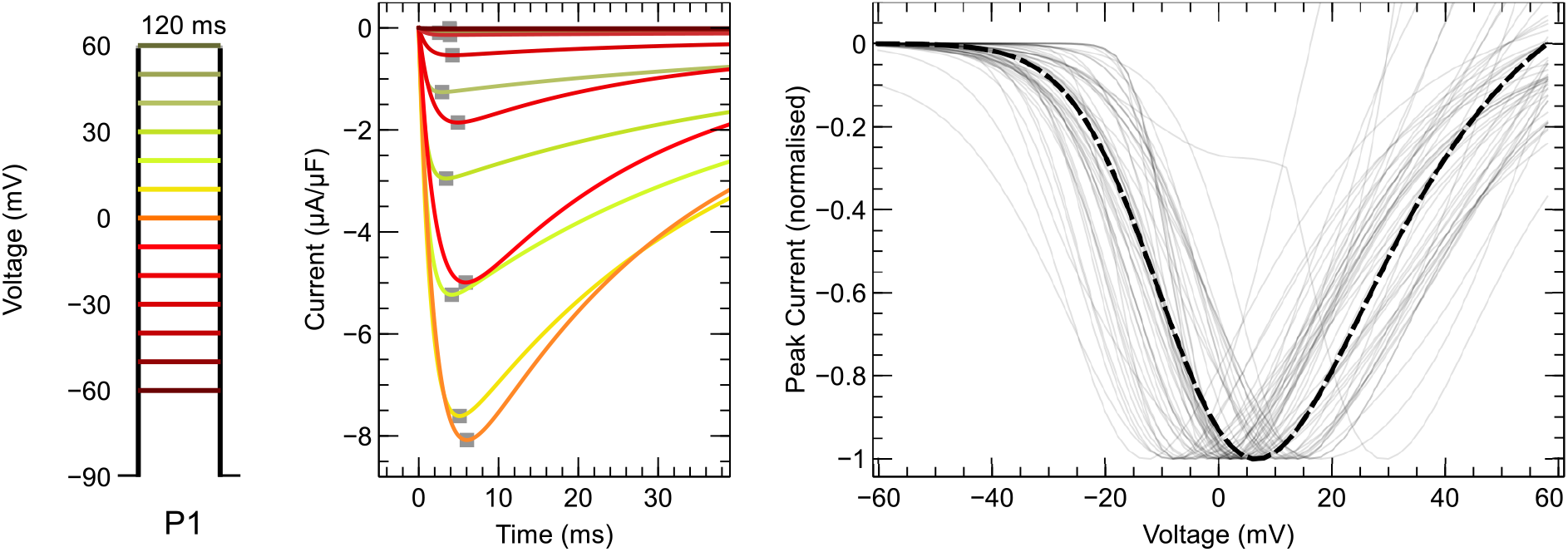
**Left**: Voltage-clamp protocol used to characterise activation (see main text for details). **Centre**: The current predicted by the Grandi et al. (2010) model, simulated with fixed internal and external concentrations and a constant level of CDI. Traces from successive sweeps are overlaid and colour-coded to match the P1 steps in the first panel. The peak I_CaL_ at each voltage step is marked with a grey square. These values are used to construct the I-V curves on the right, as shown in the animation online at https://github.com/CardiacModelling/ical-model-review. **Right**: Normalised peak I_CaL_ versus step voltage for 56 (out of 60) models (the direction of the current was preserved during normalisation). The black dashed line shows the I-V curve from Faber et al. (2007), which is close to the median response. The models by McAllister et al. (1975), Beeler and Reuter (1970), Liu et al. (1993), and Corrias et al. (2011) showed outlier responses, and are plotted separately in Figure 8. An interactive view of these results is available at https://chaste.cs.ox.ac.uk/q/2022/ical/fig7.

Most I_CaL_ models predict that channels begin to activate at membrane voltages around −40 *mV* and that I_CaL_ reaches its maximum magnitude between −20 to +20 mV. The *V*_0.5_ varies between −48 to +15.3 mV for the models in this study while the median *V*_0.5_ is −13.5 mV. By comparison, examples from the experimental (patch-clamp) literature show that LCCs start to activate at around −40 mV, that peak I_CaL_ is observed between about −5 to 10 mV, and that *V*_0.5_ lies between −25 to −10 mV (Pelzmann et al., 1998; Van Wagoner et al., 1999; Magyar et al., 2000).

In Figure 8 we take a closer look at the outlier models not included in Figure 7. The left panel shows the I-V curve for all 60 models, including outliers which are highlighted in colour. This includes the two oldest models in our quantitative review (McAllister et al., 1975; Beeler and Reuter, 1977), both of which are based on tissue rather than isolated-cell measurements and so a difference in their activation kinetics is not surprising. However, some more interesting effects are observed when we dissect the currents into open probability and driving force, as is done in the right panel which shows *normalised* open probability versus voltage (defined as 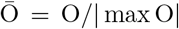). Of the 4 outliers, only the McAllister et al. (1975) model has an unusual O-V curve, causing peak I_CaL_ to be reached at a lower voltage than other models. The models by Beeler and Reuter (1977), and Corrias et al. (2011) have more typical O-V curves, but exhibit unusually large currents at higher voltages (left panel) due to their unusually large driving force (see Figure 6 and Table 3). Interestingly, the model by Priebe and Beuckelmann (1998) has a similar driving term (Figure 6), but this is compensated by an outlier O-V curve, leading to a regular I-V curve (Figure 8, but see also Figure 17). Finally, the model by Liu et al. (1993) has an unusual I-V curve not due to its O-V curve but due to its exponentially increasing driving term. From this dissection of I_CaL_ into O and *δ* we can see how irregularities in one aspect of a model (e.g. *O*) can sometimes be compensated by irregularities in another (e.g. *δ*).

**Figure 8:**
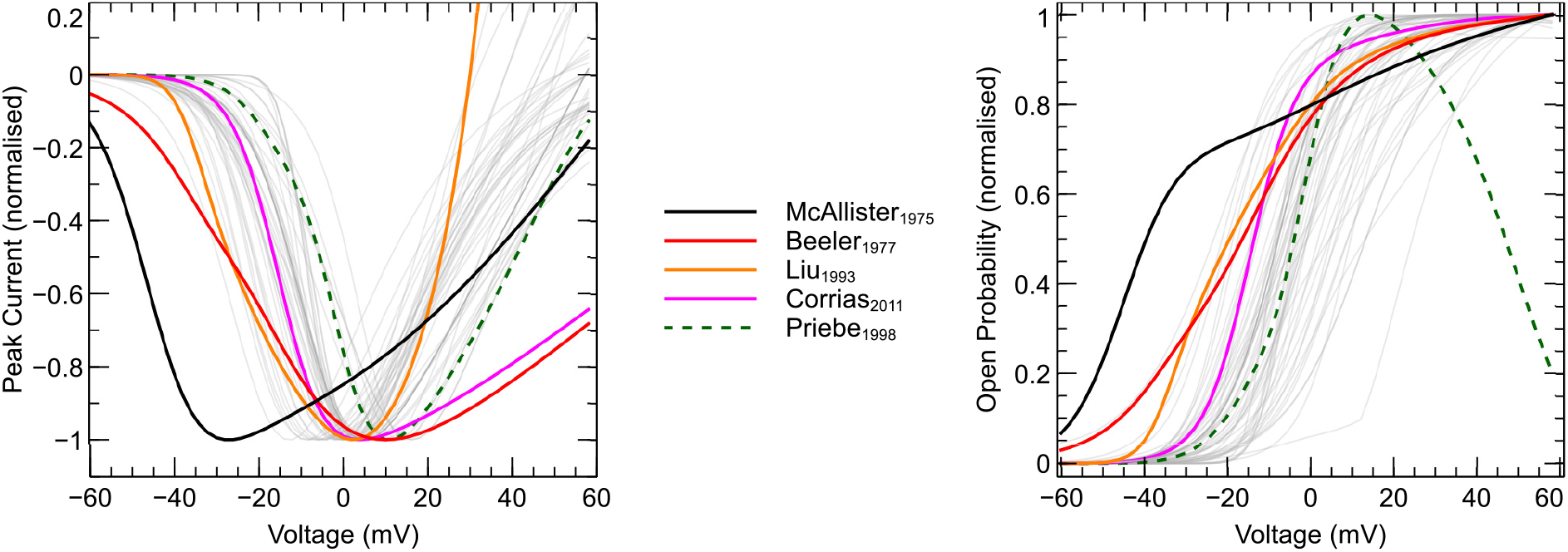
**Left**: I-V curve and **Right**: Normalised peak-open probability plotted against test potential for all 60 models. The 4 outliers and the model by Priebe and Beuckelmann (1998) are highlighted in colour and discussed in the main text; the others are shown in grey. An interactive view of these results is available at https://chaste.cs.ox.ac.uk/q/2O22/ical/fig8.

### 3.4 Voltage-dependent inactivation

Next, we illustrate voltage-dependent inactivation (VDI) using simulations with a protocol adapted from Magyar et al. (2000), shown in Figure 9 (left). As with activation, we show predictions for ideal voltage clamp and perfectly buffered ionic concentrations leading to a fixed but model-dependent level of CDI, so that there are important caveats when comparing these simulations to experimental data. Each sweep in the protocol starts with 20 s at a holding potential of —90 mV, followed by a 500 ms step (P1) to a variable *conditioning potential* and a 120 ms step (P2) to a fixed *test potential* of 0 mV. The conditioning potential at P1 is –90 mV for the first sweep and increases by 2 mV per sweep up to 30 mV for the final sweep (a subset of which is shown in the figure). Current was recorded during P2 to measure peak current (I, Figure 9, centre), which was normalised 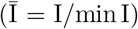 and plotted against conditioning potential to construct the I-V curves shown in Figure 9 (right).

**Figure 9:**
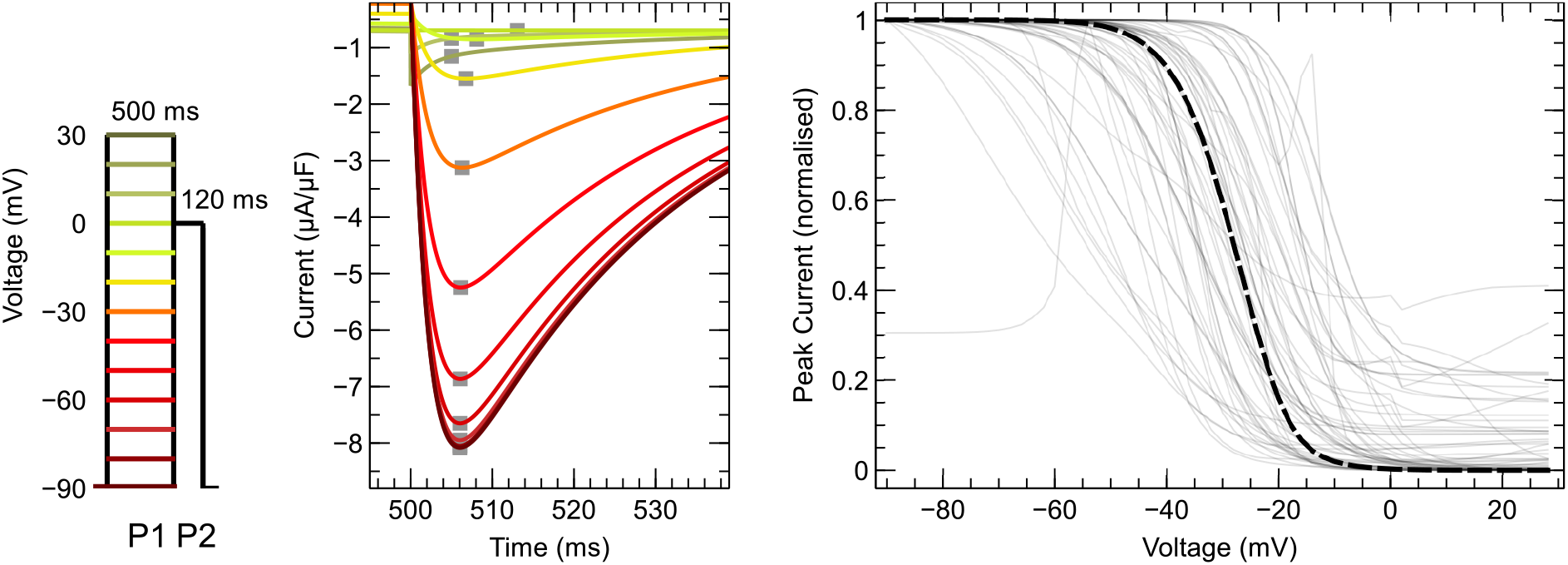
**Left**: Voltage-clamp protocol used to characterise voltage-dependent inactivation (see main text for details). **Centre**: Current response during P2 as predicted by the Grandi et al. (2010) model, with constant ionic concentrations and a fixed level of CDI. Traces from successive sweeps are overlaid and colour-coded to match the P1 steps in the left panel. The peak I_CaL_ at each voltage step (marked with grey squares) is used to construct the I-V curves on the right, as shown in an animation online at https://github.com/CardiacModelling/ical-model-review. **Right**: Normalised peak I_CaL_ versus step voltage curve for 60 models. The I-V curve from Kurata et al. (2002), which is close to the median response over all models, is highlighted with a black dashed line. An interactive view of these results is available at https://chaste.cs.ox.ac.uk/q/2022/ical/fig9.

This figure shows that most models predict inactivation to set in at around –60 mV or greater. The conditioning potential after which the peak I_CaL_ is halved (*V*_0.5_) varied between –59.3 to +10 mV for the models in this study while the median *V*_0.5_ was –29.9 mV. By comparison, examples from ventricular cells at physiological temperature show LCCs beginning to inactivate after a conditioning voltage of around –60mV, with *V*_0.5_ lying between –30 and –24 mV (Li et al., 1999; Kim et al., 2016).

#### 3.4.1 Time-course of recovery

Model predictions of the time I_CaL_ takes to recover from inactivation are shown using a protocol adapted from Fülöp et al. (2004) (Figure 10, left). Each sweep in this protocol starts with 10 s at a holding potential of −90 mV, followed by a 500 ms step (P1) to a conditioning potential of 0 mV. The potential is then set back to −90 mV for a duration that varies from sweep to sweep, followed by a 120 ms second step (P2) to 0 mV. The interval duration is 10 ms for the first sweep and increases by 10 ms per sweep up to 50 ms, after which it increases by 25 ms per sweep up to 1000 ms (a subset of which are shown in the figure).

**Figure 10:**
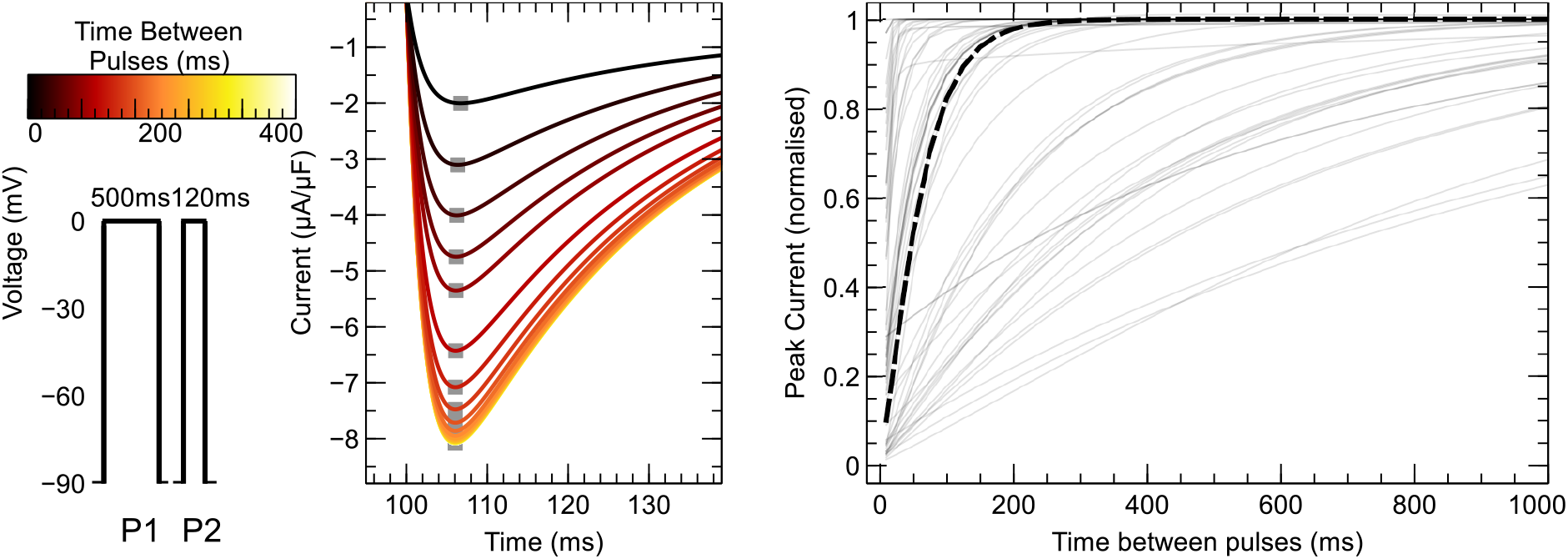
**Left**: Voltage-clamp protocol to characterise recovery from inactivation, with a variable time interval between P1 and P2 of 10, 20, 30, 40, 50 ms and increment steps of 25 ms thereafter until 1000 ms (see main text for details). **Centre**: Example of I_CaL_ predictions (constant [Ca^2+^]) during P2, from the model by Grandi et al. (2010). The colours of each line match the colour shown above the interval length in the left panel. Traces from successive sweeps are overlaid and the peak I_CaL_ is marked at each sweep. This is then used to construct the curves on the right, as shown in an animation online at https://github.com/CardiacModelling/ical-model-review. **Right**: Normalised peak I_CaL_ versus time curves for all 60 models. The prediction from Fink et al. (2008) is close to the median response across all models and is highlighted with a black dashed line. An interactive view of these results is available at https://chaste.cs.ox.ac.uk/q/2022/ical/fig10.

Peak I_CaL_ was recorded during P1 (I_conditioning_) and P2 (I_test_) for all sweeps (Figure 10, centre). This was normalised as 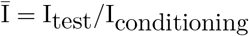 and the normalised peak 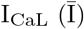 was plotted against interval length to construct the current-time curves shown in figure 10 (right).

An exponential function 1 – exp(–*t/τ*) was fitted to each model’s current-time curve to yield the time constant of recovery from inactivation *τ*. This *τ* varies between 1.4 to 995.7 ms for the models in this study while the median *τ* is 49.2 ms. This is much faster than examples from ventricular cells at physiological temperature, in which *τ* ranged from 50 to 700 ms (Li et al., 1999; Kim et al., 2016).

### 3.5 Calcium-dependent inactivation

To study the effects of calcium-dependent inactivation (CDI), we repeated the inactivation protocol of Figure 9 at constant internal (bulk, dyadic, and submembrane) [Ca^2+^] of 1 pM 0.1 μM, and 300 μM. The lowest value represents a situation with near to no Ca^2+^, while the other two values represent the normal range of calcium concentrations in the dyadic space during an AP (Fearnley et al., 2011). Post-processing proceeded similarly to before: peak I_CaL_ was identified at each sweep, but normalisation was performed using the maximum peak I_CaL_ measured at 1 pM, so that the magnitude of the low, moderate, and high [Ca^2+^] curves can be compared.

Figure 11 shows the resulting I-V curves for the three experimental conditions. For ease of presentation, results have been split into three categories: insensitive or weakly sensitive to [Ca^2+^]; mildly sensitive; and strongly sensitive to [Ca^2+^]. To assign a model to a sensitivity class, we defined a ‘root mean squared difference’ measure between the I-V curves for extreme concentrations as

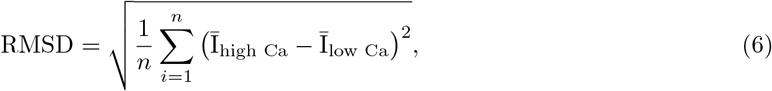

where *n* is the number of points in each I-V curve. We classified a model as weakly sensitive if RMSD < 0.1 (9 models), mildly sensitive if 0.1 ≤ RMSD ≤ 0.5 (17 models), and strongly sensitive if RMSD > 0.5 (34 models). Of the 9 models in the lowest sensitivity category, only Noble et al. (1991) and Pandit et al. (2001) account for [Ca^2+^] in their equations at all; the remaining 7 have no [Ca^2+^] terms in either O or *δ* (see Table 1, Row 1).

**Figure 11:**
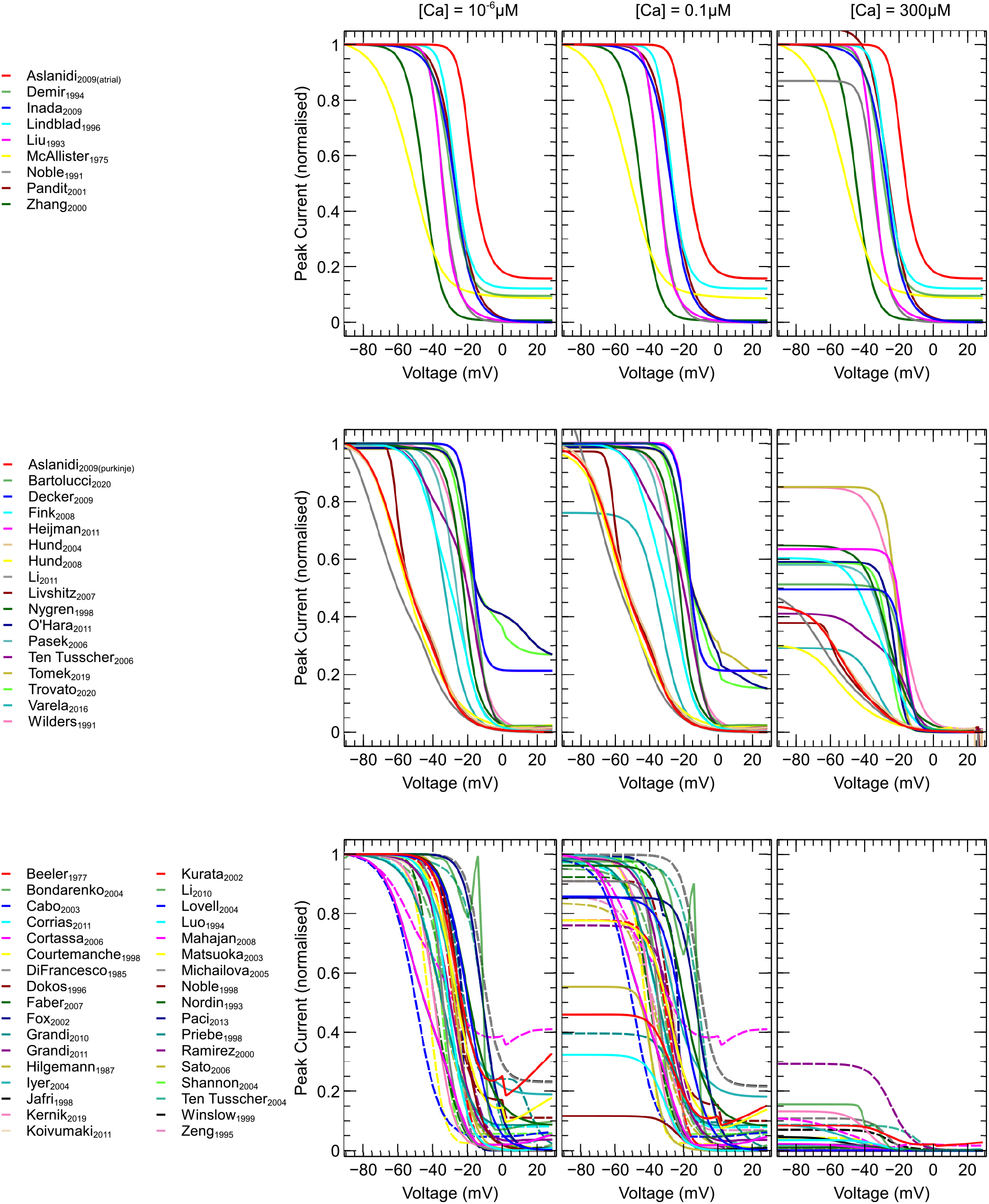
The inactivation protocol of Figure 9, for an internal [Ca^2+^] of 1pM (**Left**), 0.1 μM (**Centre**), and 300μM (**Right**). Results are grouped into three categories based on their sensitivity to [Ca^2+^]: insensitive or weakly sensitive (**Top**, 9 models); mildly sensitive (**Middle**, 17 models); and strongly sensitive (**Bottom**, 34 models). An interactive view of these results is available at https://chaste.cs.ox.ac.uk/q/2O22/ical/figll.

#### 3.5.1 [Ca^2+^] sensitivity and localisation

To further investigate the differences in predicted sensitivity to [Ca^2+^] we used a simple protocol consisting of a –90 mV holding potential and a single brief step to 0mV (i.e. the first sweep shown in Figure 9). Starting from an internal (bulk, dyadic, and submembrane) [Ca^2+^] of 1 pM (i.e. almost no Ca^2+^), we repeated the experiments with increasing concentrations until a 50% drop in the peak I_CaL_ was observed. Note that each repeat was started from the same initial conditions: no build-up of CDI (or CDF) occurred between repeats. We termed the resulting concentration the *CDI_50_* [Ca^2+^].

Using this procedure, the CDI_50_ [Ca^2+^] was obtained for 49 out of the 51 mildly and strongly [Ca^2+^] sensitive models, as shown in Figure 12. The models by Wilders et al. (1991) and Nygren et al. (1998) did not produce a 50% drop at any of the levels we tested. The calculated CDI_50_ is shown in Figure 12 (left) and varies across different models from 0.16 μM (10^th^ percentile) to 338 μM (90^th^ percentile) while the median CDI50 is 83.6 μM.

**Figure 12:**
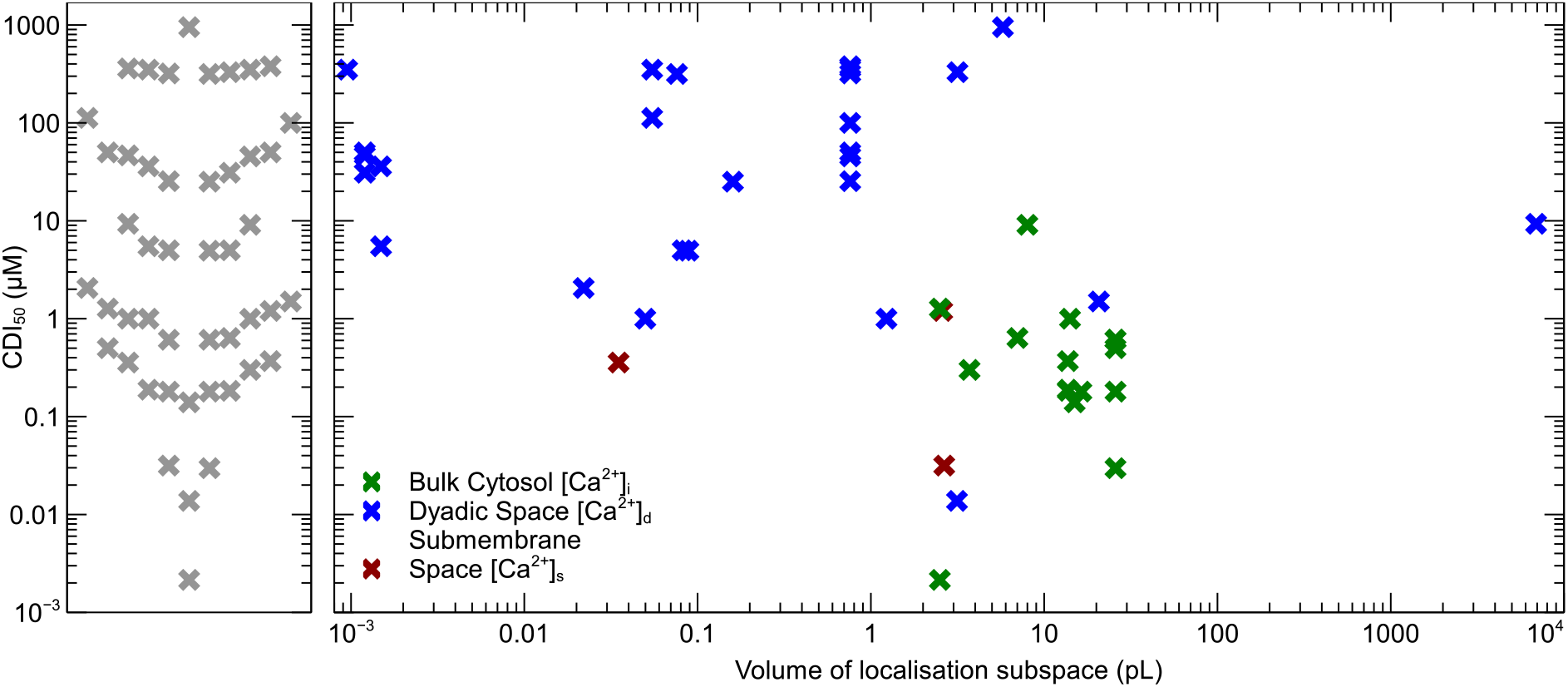
**Left**: CDI_50_ for 47 out of the 49 mildly or strongly [Ca^2+^] sensitive models (see main text for details). **Right**: CDI_50_ plotted against the volume of the subspace assumed to contain the LCCs, for the 47 models. Colours indicate the subspace containing the channels, or the subspace containing the majority of channels in situations where LCCs are assigned to multiple spaces.

In a more realistic situation, where [Ca^2+^] is not held constant throughout the cell, Ca^2+^ flowing into a subspace will induce a change in local [Ca^2+^] with a rate that depends inversely on the subspace volume *V*_local_ (assuming there is no buffering or diffusion of Ca^2+^). We may therefore expect models in which I_CaL_ flows into large subspaces to compensate for the slower local [Ca^2+^]-change with a more sensitive CDI, and vice versa. This is explored in Figure 12 (right), in which the CDI_50_ concentration is plotted against *V*_local_. For models in which LCCs are present in multiple biological subspaces, the volume was calculated as a weighted average: For example, 80% of the LCCs in the model by Tomek et al. (2019) lie in the dyadic space, whereas only 20% lie in the submembrane space, leading to a weighted average volume *V*_local_ = 0.8*V_d_* + 0.2*V_s_*. The models by Beeler and Reuter (1977) and Mahajan et al. (2008b) were omitted as the volume of their localisation space could not be determined from the manuscripts.

Figure 12 (right) shows a weak correlation between the predicted CDI_50_ and the corresponding *V*_local_, suggesting that modelled CDI is to some limited extent influenced by the size of the compartment that I_CaL_ flows into. This is another example where we observe that one aspect of the model can compensate for another.

### 3.6 AP and AP-CaT clamp predictions

The experiments simulated so far use voltage-step protocols which bring out a wide range of I_CaL_ kinetics but are very different from the voltage signals experienced by LCCs *in situ*. The final aspect we review is the models’ prediction of the I_CaL_ current elicited by a predefined AP. As before, we simulate an idealised voltage-clamp, but this time we allow internal [Ca^2+^] to vary. Because models including CDI differ in their geometrical layout, there is no obvious ‘best’ way to do this. Assumptions about internal calcium dynamics will impact the (estimated or calibrated) parameters governing I_CaL_ kinetics, and to bring these to the foreground we may choose to let the internal calcium concentrations in these simulations vary as defined by the model equations. If, however, we are interested only in the models’ I_CaL_ equations, it is preferable to force an identical CaT at the internal channel mouth in each simulation.

To address these conflicting aims, we performed a series of three complementary simulations. First, we simulated an AP-clamp in which we allowed the internal calcium concentration(s) to vary according to model equations (**AP clamp 1**). Secondly, we performed a dual AP-CaT-clamp, in which all internal [Ca^2+^] (including the subspaces) were clamped to the same pre-calculated bulk cytosolic CaT (**AP clamp 2**). Thirdly, we performed a combined AP-CaT-clamp in which bulk cytosol, submembrane space, and dyadic space [Ca^2+^] were clamped to pre-calculated transients appropriate to the subspace (**AP clamp 3**). We chose to use the AP and CaT measurements from the Grandi et al. (2010) human ventricle model during 1 Hz at steady pacing of the model (Figure 13, 3), as this is the earliest human model in which all three subspaces are defined. (Note that these strategies are meant to illustrate model characteristics, and are not necessarily experimentally feasible.) Because AP clamp 1 requires an AP model, this simulation was only performed for the 44 models for which a CellML file for the full AP model was available. For AP models with epi-, endo-, and mid-myocardial variants, the epicardial version was used. AP clamp 2 and 3 were performed for all 60 models. In all three simulations, I_CaL_ was normalised with respect to the total charge carried: 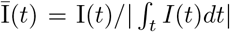. The resulting normalised currents are shown in Figure 14.

**Figure 13:**
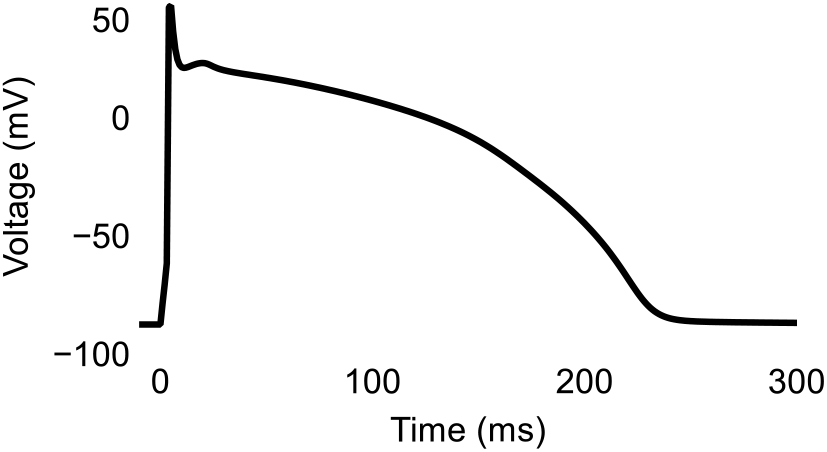
The AP waveform used in AP- and AP-CaT-clamp simulations in Figures 14–16. This action potential was derived from a 1 Hz simulation of the model by Grandi et al. (2010).

**Figure 14:**
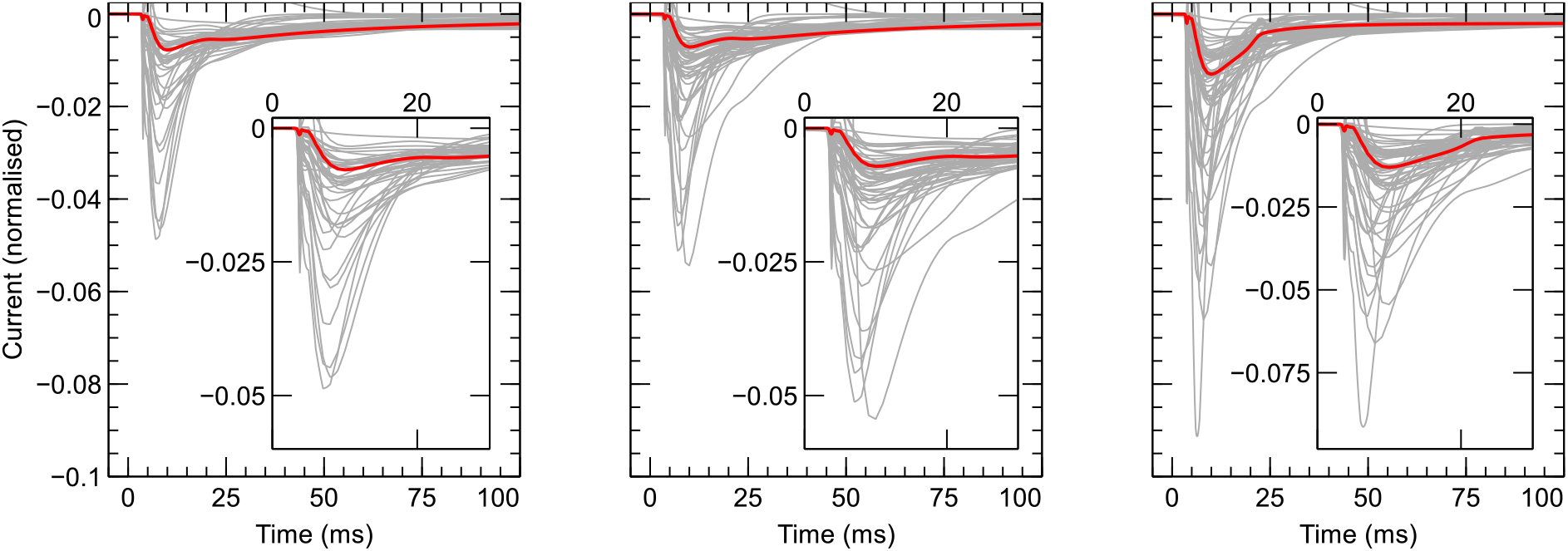
Variation in the I_CaL_ predictions in response to AP and AP-CaT clamp. Currents are shown normalised to the total charge carried (see text). **Left**: Simulated response to AP clamp 1, in which internal [Ca^2+^] are determined by each model’s equations. **Centre**: Simulated response to AP clamp 2, in which all internal [Ca^2+^] are clamped to an identical predetermined bulk-cytosolic CaT. **Right**: Simulated response to AP clamp 3, in which [Ca^2+^]_*i*_, [Ca^2+^]_s_, and [Ca^2+^]_*d*_ are clamped to the bulk cytosolic, submembrane, and dyadic CaT from the model by Grandi et al. (2010). Results for AP clamp 1 are limited to 44 cases where an AP model CellML file was available, whereas all 60 models are shown for the remaining AP clamps. Predictions from Grandi et al. (2010) are highlighted in red. An interactive view of these results is available at https://chaste.cs.ox.ac.uk/q/2022/ical/fig14.

#### 3.6.1 Variation in gating across models is not easily explained

All three AP clamps in Figure 14 show considerable variation in predicted (normalised) I_CaL_ between models. Some of this variation may be explained as a result of differences in e.g. species and cell-type, or more mundane factors such as the year of publication. To test this assumption, we divided the 60 models into groups according to several criteria and investigated whether this resulted in groups with reduced variation. We chose the ‘compromise’ protocol AP Clamp 3 for this purpose. To remove effects of varying driving terms and maximal conductances, we compared the predicted open probability (*O*) rather than I_CaL_. To give an indication of the variation within a group, we used the ‘normalised root mean square difference’ (NRMSD) of the observed open probabilities of all models in the group, defined as:

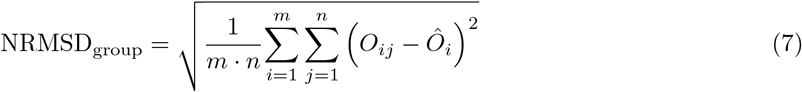

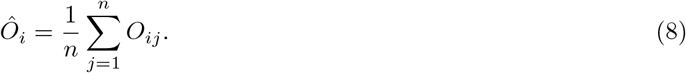

Here, *O_ij_* represents the open probability measured in response to the ‘AP-CaT clamp 3’ protocol at the *i^th^* time step for the *j^th^* model, *m* is the total number of time points, and *n* is the total number of models within the group.

If grouping by a particular category reduced variability, we would expect the NRMSD within each group to be lower than the NRMSD of the original group-of-all-models (NRMSD_total_ = 0.139). This can be quantified by defining a relative measure NRMSD_relative_ = NRMSD_group_/NRMSD_total_. A grouping can be considered to be better than baseline if its NRMSDrelative is less than one.

First, we investigated the effects of grouping models chronologically by grouping each model into one of four categories depending on the year of publication. This is shown in Figure 15, where we can observe some convergence in model predictions over time, indicated by a decreasing NRMSD_relative_. Although this is encouraging, some degree of convergence should be expected as more recent models are often refinements of previous models in the same age category. Large variability should also expected from the models published before the 1990s, which depend on data obtained from a wider range of experimental techniques.

**Figure 15:**
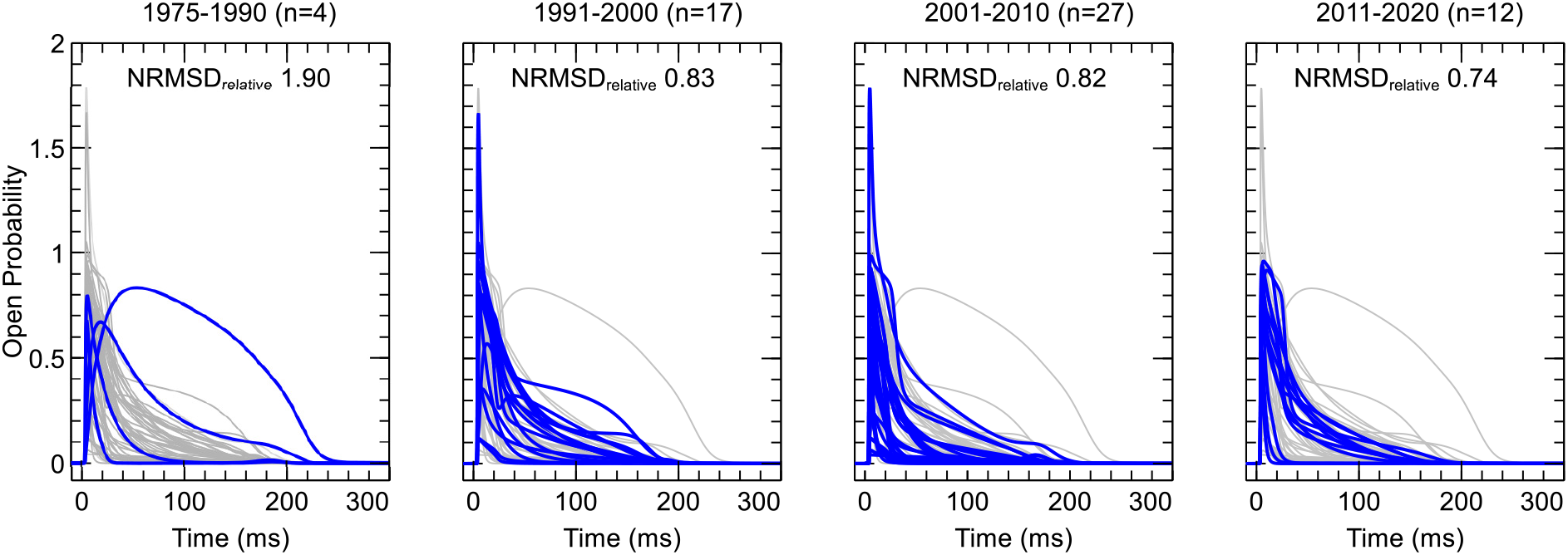
Open probability in response to AP clamp 3, grouped according to date of publication, shows a degree of convergence over time. Blue curves in each panel correspond to I_CaL_ predictions of models within the corresponding period, while the remaining predictions are shown in grey. (Note how some models allow *O* to be greater than 1, so that it is not strictly a probability in these models.)

Next, we grouped models according to the approach used to model how the calcium concentrations affect the open probability, depicted by the rows in Table 1. On this basis, Figure 16 shows the models grouped into 6 classes: no [Ca^2+^] (row 1, 2), [Ca^2+^]_*i*_ (row 3, 6), [Ca^2+^]_*s*_ (row 4, 7), [Ca^2+^]_*d*_ (row 5, 8, 11, 12), [Ca^2+^]_*i*_ and [Ca^2+^]_*d*_ (row 9), and [Ca^2+^]_*s*_ and [Ca^2+^]_*d*_ (row 10). It shows that that most groups formed on the basis of the local [Ca^2+^] near the LCC have at least ~20% less disagreement amongst the predictions across models than the baseline. This indicates that the modelled localisation of the LCCs plays an important role in determining model predictions. Note that class 5 contains only a single model (Tomek et al., 2019) so that no NRMSD_relative_ can be calculated. Similarly, all three models in class 6 (Shannon et al., 2004; Grandi et al., 2010, 2011) were developed by the same research group, which explains the very low NRMSD in this group.

**Figure 16:**
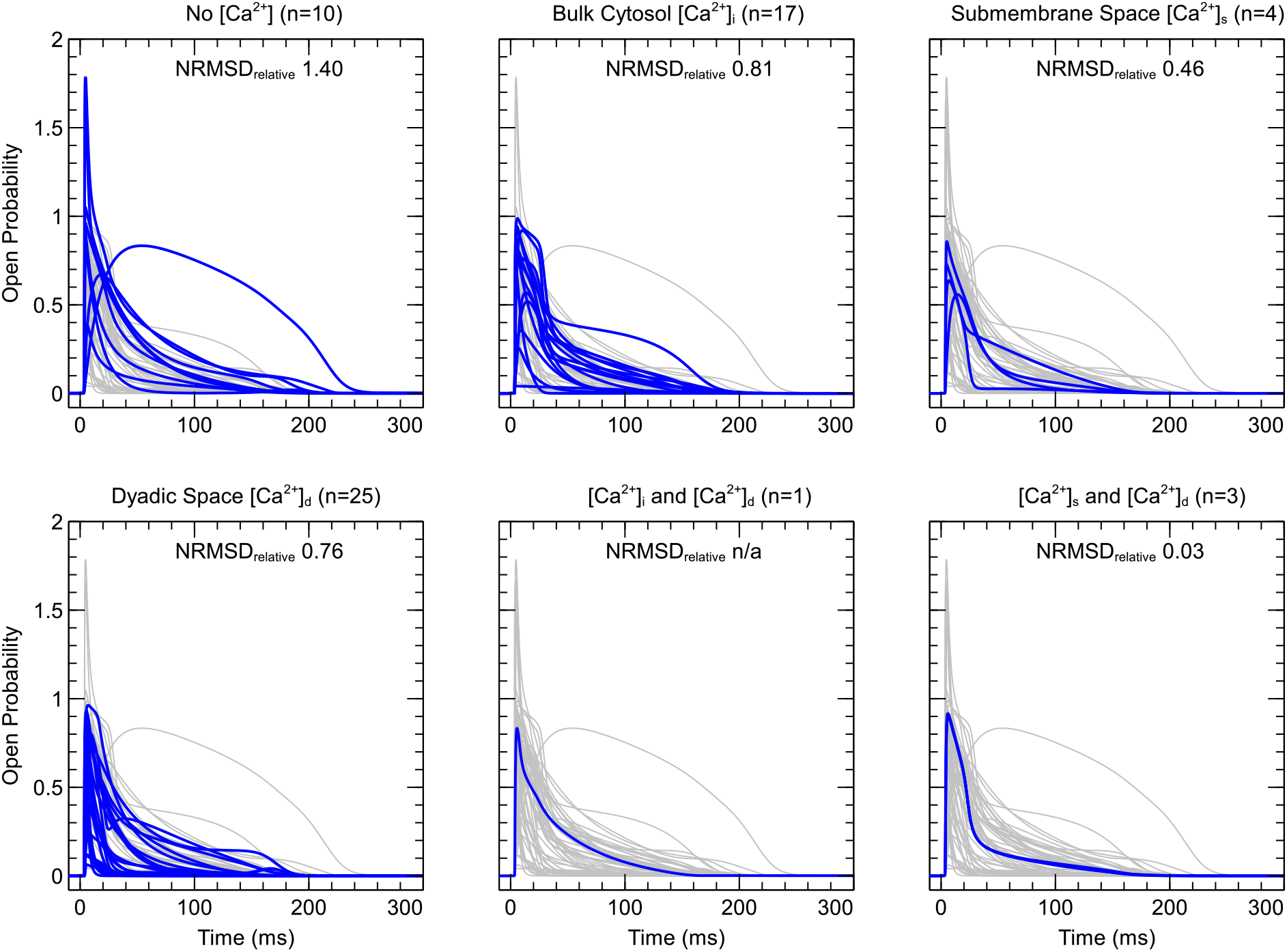
Open probability (*O*) in response to AP Clamp 3, grouped by local [Ca^2+^] as shown by the rows in Table 1 where each row represents an O or driving term dependence on different local [Ca^2+^]. The first panel ‘No [Ca^2+^]’ shows the models from rows 1 and 2, followed by ‘Bulk cytosol [Ca^2+^]’ (rows 3 and 6); ‘Submembrane space [Ca^2+^]’ (rows 4 and 7); ‘Dyadic space [Ca^2+^]’ (rows 5, 8, 11, and 12); ‘[Ca^2+^]_*i*_ and [Ca^2+^]_*d*_’ (row 9); and ‘[Ca^2+^]_*s*_ and [Ca^2+^]_*d*_’ (row 10). In each panel, blue curves show I_CaL_ predictions from models within the corresponding group, while the remaining predictions are shown in grey.

Finally, we investigated grouping the models by species and cell type. NRMSD_relative_ were obtained for groups of human (0.82, n=14), canine (0.62, n=14), rabbit (0.9, n=14), and guinea pig (0.83, n=8) models. Grouping by cell type we calculated NRMSD_relative_ for atrial (0.94, n=10), ventricular (1.05, n=35), sino-atrial (0.87, n=6), and Purkinje cells (0.83, n=6). (Examples of I_CaL_ in response to AP waveforms from an epicardial ventricular, endocardial ventricular, and atrial model can be viewed online at https://chaste.cs.ox.ac.uk/q/2022/ical/more_ap). Most groupings showed a small improvement on the baseline, but the only major reduction in NMRSD_relative_ was seen in the group of canine models.

Of all tested criteria, grouping by local [Ca^2+^] near the LCCs exhibited the most consistent predictions.

## 4 What next? Open questions and challenges

In the previous sections of this review we identified 73 models of mammalian I_CaL_, mapped their historical relationships, and analysed the qualitative differences in their constituent parts. Driving force models differed in their choice of Ohmic, GHK, or modified GHK equations as well as their choice of charge carrier(s). Fifteen major gating schemes were identified, with a further thirteen subtypes bringing the total to twenty-eight. Modelling of LCC localisation was much less diverse, with nearly all recent (native myocyte) models using a cell layout where LCCs emerge into a dyadic subspace. Using simulation, we reviewed the predictions of the current from 60 models using three common voltage-clamp protocols, as well as more complex AP and AP-CaT-clamp protocols that illustrated CDI. Great variability was observed between model predictions, which was somewhat diminished when grouping the models by their assumptions about LCC localisation, but no grouping could be found that significantly reduced (or explained) this variability. On several occasions, we observed how different parts of an I_CaL_ model could compensate for each other: unusual driving terms can be cancelled out by unusual gating, and modelled calcium-sensitivity in CDI correlated with the volume of the subspace LCCs were assumed to occupy.

These 73 models, along with the further models we could not cover in this review, represent a huge effort and a great achievement by the cell electrophysiology community. It is perhaps surprising, however, that no consensus or even ‘gold-standard’ model has emerged for I_CaL_, or even subcomponents *δ* and O. This situation presents a challenge for future modellers who need to choose an I_CaL_ model for their studies, or may even wish to add a 74^th^ model. In this section of the review we will focus on two open problems: 1. How can we select the best I_CaL_ model for any particular study — and how can we make this task easier in the future? 2. What new experimental or analytical work is needed to help us choose between different models of the electrochemical driving force and (voltage and calcium-dependent) gating?

### 4.1 Choosing I_CaL_ models

A popular quote from statistician George Box runs ‘all models are wrong, but some are useful’ (Box, 1979). This has occasionally been applied to models of electrophysiology, and indeed if we consider the size of an LCC alpha subunit (~170kDa), let alone of the full macromolecular complex formed by an LCC and its various subunits and regulators, it is easy to see how a 2, 8 or even 24-state model could never hope to capture its full complexity. Instead, Box (1976) argues, we should (1) apply Occam’s razor to select the simplest, most *parsimonious*, model that can accommodate the relevant data, and (2) choose a model that is wrong in ways that are both known and acceptable. Note that these criteria depend on the model’s intended *context of use* (Pathmanathan et al., 2017; Whittaker et al., 2020), so that there is room for more than one ‘best’ I_CaL_ model in cell electrophysiology.

A good example of this philosophy is the Ohmic driving term, which is used in many models of ionic currents — despite the common knowledge that the GHK equation is more accurate, and that this in turn is a clear oversimplification of the complex process of ion permeation (see e.g. Roux et al., 2004). For many currents, the Ohmic and GHK models both provide a good enough fit in the situations of interest and so the Ohmic model can be chosen based on its simplicity. The model’s limitations (e.g. poor predictions at extreme ‘non-physiological’ voltages or ionic concentrations) are known to the modeller and understood to be acceptable within the model’s intended context of use.

Can we use these principles to choose between our 73 models? If our goal is to match a particular data set, a reasonably straightforward (but very time consuming) approach could be to simply try fitting each gating scheme and each driving term model, choosing the simplest combination that fits, and then verifying its predictive power on a separate data set (which should tell us something about its limitations). Sections 2.4 and 2.5 and the CellML code supplied with this review provide an excellent starting point for such studies.

If our goal is to pick an already parameterised model from the literature, it is much more difficult to see if these principles apply. I_CaL_ model development is relatively well described in the literature: AP modelling papers like Luo and Rudy (1994), Jafri et al. (1998), and Mahajan et al. (2008b) devote several paragraphs and figures to it. In some cases, there is evidence of model complexity increasing when new biophysical detail was added, e.g. the addition of new gates in gating schemes A-C-L, or A-B-F-O. But it is interesting to see that many of the gating schemes in Figure 5 represent the same biophysical ideas of mode-switching (Hess et al., 1984) or the removal and application of a ‘brake’ (Pitt et al., 2001). We also found very little evidence of existing I_CaL_ models being tried and found inadequate before new ones were introduced (a notable exception being Luo and Rudy, 1994). It is unclear if such comparisons were not performed, e.g. due to unavailability of previous model code and data sets, or if they were simply not published. Methods for model selection have been proposed, and applied in ion channel modelling as early as Horn and Vandenberg (1984). Recent progress includes enumerating, fitting, and comparing a large number of possible MM structures (Mangold et al., 2021), and such model selection techniques should be explored more widely in this field (Whittaker et al., 2020). Limitations of I_CaL_ models are better documented, although these are usually given exclusively in terms of potential detail that could be added in future work.

Looking forward, there are several things we can do to facilitate future model comparison and selection. The field of cardiac cell electrophysiology has an unusually long history of sharing model and simulation code, dating back at least to the 1980s (Noble et al., 2012), and considerable effort has been expended to retroactively encode and share models in CellML (Yu et al., 2011). There is some distance to go yet — just over half the models in this review could be downloaded and used without corrections — but model code sharing on the whole appears to be getting both easier and more common.

Data sharing has a more limited history. Although the need for sharing data (and sufficient meta data) is widely acknowledged (Quinn et al., 2011) and technical barriers have all but disappeared in the last decade, it is still uncommon for electrophysiological recordings to be published in full digital form. As a result, modellers have frequently resorted to digitising published figures and then fitting to ‘summary curves’ (e.g. I-V curves) that represent averaged data from several cells in a condensed form. Both averaging (Golowasch et al., 2002) and fitting to summary curves (Clerx et al., 2019) can lead to inaccuracies, so there is an evident need for more widespread sharing of experimental data, perhaps in a systematic form such as the Physionet database for ECG signals (Goldberger et al., 2000). This situation will need to be drastically improved before we can answer simple questions like ‘which model best fits the currently known data’ (for a particular species and cell type), or set up public ‘benchmarks’ that models can be tested against, e.g. using the Cardiac Electrophysiology Web Lab (Cooper et al., 2016; Daly et al., 2018).

In the papers we reviewed, there is a notable shift in focus from the model itself (with new insights gained from the model in a short section at the end of the paper) to the model application and clinical/biophysical interpretation of its predictions. This can result in vital information about the model or data being moved into (less thoroughly examined) supplements, or even omitted. An interesting new approach to making models and data useful beyond a single paper, is to supply a detailed description (but also comparisons to previous models and more information about model limitations) in a *secondary publication* e.g. in The Physiome Journal for models and in e.g. Scientific Data, F1000Research, or PLOS ONE for data.

### 4.2 ChoosingI_CaL_ model parts

A major difficulty when comparing I_CaL_ models is that, although they share a common structure, their driving term (*δ*), voltage-dependent kinetics, and calcium sensitivity interact in ways that allow irregularities in one aspect to be compensated in another. For example the unusual open probability (*O*) of the model by Priebe and Beuckelmann (1998) (Figure 8) was shown to be compensated by a large reversal potential in its driving term (Figure 6) to give a typical I_CaL_ (https://chaste.cs.ox.ac.uk/q/2022/ical/priebe). Such compensation allows the model to predict a current similar to other models for some protocols (Figure 17, middle) but different current at other protocols (Figure 17, right). Similarly, the models by Demir et al. (1994), Lindblad et al. (1996), and Aslanidi et al. (2009a) allow O to be greater than one (so that the usual interpretation of open probability does not apply), which can be compensated for in either *δ* or a reduced maximal conductance. The model by Beeler and Reuter (1970) has a strong sensitivity to [Ca^2+^] (Figure 11) but does not include CDI, deriving its [Ca^2+^]-dependence from an unusually strong driving force (Figure 6) instead.

**Figure 17:**
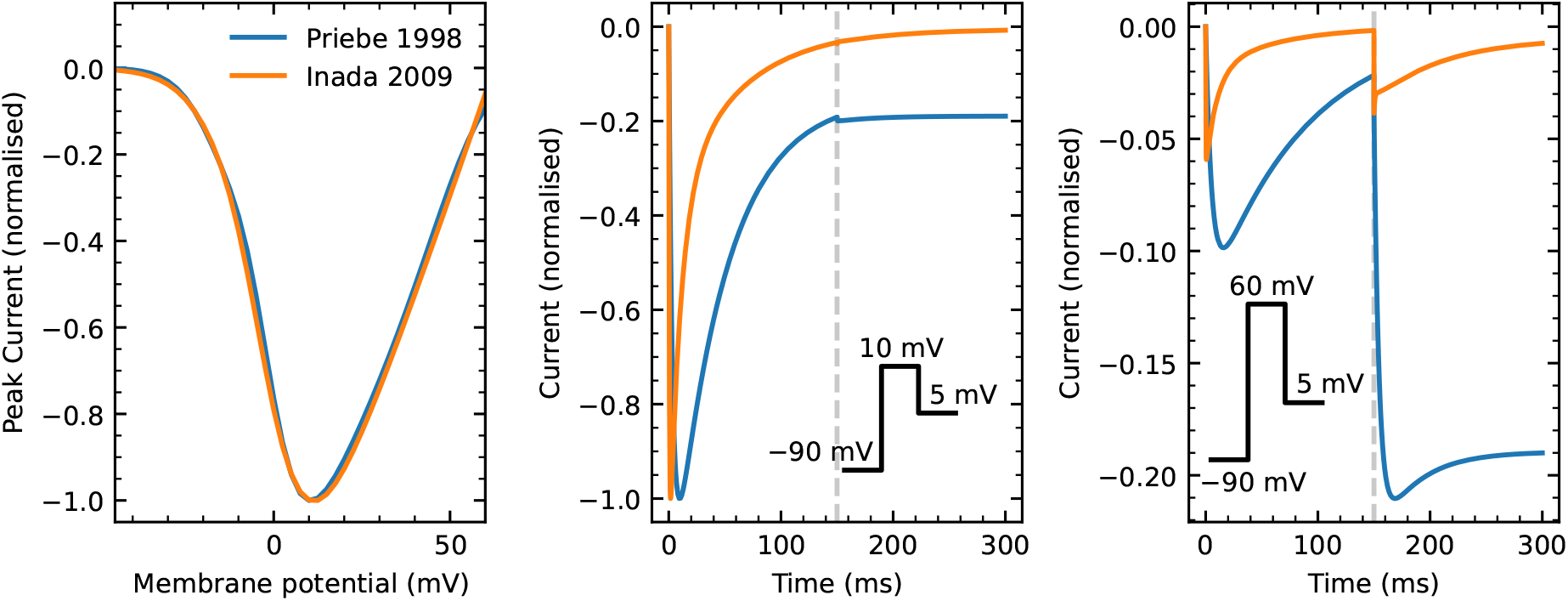
Compensation can hide model differences. **Left**: The IV curves for Inada et al. (2009) and Priebe and Beuckelmann (1998), normalised to the peak I_CaL_· These models were chosen because their IV curves almost overlap, suggesting similar activation characteristics. However, the model by Priebe & Beuckelmann is known to have an unusually low open probability at higher voltages (Figure 8) which is compensated by a stronger driving term (Figure 6). **Middle**: A step protocol showing quantitative, but not qualitative model differences. The membrane potential is held at −90mV (not shown), then stepped up to 10 mV (0 to 150 ms), and then down to 5 mV. Currents are shown using the same normalisation as before. In both models the step to 10 mV elicits the maximal response, followed by inactivation, which continues after the step to 5 mV. **Right**: A step to 60 mV elicits a much weaker response in both models (notice the scale on the y-axes). At this potential, we expect the models to fully activate and inactivate. However, the subsequent step to 5 mV reveals further activation in the model by Priebe & Beuckelmann, leading to an unexpected biphasic response, and a qualitative difference between the models’ activation characteristics.

Compensation also comes into play when creating or modifying AP models, which often require calibration of the sub-models (e.g. the I_CaL_ model) to cell-level observations (Whittaker et al., 2020). For example, Figure 12 shows how a high sensitivity to calcium can be compensated by postulating a bigger subspace near the LCCs, leading to a lower local [Ca^2+^]. More generally, an I_CaL_ model’s CDI gating may have to be tuned to the specific CaT in the AP model it was designed for (which suggests that some of the variability observed in Section 3.6 may be attributed to compensation for differences in CaT). Similarly, irregularities in an I_CaL_ model can, to a degree, be compensated by irregularities in the outward K^+^ currents and in the other components involved in CICR.

A model with subtly compensated defects may reproduce many electrophysiological phenomena seen in the experimental data, but through mechanisms that do not reflect those in the real cell. As a result, its predictions are likely to be unreliable when extrapolating to new, e.g. pathophysiological, situations. In this section, we once again break down an I_CaL_ model into its constituent parts, and discuss possible experiments that could let us study each in isolation.

#### 4.2.1 Driving term

The study of the electrochemical force driving I_CaL_ flux through the open channels is complicated by the wide range of calcium concentrations in a myocyte and the much higher concentrations of Na^+^ and K^+^ ions near the LCCs. Different models have been proposed as a result, starting with the Ohmic and GHK forms discussed in this review, but quickly moving on to more elaborate equations (Hess and Tsien, 1984; Roux et al., 2004) that (perhaps due to their computational cost) were never incorporated into AP models. This has left the question of how best to *approximate δ* in a computationally tractable I_CaL_ model unanswered, and none of the studies we surveyed showed a comparison of different driving-term models to motivate their choice.

To decide between driving force models, specialised experiments may be required that can separate driving force from kinetics and CDI. Single-channel ‘excised-patch’ experiments have shown promise in this respect (Hess et al., 1986; Greenstein and Winslow, 2002), and have the potential to allow rapid variation of internal (inside-out) or external calcium concentrations (outside-out), to levels that a whole cell may not tolerate, and in removing obfuscating effects from intracellular buffers and transporters (Gradogna et al., 2009). Single-channel results could then be scaled up to whole-cell level, with stochastic channel simulations used to check if a simple scaling approximation holds well enough when only small numbers of channels are open, or if a more complex method is needed.

Another interesting approach could be to fit Ohmic and/or GHK models to predictions from more sophisticated models of ionic permeation (Roux et al., 2004).

The same approaches could be used to try and resolve differences in relative permeability to different ionic species (Table 4). In the models reviewed in this study we found that the activity coefficients were highly similar between models. It is interesting to note that their values depend on experimental conditions and so could be calculated as part of the model (although within physiological ranges of temperature and ionic concentrations they change by less than 0.1% during an action potential at 1 Hz pacing).

#### 4.2.2 CDI and calcium dynamics

Calcium-dependent inactivation (CDI) is one of the most challenging aspects of modelling I_CaL_ and the degree of CDI predicted by different models varies widely (Figure 11). CDI depends on the (fluctuating) calcium concentration at the intracellular channel side, and so model assumptions about channel localisation and cell geometry have a strong impact on modelled CDI, as evidenced by the relationship between subspace volume and CDI50 in Figure 12 and by the reduction in variability between models when grouping by channel localisation.

Experimentally, CDI can be separated from voltage-dependent kinetics and driving term to some degree by using voltage protocols with a single repeating step, so that voltage effects are consistent — if still unknown. However, the local calcium concentration is difficult to control, as the very low [Ca^2+^] maintained in living cells means the calcium influx through I_CaL_ causes a significant change. In myocytes, a bewildering array of processes affect and are affected by local [Ca^2+^] (Bers, 2008). In heterologous expression systems, calcium-buffers are required to keep [Ca^2+^] at levels the cells can tolerate (You et al., 1997). Bulk cytosolic [Ca^2+^] can be measured optically using fluorescent dyes (Herron et al., 2012) but measuring local calcium elevations in subspaces (or ‘microdomains’) is technically challenging (Acsai et al., 2011). As a result, there is no simple set-up in which CDI can be studied, and analysing data from experiments targeting CDI requires consideration of intracellular [Ca^2+^] gradients, diffusion, and buffering. Although analytical solutions exist for simplified scenarios (Smith, 1996), it is likely that numerical simulations (e.g. Rice et al., 1999; Greenstein and Winslow, 2002; Koivumäki et al., 2011; Nivala et al., 2012) will be required for this purpose.

#### 4.2.3 Voltage-dependent kinetics

The equations for *O* vary more from model to model than those for any other aspect of I_CaL_ (Figure 5), and we found large variability in the predicted response to voltage-clamp protocols, which cannot be easily reduced by grouping models by e.g. cell type. While most studies of I_CaL_ kinetics have used conventional voltage-step protocols (such as shown in Figures 7, 9, and 10), recent advances in protocol design for I_Kr_ (Beattie et al., 2018; Clerx et al., 2019) may be translatable to I_CaL_. Application of such protocols in high-throughput settings (Lei et al., 2019) with careful consideration of experimental artefacts (Lei et al., 2020a,b) may help produce rich data sets that will allow us to distinguish between the different proposed gating mechanisms and study (biological or experimental) variability. Such an approach, however, would require that voltage and calcium effects are independent (or at least can be studied independently) and may rely on accurate CDI and driving term models already being available. Alternatively, it may be possible to derive gating models as simplifications of more detailed molecular dynamics models (Silva, 2018; Ramasubramanian and Rudy, 2018). In either case, care must be taken to create independent training and validation data sets, so that the most crucial part of the gating model, its predictive power, can be assessed (Whittaker et al., 2020).

### 4.3 Determining the impact of I_CaL_ model choice on electrophysiological properties

I_CaL_ is an important determinant of key electrophysiological properties at the cell, tissue, and organ level, including APD restitution (Guzman et al., 2010; Mahajan et al., 2008a), development of cellular, discordant, and T-wave alternans (Zhu and Clancy, 2007), EAD formation (Weiss et al., 2010; Heijman et al., 2014; Kettlewell et al., 2019) and cell-to-cell conduction (Joyner et al., 1996; Rohr and Kucera, 1997; Shaw and Rudy, 1997). It is therefore of interest to compare how the 73 models described in this study affect such emergent properties. However, a comprehensive study along these lines requires not just an isolated I_CaL_ model, but rather an I_CaL_ model in the context of a full AP model (or tissue model). An obvious way to start such a study would be to pick an AP model, replace its I_CaL_ formulation and test its predictions (repeating the process for all or a subset of the models in this review). This is a non-trivial exercise for several reasons, including: 1. The shape of the AP could be significantly affected, even after tuning the I_CaL_ conductance. 2. The CaT and detailed calcium dynamics could be similarly affected, and it may not be possible to set a new maximum conductance that simultaneously corrects the AP shape and CaT. 3. The I_CaL_ model may be incompatible with the CaT and AP that the cell model provides (e.g. it may not activate or inactivate to the right degree at the voltages and [Ca^2+^] that the AP model predicts). 4. Different assumptions about LCC localisation would need to be reconciled. 5. The new I_CaL_ model could predict different Na^+^ and K^+^ fluxes, which could affect the model’s long-term stability.

One way in which a preliminary study can be set up is by only changing the O from one I_CaL_ model to the other, and subsequently tuning the g_Ca_/P_Ca_ (along with P_Na_/P_K_ if applicable) so that the resultant AP and/or CaT most closely resemble their original shapes (thereby treating obstacles 1 and 2). Assuming this can be done, this leaves obstacles 3-5, but these might be avoided if a subset of ‘similar enough’ (but interestingly different) I_CaL_ models can be found. By changing the I_CaL_ kinetics through O, such a the study could then evaluate e.g. how changes in the time constant of recovery affect restitution (Mahajan et al., 2008b) or how changes in a model’s I_CaL_ ‘window-current’ affect the onset of EADs (Kettlewell et al., 2019). Note however, that such properties are not always directly encoded in a single model parameter, so that more complex multi-parameter adjustments may need to be made.

A final important complication arises since emergent cell, tissue, and organ properties depend on more than a single ‘molecular factor’ (Weiss et al., 2015); nonlinear interactions of I_CaL_ with other currents that are also modelled differently in various AP models will influence emergent behaviour. For instance, an I_CaL_ model ‘correctly’ producing alternans in one AP model may fail to do so in another (Hopenfeld, 2006). So any study inserting different I_CaL_ models into an AP model would also need to consider the influence of the AP model choice.

## 5 Conclusion

Five decades of careful research has resulted in an impressive list of 73 mammalian I_CaL_ models, 15 models of LCC gating, and a smaller but similar lack of consensus on how to model I_CaL_’s electrochemical driving force. Accordingly, the 60 models studied quantitatively made very different predictions about activation, inactivation, recovery, and CDI. However, as practical limitations on model sharing, data sharing, and extended publication forms are rapidly disappearing, the field is poised to take advantage of new technological possibilities (and publishing norms) that might allow these decades old questions to be systematically investigated and resolved. We believe that this review is the first to systematically survey and compare I_CaL_ models and hope that it serves as a starting point for a critical re-assessment of L-type calcium current modelling from which a synthesised, community consensus model may emerge.

## Acknowledgements

The authors thank Wayne R. Giles for his careful reading of an early draft of the manuscript and his helpful comments.

## References

Abriel, H., Rougier, J.-S. and Jalife, J. (2015) Ion channel macromolecular complexes in cardiomyocytes: roles in sudden cardiac death. Circulation Research, 116, 1971–1988.

Acsai, K., Antoons, G., Livshitz, L., Rudy, Y. and Sipido, K. R. (2011) Microdomain Ca^2+^near ryanodine receptors as reported by L-type Ca^2+^and Na^+^/Ca^2+^exchange currents. The Journal of Physiology, 589, 2569–2583.

Aggarwal, R. and Boyden, P. A. (1995) Diminished Ca^2+^and Ba^2+^currents in myocytes surviving in the epicardial border zone of the 5-day infarcted canine heart. Circulation Research, 77, 1180–1191.

Aggarwal, R. and Boyden, P. A. (1996) Altered pharmacologic responsiveness of reduced L-type calcium currents in myocytes surviving in the infarcted heart. Journal of Cardiovascular Electrophysiology, 7, 20–35.

Amin, A., Verkerk, A., Bhuiyan, Z., Wilde, A. and Tan, H. (2005) Novel Brugada syndrome-causing mutation in ion-conducting pore of cardiac Na^+^channel does not affect ion selectivity properties. Acta physiologica scandinavica, 185, 291–301.

Asakura, K., Cha, C. Y., Yamaoka, H., Horikawa, Y., Memida, H., Powell, T., Amano, A. and Noma, A. (2014) EAD and DAD mechanisms analyzed by developing a new human ventricular cell model. Progress in Biophysics and Molecular Biology, 116, 11–24.

Aslanidi, O. V., Boyett, M. R., Dobrzynski, H., Li, J. and Zhang, H. (2009a) Mechanisms of transition from normal to reentrant electrical activity in a model of rabbit atrial tissue: interaction of tissue heterogeneity and anisotropy. Biophysical Journal, 96, 798–817.

Aslanidi, O. V., Stewart, P., Boyett, M. R. and Zhang, H. (2009b) Optimal velocity and safety of discontinuous conduction through the heterogeneous Purkinje-ventricular junction. Biophysical Journal, 97, 20–39.

Bartolucci, C., Passini, E., Hyttinen, J., Paci, M. and Severi, S. (2020) Simulation of the effects of extracellular calcium changes leads to a novel computational model of human ventricular action potential with a revised calcium handling. Frontiers in Physiology, 11, 314.

Bassingthwaighte, J. B. and Reuter, H. (1972) Calcium movements and excitation-contraction coupling in cardiac cells. In Electrical Phenomena in the Heart (ed. W. C. De Mello), chap. 13, 266–290. Academic Press.

Beattie, K. A., Hill, A. P., Bardenet, R., Cui, Y., Vandenberg, J. I., Gavaghan, D. J., de Boer, T. P. and Mirams, G. R. (2018) Sinusoidal voltage protocols for rapid characterisation of ion channel kinetics. The Journal of Physiology, 596, 1813–1828.

Beeler, G. W. and Reuter, H. (1970) Membrane calcium current in ventricular myocardial fibres. The Journal of Physiology, 207, 191–209.

Beeler, G. W. and Reuter, H. (1977) Reconstruction of the action potential of ventricular myocardial fibres. The Journal of Physiology, 268, 177–210.

Bénitah, J.-P., Bailly, P., D’Agrosa, M.-C., Da Ponte, J.-P., Delgado, C. and Lorente, P. (1992) Slow inward current in single cells isolated from adult human ventricles. Pflügers Archiv-European Journal of Physiology, 421, 176–187.

Bers, D. M. (2008) Calcium cycling and signaling in cardiac myocytes. Annual Review of Physiology, 70, 23–49.

Bers, D. M. and Morotti, S. (2014) Ca^2+^current facilitation is CaMKII-dependent and has arrhythmogenic consequences. Frontiers in Pharmacology, 5, 144.

Beuckelmann, D. J., Näbauer, M. and Erdmann, E. (1991) Characteristics of calcium-current in isolated human ventricular myocytes from patients with terminal heart failure. Journal of Molecular and Cellular Cardiology, 23, 929–937.

Beuckelmann, D. J., Näbauer, M. and Erdmann, E. (1992) Intracellular calcium handling in isolated ventricular myocytes from patients with terminal heart failure. Circulation, 85, 1046–1055.

Bondarenko, V. E., Szigeti, G. P., Bett, G. C., Kim, S.-J. and Rasmusson, R. L. (2004) Computer model of action potential of mouse ventricular myocytes. American Journal of Physiology-Heart and Circulatory Physiology, 287, H1378–H1403.

Box, G. E. (1976) Science and statistics. Journal of the American Statistical Association, 71, 791–799.

Box, G. E. (1979) Robustness in the strategy of scientific model building. In Robustness in statistics, 201–236. Elsevier.

Boyett, M. R., Honjo, H., Harrison, S. M., Zang, W. J. and Kirby, M. S. (1994) Ultra-slow voltage-dependent inactivation of the calcium current in guinea-pig and ferret ventricular myocytes. Pflügers Archiv-European Journal of Physiology, 428, 39–50.

Cabo, C. and Boyden, P. A. (2003) Electrical remodeling of the epicardial border zone in the canine infarcted heart: a computational analysis. American Journal of Physiology-Heart and Circulatory Physiology, 284, H372–H384.

Campbell, D. L., Giles, W. R., Hume, J. R., Noble, D. and Shibata, E. F. (1988) Reversal potential of the calcium current in bull-frog atrial myocytes. The Journal of Physiology, 403, 267–286.

Catterall, W. A. (2011) Voltage-gated calcium channels. Cold Spring Harbor Perspectives in Biology, 3, a003947.

Cavalié, A., Pelzer, D. and Trautwein, W. (1986) Fast and slow gating behaviour of single calcium channels in cardiac cells. Pflügers Archiv-European Journal of Physiology, 406, 241–258.

Cens, T., Rousset, M., Leyris, J.-P., Fesquet, P. and Charnet, P. (2006) Voltage-and calcium-dependent inactivation in high voltage-gated Ca^2+^channels. Progress in biophysics and molecular biology, 90, 104–117.

Clerx, M., Beattie, K. A., Gavaghan, D. J. and Mirams, G. R. (2019) Four ways to fit an ion channel model. Biophysical Journal, 117, 2420–2437.

Clerx, M., Collins, P., de Lange, E. and Volders, P. G. A. (2016) Myokit: a simple interface to cardiac cellular electrophysiology. Progress in Biophysics and Molecular Biology, 120, 100–114.

Colman, M. A., Alvarez-Lacalle, E., Echebarria, B., Sato, D., Sutanto, H. and Heijman, J. (2022) Multiscale computational modeling of spatial calcium handling from nanodomain to whole-heart: Overview and perspectives. Frontiers in Physiology, 278.

Cooper, J., Mirams, G. R. and Niederer, S. A. (2011) High-throughput functional curation of cellular electrophysiology models. Progress in Biophysics and Molecular Biology, 107, 11–20.

Cooper, J., Scharm, M. and Mirams, G. R. (2016) The cardiac electrophysiology web lab. Biophysical Journal, 110, 292–300.

Corrias, A., Giles, W. and Rodriguez, B. (2011) Ionic mechanisms of electrophysiological properties and repolarization abnormalities in rabbit Purkinje fibers. American Journal of Physiology-Heart and Circulatory Physiology, 300, H1806–H1813.

Cortassa, S., Aon, M. A., O’Rourke, B., Jacques, R., Tseng, H.-J., Marbán, E. and Winslow, R. L. (2006) A computational model integrating electrophysiology, contraction, and mitochondrial bioenergetics in the ventricular myocyte. Biophysical Journal, 91, 1564–1589.

Courtemanche, M., Ramirez, R. J. and Nattel, S. (1998) Ionic mechanisms underlying human atrial action potential properties: insights from a mathematical model. American Journal of Physiology-Heart and Circulatory Physiology, 275, H301–H321.

Daly, A. C., Clerx, M., Beattie, K. A., Cooper, J., Gavaghan, D. J. and Mirams, G. R. (2018) Reproducible model development in the cardiac electrophysiology Web Lab. Progress in Biophysics and Molecular Biology, 139, 3–14.

Davies, C. W. and Malpass, V. E. (1964) Ion association and the viscosity of dilute electrolyte solutions. Transactions of the Faraday Society, 60, 2075–2084.

Decker, K. F., Heijman, J., Silva, J. R., Hund, T. J. and Rudy, Y. (2009) Properties and ionic mechanisms of action potential adaptation, restitution, and accommodation in canine epicardium. American Journal of Physiology-Heart and Circulatory Physiology, 296, H1017–H1026.

Demir, S. S., Clark, J. W. and Giles, W. R. (1999) Parasympathetic modulation of sinoatrial node pacemaker activity in rabbit heart: a unifying model. American Journal of Physiology-Heart and Circulatory Physiology, 276, H2221–H2244.

Demir, S. S., Clark, J. W., Murphey, C. R. and Giles, W. R. (1994) A mathematical model of a rabbit sinoatrial node cell. American Journal of Physiology-Cell Physiology, 266, C832–C852.

DiFrancesco, D. and Noble, D. (1985) A model of cardiac electrical activity incorporating ionic pumps and concentration changes. Philosophical Transactions of the Royal Society of London. B, Biological Sciences, 307, 353–398.

Dokos, S., Celler, B. and Lovell, N. (1996) Ion currents underlying sinoatrial node pacemaker activity: a new single cell mathematical model. Journal of Theoretical Biology, 181, 245–272.

Dolphin, A. C. (2016) Voltage-gated calcium channels and their auxiliary subunits: physiology and pathophysiology and pharmacology. The Journal of Physiology, 594, 5369–5390.

Egan, T. M., Noble, D., Noble, S. J., Powell, T., Spindler, A. J. and Twist, V. W. (1989) Sodium-calcium exchange during the action potential in guinea-pig ventricular cells. The Journal of Physiology, 411, 639–661.

Ehrlich, J. R., Cha, T.-J., Zhang, L., Chartier, D., Melnyk, P., Hohnloser, S. H. and Nattel, S. (2003) Cellular electrophysiology of canine pulmonary vein cardiomyocytes: action potential and ionic current properties. The Journal of Physiology, 551, 801–813.

Eisner, D. A., Caldwell, J. L., Kistamás, K. and Trafford, A. W. (2017) Calcium and excitation-contraction coupling in the heart. Circulation Research, 121, 181–195.

Escande, D., Coulombe, A., Faivre, J. F. and Coraboeuf, E. (1986) Characteristics of the time-dependent slow inward current in adult human atrial single myocytes. Journal of Molecular and Cellular Cardiology, 18, 547–551.

Faber, G. M. and Rudy, Y. (2000) Action potential and contractility changes in [Na^+^]_i_ overloaded cardiac myocytes: a simulation study. Biophysical Journal, 78, 2392–2404.

Faber, G. M., Silva, J., Livshitz, L. and Rudy, Y. (2007) Kinetic properties of the cardiac L-type Ca^2+^channel and its role in myocyte electrophysiology: a theoretical investigation. Biophysical Journal, 92, 1522–1543.

Fawcett, D. W. and McNutt, N. S. (1969) The ultrastructure of the cat myocardium: I. ventricular papillary muscle. The Journal of cell biology, 42, 1–45.

Fearnley, C. J., Roderick, H. L. and Bootman, M. D. (2011) Calcium signaling in cardiac myocytes. Cold Spring Harbor Perspectives in Biology, 3, a004242.

Fermini, B. and Nathan, R. D. (1991) Removal of sialic acid alters both T-and L-type calcium currents in cardiac myocytes. American Journal of Physiology-Heart and Circulatory Physiology, 260, H735–H743.

Ferreira, G., Ríos, E. and Reyes, N. (2003) Two components of voltage-dependent inactivation in Cav1.2 channels revealed by its gating currents. Biophysical journal, 84, 3662–3678.

Findlay, I. (2002) Voltage-and cation-dependent inactivation of L-type Ca^2+^channel currents in guineapig ventricular myocytes. The Journal of Physiology, 541, 731–740.

Findlay, I. (2004) Physiological modulation of inactivation in L-type Ca^2+^channels: one switch. The Journal of Physiology, 554, 275–283.

Fink, M., Noble, D., Virag, L., Varro, A. and Giles, W. R. (2008) Contributions of HERG K^+^current to repolarization of the human ventricular action potential. Progress in Biophysics and Molecular Biology, 96, 357–376.

Fox, J. J., McHarg, J. L. and Gilmour Jr, R. F. (2002) Ionic mechanism of electrical alternans. American Journal of Physiology-Heart and Circulatory Physiology, 282, H516–H530.

Fozzard, H. A. (2002) Cardiac sodium and calcium channels: a history of excitatory currents. Cardiovascular Research, 55, 1–8.

Frankenhaeuser, B. (1960) Sodium permeability in toad nerve and in squid nerve. The Journal of Physiology, 152, 159–166.

Friedman, N., Vinet, A. and Roberge, F. (1996) A study of a new model of the cardiac ventricular cell incorporating myoplasmic calcium regulation. In 22^nd^ Canadian Medical and Biological Engineering Society Conference, 92–93.

Fülöp, L., Bányász, T., Magyar, J., Szentandrássy, N., Varró, A. and Nánási, P. P. (2004) Reopening of L-type calcium channels in human ventricular myocytes during applied epicardial action potentials. Acta Physiologica Scandinavica, 180, 39–47.

Gaborit, N., Le Bouter, S., Szuts, V., Varro, A., Escande, D., Nattel, S. and Demolombe, S. (2007) Regional and tissue specific transcript signatures of ion channel genes in the non-diseased human heart. The Journal of Physiology, 582, 675–693.

Gao, T., Yatani, A., Dell’Acqua, M. L., Sako, H., Green, S. A., Dascal, N., Scott, J. D. and Hosey, M. M. (1997) cAMP-dependent regulation of cardiac L-type Ca^2+^channels requires membrane targeting of PKA and phosphorylation of channel subunits. Neuron, 19, 185–196.

Gettes, L. S. and Reuter, H. (1974) Slow recovery from inactivation of inward currents in mammalian myocardial fibres. The Journal of Physiology, 240, 703–724.

Goldberger, A. L., Amaral, L. A., Glass, L., Hausdorff, J. M., Ivanov, P. C., Mark, R. G., Mietus, J. E., Moody, G. B., Peng, C.-K. and Stanley, H. E. (2000) PhysioBank, PhysioToolkit, and PhysioNet: components of a new research resource for complex physiologic signals. Circulation, 101, e215–e220.

Goldman, D. E. (1943) Potential, impedance, and rectification in membranes. The Journal of General Physiology, 27, 37–60.

Golowasch, J., Goldman, M. S., Abbott, L. F. and Marder, E. (2002) Failure of averaging in the construction of a conductance-based neuron model. Journal of Neurophysiology, 87, 1129–1131.

Goonasekera, S. A., Hammer, K., Auger-Messier, M., Bodi, I., Chen, X., Zhang, H., Reiken, S., Elrod, J. W., Correll, R. N., York, A. J. et al. (2012) Decreased cardiac L-type Ca^2+^channel activity induces hypertrophy and heart failure in mice. The Journal of Clinical Investigation, 122, 280–290.

Gradogna, A., Scholz-Starke, J., Gutla, P. V. K. and Carpaneto, A. (2009) Fluorescence combined with excised patch: measuring calcium currents in plant cation channels. The Plant Journal, 58, 175–182.

Grandi, E., Pandit, S. V., Voigt, N., Workman, A. J., Dobrev, D., Jalife, J. and Bers, D. M. (2011) Human atrial action potential and Ca^2+^model: sinus rhythm and chronic atrial fibrillation. Circulation Research, 109, 1055–1066.

Grandi, E., Pasqualini, F. S. and Bers, D. M. (2010) A novel computational model of the human ventricular action potential and Ca transient. Journal of Molecular and Cellular Cardiology, 48, 112–121.

Greenstein, J. L., Hinch, R. and Winslow, R. L. (2006) Mechanisms of excitation-contraction coupling in an integrative model of the cardiac ventricular myocyte. Biophysical Journal, 90, 77–91.

Greenstein, J. L. and Winslow, R. L. (2002) An integrative model of the cardiac ventricular myocyte incorporating local control of Ca^2+^release. Biophysical Journal, 83, 2918–2945.

Guzman, K. M., Jing, L. and Patwardhan, A. (2010) Effects of changes in the l-type calcium current on hysteresis in restitution of action potential duration. Pacing and clinical electrophysiology, 33, 451–459.

Habuchi, Y., Noda, T., Nishimura, M. and Watanabe, Y. (1990) Recovery of the slow inward current from Ca^2+^-mediated and voltage-dependent inactivation in the rabbit sinoatrial node. Journal of Molecular and Cellular Cardiology, 22, 469–482.

Hadley, R. W. and Lederer, W. J. (1991) Ca^2+^and voltage inactivate Ca^2+^channels in guinea-pig ventricular myocytes through independent mechanisms. The Journal of Physiology, 444, 257–268.

Hagiwara, N., Irisawa, H. and Kameyama, M. (1988) Contribution of two types of calcium currents to the pacemaker potentials of rabbit sino-atrial node cells. The Journal of Physiology, 395, 233–253.

Han, W., Chartier, D., Li, D. and Nattel, S. (2001) Ionic remodeling of cardiac Purkinje cells by congestive heart failure. Circulation, 104, 2095–2100.

Harvey, R. D. and Hell, J. W. (2013) Ca_V_1.2 signaling complexes in the heart. Journal of Molecular and Cellular Cardiology, 58, 143–152.

Hashambhoy, Y. L., Winslow, R. L. and Greenstein, J. L. (2009) CaMKII-induced shift in modal gating explains L-type Ca^2+^current facilitation: a modeling study. Biophysical Journal, 96, 1770–1785.

Hedley, W. J., Nelson, M. R., Bellivant, D. P. and Nielsen, P. F. (2001) A short introduction to CellML. Philosophical Transactions of the Royal Society of London A: Mathematical, Physical and Engineering Sciences, 359, 1073–1089.

Heijman, J. (2012) Computational analysis of β-adrenergic stimulation and its effects on cardiac ventricular electrophysiology. Ph.D. thesis, Maastricht University.

Heijman, J., Voigt, N., Nattel, S. and Dobrev, D. (2014) Cellular and molecular electrophysiology of atrial fibrillation initiation, maintenance, and progression. Circulation Research, 114, 1483–1499.

Heijman, J., Volders, P. G. A., Westra, R. L. and Rudy, Y. (2011) Local control of *β*-adrenergic stimulation: effects on ventricular myocyte electrophysiology and Ca^2+^-transient. Journal of Molecular and Cellular Cardiology, 50, 863–871.

Herron, T. J., Lee, P. and Jalife, J. (2012) Optical imaging of voltage and calcium in cardiac cells & tissues. Circulation Research, 110, 609–623.

Hess, P., Lansman, J. B. and Tsien, R. W. (1984) Different modes of Ca channel gating behaviour favoured by dihydropyridine Ca agonists and antagonists. Nature, 311, 538–544.

Hess, P., Lansman, J. B. and Tsien, R. W. (1986) Calcium channel selectivity for divalent and monovalent cations. voltage and concentration dependence of single channel current in ventricular heart cells. The Journal of General Physiology, 88, 293–319.

Hess, P. and Tsien, R. W. (1984) Mechanism of ion permeation through calcium channels. Nature, 309, 453–456.

van der Heyden, M. A., Wijnhoven, T. J. and Opthof, T. (2005) Molecular aspects of adrenergic modulation of cardiac L-type Ca^2+^channels. Cardiovascular research, 65, 28–39.

Hilgemann, D. W. and Noble, D. (1987) Excitation-contraction coupling and extracellular calcium transients in rabbit atrium: reconstruction of basic cellular mechanisms. Proceedings of the Royal Society of London B: Biological Sciences, 230, 163–205.

Himeno, Y., Asakura, K., Cha, C. Y., Memida, H., Powell, T., Amano, A. and Noma, A. (2015) A human ventricular myocyte model with a refined representation of excitation-contraction coupling. Biophysical Journal, 109, 415–427.

Hinch, R., Greenstein, J. L., Tanskanen, A. J., Xu, L. and Winslow, R. L. (2004) A simplified local control model of calcium-induced calcium release in cardiac ventricular myocytes. Biophysical Journal, 87, 3723–3736.

Hirano, Y., Fozzard, H. A. and January, C. T. (1989) Characteristics of L-and T-type Ca^2+^currents in canine cardiac Purkinje cells. American Journal of Physiology-Heart and Circulatory Physiology, 256, H1478–H1492.

Hirano, Y. and Hiraoka, M. (2003) Ca^2+^entry-dependent inactivation of L-type Ca current: a novel formulation for cardiac action potential models. Biophysical journal, 84, 696–708.

Hodgkin, A. L. and Huxley, A. F. (1952) A quantitative description of membrane current and its application to conduction and excitation in nerve. The Journal of Physiology, 117, 500–544.

Hodgkin, A. L. and Katz, B. (1949) The effect of sodium ions on the electrical activity of the giant axon of the squid. The Journal of Physiology, 108, 37–77.

Hoefen, R., Reumann, M., Goldenberg, I., Moss, A. J., Jin, O., Gu, Y., McNitt, S., Zareba, W., Jons, C., Kanters, J. K. et al. (2012) In silico cardiac risk assessment in patients with long QT syndrome: type 1: clinical predictability of cardiac models. Journal of the American College of Cardiology, 60, 2182–2191.

Höfer, G. F., Hohenthanner, K., Baumgartner, W., Groschner, K., Klugbauer, N., Hofmann, F. and Romanin, C. (1997) Intracellular Ca^2+^inactivates L-type Ca^2+^channels with a Hill coefficient of ~1 and an inhibition constant of ~4 microM by reducing channel’s open probability. Biophysical Journal, 73, 1857–1865.

Hofmann, F., Flockerzi, V., Kahl, S. and Wegener, J. W. (2014) L-type Ca_V_1.2 calcium channels: from in vitro findings to in vivo function. Physiological Reviews, 94, 303–326.

Hopenfeld, B. (2006) Mechanism for action potential alternans: the interplay between L-type calcium current and transient outward current. Heart Rhythm, 3, 345–352.

Horn, R. and Vandenberg, C. A. (1984) Statistical properties of single sodium channels. Journal of General Physiology, 84, 505–534.

Hund, T. J., Decker, K. F., Kanter, E., Mohler, P. J., Boyden, P. A., Schuessler, R. B., Yamada, K. A. and Rudy, Y. (2008) Role of activated CaMKII in abnormal calcium homeostasis and I_Na_ remodeling after myocardial infarction: Insights from mathematical modeling. Journal of Molecular and Cellular Cardiology, 45, 420–428.

Hund, T. J. and Rudy, Y. (2004) Rate dependence and regulation of action potential and calcium transient in a canine cardiac ventricular cell model. Circulation, 110, 3168–3174.

Imredy, J. P. and Yue, D. T. (1994) Mechanism of Ca^2+^-sensitive inactivation of L-type Ca^2+^channels. Neuron, 12, 1301–1318.

Inada, S., Hancox, J. C., Zhang, H. and Boyett, M. R. (2009) One-dimensional mathematical model of the atrioventricular node including atrio-nodal, nodal, and nodal-his cells. Biophysical Journal, 97, 2117–2127.

Iyer, V., Mazhari, R. and Winslow, R. L. (2004) A computational model of the human left-ventricular epicardial myocyte. Biophysical Journal, 87, 1507–1525.

Jafri, M. S., Rice, J. J. and Winslow, R. L. (1998) Cardiac Ca^2+^dynamics: the roles of ryanodine receptor adaptation and sarcoplasmic reticulum load. Biophysical Journal, 74, 1149–1168.

Joyner, R. W., Kumar, R., Wilders, R., Jongsma, H. J., Verheijck, E. E., Golod, D. A., Van Ginneken, A., Wagner, M. B. and Goolsby, W. N. (1996) Modulating l-type calcium current affects discontinuous cardiac action potential conduction. Biophysical journal, 71, 237–245.

Käaaäb, S., Nuss, H. B., Chiamvimonvat, N., O’Rourke, B., Pak, P. H., Kass, D. A., Marban, E. and Tomaselli, G. F. (1996) Ionic mechanism of action potential prolongation in ventricular myocytes from dogs with pacing-induced heart failure. Circulation Research, 78, 262–273.

Kamp, T. J., Hu, H. and Marban, E. (2000) Voltage-dependent facilitation of cardiac L-type Ca channels expressed in HEK-293 cells requires *β*-subunit. American Journal of Physiology-Heart and Circulatory Physiology, 278, H126–H136.

Kass, R. S. and Sanguinetti, M. C. (1984) Inactivation of calcium channel current in the calf cardiac Purkinje fiber. Evidence for voltage-and calcium-mediated mechanisms. The Journal of General Physiology, 84, 705–726.

Katsube, Y., Yokoshiki, H., Nguyen, L., Yamamoto, M. and Sperelakis, N. (1998) L-type Ca^2+^currents in ventricular myocytes from neonatal and adult rats. Canadian Journal of Physiology and Pharmacology, 76, 873–881.

Kawano, S. and Hiraoka, M. (1991) Transient outward currents and action potential alterations in rabbit ventricular myocytes. Journal of Molecular and Cellular Cardiology, 23, 681–693.

Keener, J. P. and Sneyd, J. (1998) Mathematical Physiology I: Cellular Physiology, vol. 1. Springer.

Kernik, D. C., Morotti, S., Wu, H., Garg, P., Duff, H. J., Kurokawa, J., Jalife, J., Wu, J. C., Grandi, E. and Clancy, C. E. (2019) A computational model of induced pluripotent stem-cell derived cardiomyocytes incorporating experimental variability from multiple data sources. The Journal of Physiology, 597, 4533–4564.

Kettlewell, S., Saxena, P., Dempster, J., Colman, M. A., Myles, R. C., Smith, G. L. and Workman, A. J. (2019) Dynamic clamping human and rabbit atrial calcium current: narrowing *i_CaL_* window abolishes early afterdepolarizations. The Journal of Physiology, 597, 3619–3638.

Kim, J., Ghosh, S., Nunziato, D. A. and Pitt, G. S. (2004) Identification of the components controlling inactivation of voltage-gated Ca^2+^channels. Neuron, 41, 745–754.

Kim, J. G., Sung, D. J., Kim, H.-j., Park, S. W., Won, K. J., Kim, B., Shin, H. C., Kim, K.-S., Leem, C. H., Zhang, Y. H. et al. (2016) Impaired inactivation of L-Type Ca^2+^current as a potential mechanism for variable arrhythmogenic liability of HERG K^+^channel blocking drugs. PloS One, 11.

Ko, J. H., Park, W. S., Kim, S. J. and Earm, Y. E. (2006) Slowing of the inactivation of voltage-dependent sodium channels by staurosporine, the protein kinase C inhibitor, in rabbit atrial myocytes. European Journal of Pharmacology, 534, 48–54.

Kodama, I., Boyett, M. R., Nikmaram, M. R., Yamamoto, M., Honjo, H. and Niwa, R. (1999) Regional differences in effects of E-4031 within the sinoatrial node. American Journal of Physiology-Heart and Circulatory Physiology, 276, H793–H802.

Koivumäki, J. T., Korhonen, T. and Tavi, P. (2011) Impact of sarcoplasmic reticulum calcium release on calcium dynamics and action potential morphology in human atrial myocytes: a computational study. PLoS Computational Biology, 7, e1001067.

Kurata, Y., Hisatome, I., Imanishi, S. and Shibamoto, T. (2002) Dynamical description of sinoatrial node pacemaking: improved mathematical model for primary pacemaker cell. American Journal of Physiology-Heart and Circulatory Physiology, 283, H2074–H2101.

Le Grand, B., Hatem, S., Deroubaix, E., Couetil, J.-P. and Coraboeuf, E. (1991) Calcium current depression in isolated human atrial myocytes after cessation of chronic treatment with calcium antagonists. Circulation Research, 69, 292–300.

Le Grand, B. L., Hatem, S., Deroubaix, E., Couétil, J.-P. and Coraboeuf, E. (1994) Depressed transient outward and calcium currents in dilated human atria. Cardiovascular Research, 28, 548–556.

Lei, C., Clerx, M., Gavaghan, D. J., Polonchuk, L., Mirams, G. R. and Wang, K. (2019) Rapid characterization of hERG channel kinetics I: using an automated high-throughput system. Biophysical Journal, 117, 2438–2454.

Lei, C., Clerx, M., Whittaker, D. G., Gavaghan, D. J., de Boer, T. P. and Mirams, G. R. (2020a) Accounting for variability in ion current recordings using a mathematical model of artefacts in voltage-clamp experiments. Philosophical Transactions of the Royal Society A: Mathematical, Physical and Engineering Sciences, 378, 20190348.

Lei, C., Fabbri, A., Whittaker, D. G., Clerx, M., Windley, M. J., Hill, A. P., Mirams, G. R. and de Boer, T. P. (2020b) A nonlinear and time-dependent leak current in the presence of calcium fluoride patchclamp seal enhancer. Wellcome Open Research, 5, 152.

Li, G. R. and Nattel, S. (1997) Properties of human atrial I_Ca_ at physiological temperatures and relevance to action potential. American Journal of Physiology-Heart and Circulatory Physiology, 272, H227–H235.

Li, G.-R., Yang, B., Feng, J., Bosch, R. F., Carrier, M. and Nattel, S. (1999) Transmembrane I_Ca_ contributes to rate-dependent changes of action potentials in human ventricular myocytes. American Journal of Physiology-Heart and Circulatory Physiology, 276, H98–H106.

Li, L., Niederer, S. A., Idigo, W., Zhang, Y. H., Swietach, P., Casadei, B. and Smith, N. P. (2010) A mathematical model of the murine ventricular myocyte: a data-driven biophysically based approach applied to mice overexpressing the canine NCX isoform. American Journal of Physiology-Heart and Circulatory Physiology, 299, H1045–H1063.

Li, M., Kanda, Y., Ashihara, T., Sasano, T., Nakai, Y., Kodama, M., Hayashi, E., Sekino, Y., Furukawa, T. and Kurokawa, J. (2017) Overexpression of KCNJ2 in induced pluripotent stem cell-derived cardiomyocytes for the assessment of QT-prolonging drugs. Journal of Pharmacological Sciences, 134, 75–85.

Li, P. and Rudy, Y. (2011) A model of canine Purkinje cell electrophysiology and Ca^2+^cycling: rate dependence, triggered activity, and comparison to ventricular myocytes. Circulation Research, 109, 71–79.

Li, Z., Ridder, B. J., Han, X., Wu, W. W., Sheng, J., Tran, P. N., Wu, M., Randolph, A., Johnstone, R. H., Mirams, G. R. et al. (2019) Assessment of an *in silico* mechanistic model for proarrhythmia risk prediction under the CiPa initiative. Clinical Pharmacology & Therapeutics, 105, 466–475.

Lindblad, D. S., Murphey, C. R., Clark, J. W. and Giles, W. R. (1996) A model of the action potential and underlying membrane currents in a rabbit atrial cell. American Journal of Physiology-Heart and Circulatory Physiology, 271, H1666–H1696.

Linz, K. W. and Meyer, R. (1997) Modulation of L-type calcium current by internal potassium in guinea pig ventricular myocytes. Cardiovascular Research, 33, 110–122.

Liu, Y., Zeng, W., Delmar, M. and Jalife, J. (1993) Ionic mechanisms of electronic inhibition and concealed conduction in rabbit atrioventricular nodal myocytes. Circulation, 88, 1634–1646.

Livshitz, L. M. and Rudy, Y. (2007) Regulation of Ca^2+^and electrical alternans in cardiac myocytes: role of CAMKII and repolarizing currents. American Journal of Physiology-Heart and Circulatory Physiology, 292, H2854–H2866.

Lovell, N. H., Cloherty, S. L., Celler, B. G. and Dokos, S. (2004) A gradient model of cardiac pacemaker myocytes. Progress in Biophysics and Molecular Biology, 85, 301–323.

Luo, C.-H. and Rudy, Y. (1994) A dynamic model of the cardiac ventricular action potential II. Afterdepolarizations, triggered activity, and potentiation. Circulation Research, 74, 1097–1113.

Ma, J., Guo, L., Fiene, S. J., Anson, B. D., Thomson, J. A., Kamp, T. J., Kolaja, K. L., Swanson, B. J. and January, C. T. (2011) High purity human-induced pluripotent stem cell-derived cardiomyocytes: electrophysiological properties of action potentials and ionic currents. American Journal of Physiology-Heart and Circulatory Physiology, 301, H2006–H2017.

Magyar, J., Iost, N., Körtvély, A., Bányász, T., Virág, L., Szigligeti, P., Varró, A., Opincariu, M., Szécsi, J., Papp, J. G. et al. (2000) Effects of endothelin-1 on calcium and potassium currents in undiseased human ventricular myocytes. Pflügers Archiv-European Journal of Physiology, 441, 144–149.

Magyar, J., Szentandrássy, N., Bányász, T., Fülöp, L., Varró, A. and Nánási, P. P. (2002) Effects of thymol on calcium and potassium currents in canine and human ventricular cardiomyocytes. British Journal of Pharmacology, 136, 330–338.

Mahajan, A., Sato, D., Shiferaw, Y., Baher, A., Xie, L.-H., Peralta, R., Olcese, R., Garfinkel, A., Qu, Z. and Weiss, J. N. (2008a) Modifying L-type calcium current kinetics: consequences for cardiac excitation and arrhythmia dynamics. Biophysical journal, 94, 411–423.

Mahajan, A., Shiferaw, Y., Sato, D., Baher, A., Olcese, R., Xie, L.-H., Yang, M.-J., Chen, P.-S., Restrepo, J. G., Karma, A. et al. (2008b) A rabbit ventricular action potential model replicating cardiac dynamics at rapid heart rates. Biophysical Journal, 94, 392–410.

Maier, L. S. and Bers, D. M. (2007) Role of Ca^2+^/calmodulin-dependent protein kinase (CaMK) in excitation–contraction coupling in the heart. Cardiovascular Research, 73, 631–640.

Mangold, K. E., Wang, W., Johnson, E. K., Bhagavan, D., Moreno, J. D., Nerbonne, J. M. and Silva, J. R. (2021) Identification of structures for ion channel kinetic models. PLoS Computational Biology, 17, e1008932.

Matsuoka, S., Sarai, N., Kuratomi, S., Ono, K. and Noma, A. (2003) Role of individual ionic current systems in ventricular cells hypothesized by a model study. The Japanese Journal of Physiology, 53, 105–123.

McAllister, R. E., Noble, D. and Tsien, R. W. (1975) Reconstruction of the electrical activity of cardiac Purkinje fibres. The Journal of Physiology, 251, 1–59.

McDonald, T. F., Cavalié, A., Trautwein, W. and Pelzer, D. (1986) Voltage-dependent properties of macroscopic and elementary calcium channel currents in guinea pig ventricular myocytes. Pflügers Archiv-European Journal of Physiology, 406, 437–448.

McDonald, T. F., Pelzer, S., Trautwein, W. and Pelzer, D. J. (1994) Regulation and modulation of calcium channels in cardiac, skeletal, and smooth muscle cells. Physiological reviews, 74, 365–507.

Mesirca, P., Torrente, A. G. and Mangoni, M. E. (2014) T-type channels in the sino-atrial and atrioventricular pacemaker mechanism. Pflügers Archiv-European Journal of Physiology, 466, 791–799.

Mewes, T. and Ravens, U. (1994) L-type calcium currents of human myocytes from ventricle of non-failing and failing hearts and from atrium. Journal of Molecular and Cellular Cardiology, 26, 1307–1320.

Michailova, A., Saucerman, J., Belik, M. E. and McCulloch, A. D. (2005) Modeling regulation of cardiac K_ATP_ and L-type Ca^2+^currents by ATP, ADP, and Mg^2+^. Biophysical Journal, 88, 2234–2249.

Mirams, G. R., Davies, M. R., Cui, Y., Kohl, P. and Noble, D. (2012) Application of cardiac electrophysiology simulations to pro-arrhythmic safety testing. British journal of pharmacology, 167, 932–945.

Miyoshi, S., Mitamura, H., Fujikura, K., Fukuda, Y., Tanimoto, K., Hagiwara, Y., Ita, M. and Ogawa, S. (2003) A mathematical model of phase 2 reentry: role of L-type Ca current. American Journal of Physiology-Heart and Circulatory Physiology, 284, H1285–H1294.

Nakayama, T., Kurachi, Y., Noma, A. and Irisawa, H. (1984) Action potential and membrane currents of single pacemaker cells of the rabbit heart. Pflügers Archiv-European Journal of Physiology, 402, 248–257.

Niederer, S. A., Fink, M., Noble, D. and Smith, N. P. (2009) A meta-analysis of cardiac electrophysiology computational models. Experimental Physiology, 94, 486–495.

Nilius, B. (1986) Possible functional significance of a novel type of cardiac Ca channel. Biomedica Biochimica Acta, 45, K37–45.

Nilius, B., Hess, P., Lansman, J. B. and Tsien, R. W. (1985) A novel type of cardiac calcium channel in ventricular cells. Nature, 316, 443–446.

Nivala, M., de Lange, E., Rovetti, R. and Qu, Z. (2012) Computational modeling and numerical methods for spatiotemporal calcium cycling in ventricular myocytes. Frontiers in Physiology, 3, 114.

Noble, D., Garny, A. and Noble, P. J. (2012) How the Hodgkin-Huxley equations inspired the cardiac physiome project. The Journal of Physiology, 590, 2613–2628.

Noble, D., Noble, S. J., Bett, G. C., Earm, Y. E., Ho, W. K. and So, I. K. (1991) The role of sodium-calcium exchange during the cardiac action potential. Annals of the New York Academy of Sciences, 639, 334–353.

Noble, D. and Rudy, Y. (2001) Models of cardiac ventricular action potentials: iterative interaction between experiment and simulation. Philosophical Transactions of the Royal Society of London A: Mathematical, Physical and Engineering Sciences, 359, 1127–1142.

Noble, D., Varghese, A., Kohl, P. and Noble, P. (1998) Improved guinea-pig ventricular cell model incorporating a diadic space, I_Kr_ and I_Ks_, and length-and tension-dependent processes. The Canadian Journal of Cardiology, 14, 123–134.

Noma, A. and Shibasaki, T. (1985) Membrane current through adenosine-triphosphate-regulated potassium channels in guinea-pig ventricular cells. The Journal of Physiology, 363, 463–480.

Nordin, C. (1993) Computer model of membrane current and intracellular Ca^2+^flux in the isolated guinea pig ventricular myocyte. American Journal of Physiology-Heart and Circulatory Physiology, 265, H2117–H2136.

Nordin, C., Siri, F. and Aronson, R. S. (1989) Electrophysiologic characteristics of single myocytes isolated from hypertrophied guinea-pig hearts. Journal of Molecular and Cellular Cardiology, 21, 729–739.

Nygren, A., Fiset, C., Firek, L., Clark, J. W., Lindblad, D. S., Clark, R. B. and Giles, W. R. (1998) Mathematical model of an adult human atrial cell: the role of K^+^currents in repolarization. Circulation Research, 82, 63–81.

O’Hara, T., Virág, L., Varró, A. and Rudy, Y. (2011) Simulation of the undiseased human cardiac ventricular action potential: model formulation and experimental validation. PLoS Computational Biology, 7, e1002061.

O’Rourke, B., Backx, P. H. and Marban, E. (1992) Phosphorylation-independent modulation of L-type calcium channels by magnesium-nucleotide complexes. Science, 257, 245–248.

Ortner, N. J. and Striessnig, J. (2016) L-type calcium channels as drug targets in CNS disorders. Channels, 10, 7–13.

Paci, M., Hyttinen, J., Aalto-Setälä, K. and Severi, S. (2013) Computational models of ventricular- and atrial-like human induced pluripotent stem cell derived cardiomyocytes. Annals of Biomedical Engineering, 41, 2334–2348.

Pandit, S. V., Clark, R. B., Giles, W. R. and Demir, S. S. (2001) A mathematical model of action potential heterogeneity in adult rat left ventricular myocytes. Biophysical Journal, 81, 3029–3051.

Pásek, M., Simurda, J. and Christé, G. (2006) The functional role of cardiac T-tubules explored in a model of rat ventricular myocytes. Philosophical Transactions of the Royal Society A: Mathematical, Physical and Engineering Sciences, 364, 1187–1206.

Pathmanathan, P., Gray, R. A., Romero, V. J. and Morrison, T. M. (2017) Applicability analysis of validation evidence for biomedical computational models. Journal of Verification, Validation and Uncertainty Quantification, 2.

Pelzmann, B., Schaffer, P., Bernhart, E., Lang, P., Mächler, H., Rigler, B. and Koidl, B. (1998) L-type calcium current in human ventricular myocytes at a physiological temperature from children with tetralogy of Fallot. Cardiovascular Research, 38, 424–432.

Peterson, B. Z., DeMaria, C. D. and Yue, D. T. (1999) Calmodulin is the Ca^2+^sensor for Ca^2+^-dependent inactivation of L-type calcium channels. Neuron, 22, 549–558.

Pitt, G. S., Zühlke, R. D., Hudmon, A., Schulman, H., Reuter, H. and Tsien, R. W. (2001) Molecular basis of calmodulin tethering and Ca^2+^-dependent inactivation of L-type Ca^2+^channels. Journal of Biological Chemistry, 276, 30794–30802.

Pitzer, K. S. and Mayorga, G. (1973) Thermodynamics of electrolytes. II. Activity and osmotic coefficients for strong electrolytes with one or both ions univalent. The Journal of Physical Chemistry, 77, 2300–2308.

Pohl, A., Wachter, A., Hatam, N. and Leonhardt, S. (2016) A computational model of a human single sinoatrial node cell. Biomedical Physics & Engineering Express, 2, 035006.

Priebe, L. and Beuckelmann, D. J. (1998) Simulation study of cellular electric properties in heart failure. Circulation Research, 82, 1206–1223.

Puglisi, J. L., Yuan, W., Bassani, J. W. and Bers, D. M. (1999) Ca^2+^influx through Ca^2+^channels in rabbit ventricular myocytes during action potential clamp: influence of temperature. Circulation Research, 85, e7–e16.

Quinn, T. A., Granite, S., Allessie, M. A., Antzelevitch, C., Bollensdorff, C., Bub, G., Burton, R. A. B., Cerbai, E., Chen, P. S., Delmar, M. et al. (2011) Minimum Information about a Cardiac Electrophysiology Experiment (MICEE): Standardised reporting for model reproducibility, interoperability, and data sharing. Progress in Biophysics and Molecular Biology, 107, 4–10.

Ramasubramanian, S. and Rudy, Y. (2018) The structural basis of IKs ion-channel activation: Mechanistic insights from molecular simulations. Biophysical Journal, 114, 2584–2594.

Ramirez, R. J., Nattel, S. and Courtemanche, M. (2000) Mathematical analysis of canine atrial action potentials: rate, regional factors, and electrical remodeling. American Journal of Physiology-Heart and Circulatory Physiology, 279, H1767–H1785.

Rasmusson, R. L., Clark, J. W., Giles, W. R., Robinson, K., Clark, R. B., Shibata, E. F. and Campbell, D. L. (1990) A mathematical model of electrophysiological activity in a bullfrog atrial cell. American Journal of Physiology-Heart and Circulatory Physiology, 259, H370–H389.

Restrepo, J. G., Weiss, J. N. and Karma, A. (2008) Calsequestrin-mediated mechanism for cellular calcium transient alternans. Biophysical Journal, 95, 3767–3789.

Reuter, H. (1968) Slow inactivation of currents in cardiac Purkinje fibres. The Journal of Physiology, 197, 233–253.

Reuter, H. (1973) Divalent cations as charge carriers in excitable membranes. Progress in Biophysics and Molecular Biology, 26, 1–43.

Reuter, H. (1974) Localization of *beta* adrenergic receptors, and effects of noradrenaline and cyclic nucleotides on action potentials, ionic currents and tension in mammalian cardiac muscle. The Journal of Physiology, 242, 429–451.

Rice, J. J., Jafri, M. S. and Winslow, R. L. (1999) Modeling gain and gradedness of Ca^2+^release in the functional unit of the cardiac diadic space. Biophysical Journal, 77, 1871–1884.

Rohr, S. and Kucera, J. P. (1997) Involvement of the calcium inward current in cardiac impulse propagation: induction of unidirectional conduction block by nifedipine and reversal by bay k 8644. Biophysical journal, 72, 754–766.

Rose, W. C., Balke, C. W., Wier, W. G. and Marban, E. (1992) Macroscopic and unitary properties of physiological ion flux through L-type Ca^2+^channels in guinea-pig heart cells. The Journal of Physiology, 456, 267–284.

Roux, B., Allen, T., Berneche, S. and Im, W. (2004) Theoretical and computational models of biological ion channels. Quarterly Reviews of Biophysics, 37, 15–103.

Rovetti, R., Cui, X., Garfinkel, A., Weiss, J. N. and Qu, Z. (2010) Spark-induced sparks as a mechanism of intracellular calcium alternans in cardiac myocytes. Circulation Research, 106, 1582.

Rubart, M., Lopshire, J. C., Fineberg, N. S. and Zipes, D. P. (2000) Changes in left ventricular repolarization and ion channel currents following a transient rate increase superimposed on bradycardia in anesthetized dogs. Journal of Cardiovascular Electrophysiology, 11, 652–664.

Rudy, Y. and Silva, J. R. (2006) Computational biology in the study of cardiac ion channels and cell electrophysiology. Quarterly Reviews of Biophysics, 39, 57–116.

Sanguinetti, M. C., Jiang, C., Curran, M. E. and Keating, M. T. (1995) A mechanistic link between an inherited and an acquird cardiac arrthytmia: HERG encodes the I_Kr_ potassium channel. Cell, 81, 299–307.

Sato, D., Shiferaw, Y., Garfinkel, A., Weiss, J. N., Qu, Z. and Karma, A. (2006) Spatially discordant alternans in cardiac tissue: role of calcium cycling. Circulation Research, 99, 520–527.

Satoh, H. (1994) Elevation of intracellular Ca^2+^concentration by protein kinase C stimulation in isolated single rabbit sino-atrial node cells. General Pharmacology: The Vascular System, 25, 325–332.

Schulz, D. J., Goaillard, J.-M. and Marder, E. (2006) Variable channel expression in identified single and electrically coupled neurons in different animals. Nature Neuroscience, 9, 356–362.

Scriven, D. R., Dan, P. and Moore, E. D. (2000) Distribution of proteins implicated in excitation-contraction coupling in rat ventricular myocytes. Biophysical journal, 79, 2682–2691.

Shannon, T. R., Wang, F. and Bers, D. M. (2005) Regulation of cardiac sarcoplasmic reticulum Ca release by luminal [Ca] and altered gating assessed with a mathematical model. Biophysical Journal, 89, 4096–4110.

Shannon, T. R., Wang, F., Puglisi, J., Weber, C. and Bers, D. M. (2004) A mathematical treatment of integrated Ca dynamics within the ventricular myocyte. Biophysical Journal, 87, 3351–3371.

Shaw, R. M. and Rudy, Y. (1997) Ionic mechanisms of propagation in cardiac tissue: roles of the sodium and l-type calcium currents during reduced excitability and decreased gap junction coupling. Circulation research, 81, 727–741.

Shirokov, R., Levis, R., Shirokova, N. and Ríos, E. (1993) Ca^2+^-dependent inactivation of cardiac L-type Ca^2+^channels does not affect their voltage sensor. The Journal of General Physiology, 102, 1005–1030.

Silva, J. R. (2018) How to connect cardiac excitation to the atomic interactions of ion channels. Biophysical Journal, 114, 259–266.

Smith, G. D. (1996) Analytical steady-state solution to the rapid buffering approximation near an open Ca^2+^channel. Biophysical Journal, 71, 3064–3072.

Soldatov, N. M. (2003) Ca^2+^channel moving tail: link between Ca^2+^-induced inactivation and Ca^2+^signal transduction. TRENDS in pharmacological Sciences, 24, 167–171.

Soltis, A. R. and Saucerman, J. J. (2010) Synergy between CaMKII substrates and *β*-adrenergic signaling in regulation of cardiac myocyte Ca^2+^handling. Biophysical Journal, 99, 2038–2047.

Stern, M. D. (1992) Theory of excitation-contraction coupling in cardiac muscle. Biophysical Journal, 63, 497–517.

Striessnig, J., Pinggera, A., Kaur, G., Bock, G. and Tuluc, P. (2014) L-type Ca^2+^channels in heart and brain. Wiley Interdisciplinary Reviews: Membrane Transport and Signaling, 3, 15–38.

Sun, H., Leblanc, N. and Nattel, S. (1997) Mechanisms of inactivation of L-type calcium channels in human atrial myocytes. American Journal of Physiology-Heart and Circulatory Physiology, 272, H1625–H1635.

Sun, L., Fan, J.-S., Clark, J. W. and Palade, P. T. (2000) A model of the L-type Ca^2+^channel in rat ventricular myocytes: ion selectivity and inactivation mechanisms. The Journal of Physiology, 529, 139–158.

Szabó, G., Szentandrássy, N., Bíró, T., Tóth, B. I., Czifra, G., Magyar, J., Bányász, T., Varró, A., Kovács, L. and Nánási, P. P. (2005) Asymmetrical distribution of ion channels in canine and human left-ventricular wall: epicardium versus midmyocardium. Pflügers Archiv-European Journal of Physiology, 450, 307–316.

Tadross, M. R., Dick, I. E. and Yue, D. T. (2008) Mechanism of local and global Ca^2+^sensing by calmodulin in complex with a Ca^2+^channel. Cell, 133, 1228–1240.

Takagi, S., Kihara, Y., Sasayama, S. and Mitsuiye, T. (1998) Slow inactivation of cardiac L-type Ca^2+^channel induced by cold acclimation of guinea pig. American Journal of Physiology-Regulatory, Integrative and Comparative Physiology, 274, R348–R356.

Taniguchi, J., Noma, A. and Irisawa, H. (1983) Modification of the cardiac action potential by intracellular injection of adenosine triphosphate and related substances in guinea pig single ventricular cells. Circulation Research, 53, 131–139.

Ten Tusscher, K. H. W. J., Noble, D., Noble, P.-J. and Panfilov, A. V. (2004) A model for human ventricular tissue. American Journal of Physiology-Heart and Circulatory Physiology, 286, H1573–H1589.

Ten Tusscher, K. H. W. J. and Panfilov, A. V. (2006) Alternans and spiral breakup in a human ventricular tissue model. American Journal of Physiology-Heart and Circulatory Physiology, 291, H1088–H1100.

Tomek, J., Bueno-Orovio, A., Passini, E., Zhou, X., Minchole, A., Britton, O., Bartolucci, C., Severi, S., Shrier, A., Virag, L. et al. (2019) Development, calibration, and validation of a novel human ventricular myocyte model in health, disease, and drug block. eLife, 8.

Trovato, C., Passini, E., Nagy, N., Varró, A., Abi-Gerges, N., Severi, S. and Rodriguez, B. (2020) Human Purkinje in silico model enables mechanistic investigations into automaticity and pro-arrhythmic abnormalities. Journal of Molecular and Cellular Cardiology.

Tseng, G.-N. (1988) Calcium current restitution in mammalian ventricular myocytes is modulated by intracellular calcium. Circulation Research, 63, 468–482.

Tseng, G. N., Robinson, R. B. and Hoffman, B. F. (1987) Passive properties and membrane currents of canine ventricular myocytes. The Journal of General Physiology, 90, 671–701.

Van Wagoner, D. R., Pond, A. L., Lamorgese, M., Rossie, S. S., McCarthy, P. M. and Nerbonne, J. M. (1999) Atrial L-type Ca^2+^currents and human atrial fibrillation. Circulation Research, 85, 428–436.

Varela, M., Colman, M. A., Hancox, J. C. and Aslanidi, O. V. (2016) Atrial heterogeneity generates re-entrant substrate during atrial fibrillation and anti-arrhythmic drug action: mechanistic insights from canine atrial models. PLoS Computational Biology, 12, e1005245.

Veerman, C. C., Mengarelli, I., Guan, K., Stauske, M., Barc, J., Tan, H. L., Wilde, A. A. M., Verkerk, A. O. and Bezzina, C. R. (2016) hiPSC-derived cardiomyocytes from Brugada Syndrome patients without identified mutations do not exhibit clear cellular electrophysiological abnormalities. Scientific Reports, 6, 1–10.

Vitek, M. and Trautwein, W. (1971) Slow inward current and action potential in cardiac Purkinje fibres. Pfliígers Archiv-European Journal of Physiology, 323, 204–218.

Vornanen, M. and Shepherd, N. (1997) Restitution of contractility in single ventricular myocytes of guinea pig heart. Cardiovascular Research, 33, 611–622.

Weiss, J. N., Garfinkel, A., Karagueuzian, H. S., Chen, P.-S. and Qu, Z. (2010) Early afterdepolarizations and cardiac arrhythmias. Heart Rhythm, 7, 1891–1899.

Weiss, J. N., Garfinkel, A., Karagueuzian, H. S., Nguyen, T. P., Olcese, R., Chen, P.-S. and Qu, Z. (2015) Perspective: a dynamics-based classification of ventricular arrhythmias. Journal of molecular and cellular cardiology, 82, 136–152.

Whittaker, D. G., Clerx, M., Lei, C. L., Christini, D. J. and Mirams, G. R. (2020) Calibration of ionic and cellular cardiac electrophysiology models. Wiley Interdisciplinary Reviews: Systems Biology and Medicine, e1482.

Wilders, R., Jongsma, H. J. and Van Ginneken, A. C. G. (1991) Pacemaker activity of the rabbit sinoatrial node: A comparison of mathematical models. Biophysical Journal, 60, 1202–1216.

Winslow, R. L., Rice, J., Jafri, S., Marban, E. and O’Rourke, B. (1999) Mechanisms of altered excitation-contraction coupling in canine tachycardia-induced heart failure, II: model studies. Circulation Research, 84, 571–586.

Winslow, R. L., Walker, M. A. and Greenstein, J. L. (2016) Modeling calcium regulation of contraction, energetics, signaling, and transcription in the cardiac myocyte. Wiley Interdisciplinary Reviews: Systems Biology and Medicine, 8, 37–67.

You, Y., Pelzer, D. J. and Pelzer, S. (1995) Trypsin and forskolin decrease the sensitivity of L-type calcium current to inhibition by cytoplasmic free calcium in guinea pig heart muscle cells. Biophysical journal, 69, 1838–1846.

You, Y., Pelzer, D. J. and Pelzer, S. (1997) Modulation of L-type Ca^2+^current by fast and slow Ca^2+^buffering in guinea pig ventricular cardiomyocytes. Biophysical Journal, 72, 175–187.

Yu, T., Lloyd, C. M., Nickerson, D. P., Cooling, M. T., Miller, A. K., Garny, A., Terkildsen, J. R., Lawson, J., Britten, R. D., Hunter, P. J. et al. (2011) The Physiome model repository 2. Bioinformatics, 27, 743–744.

Yue, L., Feng, J., Gaspo, R., Li, G.-R., Wang, Z. and Nattel, S. (1997) Ionic remodeling underlying action potential changes in a canine model of atrial fibrillation. Circulation Research, 81, 512–525.

Zahradníková, A., Kubalová, Z., Pavelková, J., Gyorke, S. and Zahradník, I. (2004) Activation of calcium release assessed by calcium release-induced inactivation of calcium current in rat cardiac myocytes. American Journal of Physiology-Cell Physiology, 286, C330–C341.

Zamponi, G. W., Striessnig, J., Koschak, A. and Dolphin, A. C. (2015) The physiology, pathology, and pharmacology of voltage-gated calcium channels and their future therapeutic potential. Pharmacological Reviews, 67, 821–870.

Zeng, J., Laurita, K. R., Rosenbaum, D. S. and Rudy, Y. (1995) Two components of the delayed rectifier K^+^current in ventricular myocytes of the guinea pig type: Theoretical formulation and their role in repolarization. Circulation Research, 77, 140–152.

Zhang, H., Holden, A. V., Kodama, I., Honjo, H., Lei, M., Varghese, T. and Boyett, M. R. (2000) Mathematical models of action potentials in the periphery and center of the rabbit sinoatrial node. American Journal of Physiology-Heart and Circulatory Physiology, 279, H397–H421.

Zhou, Z. and Bers, D. M. (2000) Ca^2+^influx via the L-type Ca^2+^channel during tail current and above current reversal potential in ferret ventricular myocytes. The Journal of Physiology, 523, 57–66.

Zhu, Z. I. and Clancy, C. E. (2007) L-type Ca^2+^channel mutations and T-wave alternans: a model study. American Journal of Physiology-Heart and Circulatory Physiology, 293, H3480–H3489.

